# Evaluating genome assemblies with HMM-Flagger

**DOI:** 10.64898/2026.02.27.708355

**Authors:** Mobin Asri, Jordan M. Eizenga, Prajna Hebbar, Taylor D. Real, Julian Lucas, Hailey Loucks, Alessandro Calicchio, Mark Diekhans, Evan E. Eichler, Sofie Salama, Karen H. Miga, Benedict Paten

## Abstract

HMM-Flagger is a reference-free tool for detecting structural errors in haplotype-resolved genome assemblies based upon the coverage of mapped reads. It models read coverage with a hidden Markov model augmented by a Gaussian autoregressive process, which enables classifying coverage anomalies as erroneous blocks, false duplications, or collapsed blocks. Trained and tested on synthetic misassemblies, it detected synthetic errors using Pacific Biosciences HiFi and Oxford Nanopore Technologies R10 data with F1 scores of 78.4% and 60.4% respectively. When applied to six HG002 assemblies it revealed multiple large misassemblies including false duplications and collapse events in human satellites. Applied to assemblies from the Human Pangenome Reference Consortium (HPRC), HMM-Flagger demonstrated substantial improvements from release 1 (0.94% error rate) to release 2 (0.38%), reflecting technological advances. HMM-Flagger also validated NOTCH2NL assemblies in HPRC release 2 and confirmed the correctness of three novel structural configurations.

## 1 Introduction

Genome assemblies represent long DNA sequences of an individual genome[1]. Recent improvements in long-read sequencing technologies from Oxford Nanopore Technologies (ONT) and Pacific Biosciences (PacBio) have enabled a new generation of genome assemblers that produce nearly telomere-to-telomere and haplotype-resolved assemblies[2–4]. This has paved the way for large-scale initiatives such as the Human Pangenome Reference Consortium (HPRC) and the Human Genome Structural Variation Consortium (HGSVC) [5, 6]. However, even with these contemporary sequencing technologies, highly repetitive or segmentally duplicated regions still remain difficult to assemble [7, 8]. To validate such assemblies at scale, automated methods are needed to detect assembly errors.

Most of the current error detection methods are based on either mapping to a high-quality reference (so called reference-based methods) or decomposing the assembly into a set of shorter sequences (k-mers). A drawback of reference-based methods is conflating real genomic variations with assembly errors. This problem disappears when the reference comes from the same sample, which is the basis for tools like GQC[1].

However, in most cases such a truth assembly is unavailable, and to date, well-validated reference assemblies exist for only a few samples for which tremendous effort has been expended, like T2T-CHM13 (haploid European sample), T2T-Q100-HG002 (Ashkenazi), T2T-YAO (Chinese) and I002C (South Asian)[1, 9–11]. To eliminate the need for a reference, k-mer-based methods like Merqury[12] and Yak[13] extract sets of k-mers from sequencing reads and assembly and then find inconsistencies between the two sets. While useful for detecting phasing switch errors and estimating the number of base-level errors (normally expressed as a Phred scaled Quality Value (QV)), their reliance on short k-mers severely limits their performance in repetitive regions.

An alternative way to evaluate assemblies is by mapping reads back to them and identifying abnormal mapping patterns, asking in effect, how likely is the assembly given the reads? Some of the early tools with this approach include CGAL [14] and Asset [15]. CGAL, developed for short reads, computes a single probabilistic score without pinpointing misassembled regions. Asset integrates long-read data to report coverage anomalies but relies on fixed thresholds, which limits its robustness under varying sequencing depths.

More recent tools, such as NucFlag [1] and Flagger [5], have improved this approach for long-read data. NucFlag adjusts its thresholds based on sequencing coverage and considers secondary alleles (differences in the reads that indicate the reference might be wrong or contain a copy-number error) to improve detection accuracy. Flagger, developed by the Human Pangenome Reference Consortium (HPRC), fits a Gaussian Mixture Model (GMM) to the coverage distributions in 5 Mb windows to detect three types of misassemblies: collapsed blocks (where more reads map than expected, indicating the assembly has too few copies), false duplications (the reverse), and erroneous blocks (where so few reads map that the assembly is unlikely to be correct).

While Flagger played a key role in evaluating HPRC assemblies, its coarse 5 Mb resolution limits the detection of short errors. In addition, its strategy of treating each window independently prevents the propagation of coverage information between adjacent regions. Building on Flagger, we have developed HMM-Flagger which addresses these issues by using a hidden Markov model (HMM). Using either HiFi or ONT mappings, HMM-Flagger accurately identifies synthetic misassemblies in HG002-T2T-v1.1 and quantifies improvements in the assembly methods employed by the HPRC. Of particular value is HMM-Flagger’s ability to identify errors in hard-to-assemble regions, here we show that it can detect large collapses and duplications in repetitive regions such as centromeric higher-order repeats (HORs) and also complex loci like NOTCH2NL genes.

## 2 Results

### 2.1 HMM-Flagger Overview

HMM-Flagger is a read-mapping-based tool developed for assessing haplotype-resolved assemblies. It takes long read mappings of the same sample to the assembly (in BAM/SAM format), computes coverage across the genome, averages coverage within fixed windows, and finally uses an HMM to find anomalies in coverage and assigns them to different types of misassemblies (**Figs 1A and 1B**). The window size was tuned separately for HiFi (16kb), ONT-R10 (16kb) and ONT-R9 (8kb) platforms (**Supp Note 1**). HMM-Flagger outputs a BED file that partitions the genome into four categories; Erroneous (Err), Duplicated (Dup), Haploid (Hap), Collapsed (Col), where all except Hap indicate potential misassemblies. Each category corresponds to a distinct state in the HMM with emission densities selected based on our prior knowledge of coverage distributions in misassembled regions. The HMM transition probabilities are constrained by the mapping quality (MAPQ) of the reads (**Methods**). For example, a transition to the Dup state is allowed only when the fraction of low-MAPQ reads exceeds a threshold, since false duplications are typically accompanied by ambiguous mappings. Rather than relying on fixed values, the emission and transition parameters are estimated using the Expectation Maximization (EM) algorithm (**Fig 1C**), making HMM-Flagger robust across sequencing depths and platforms.

**Fig. 1.**
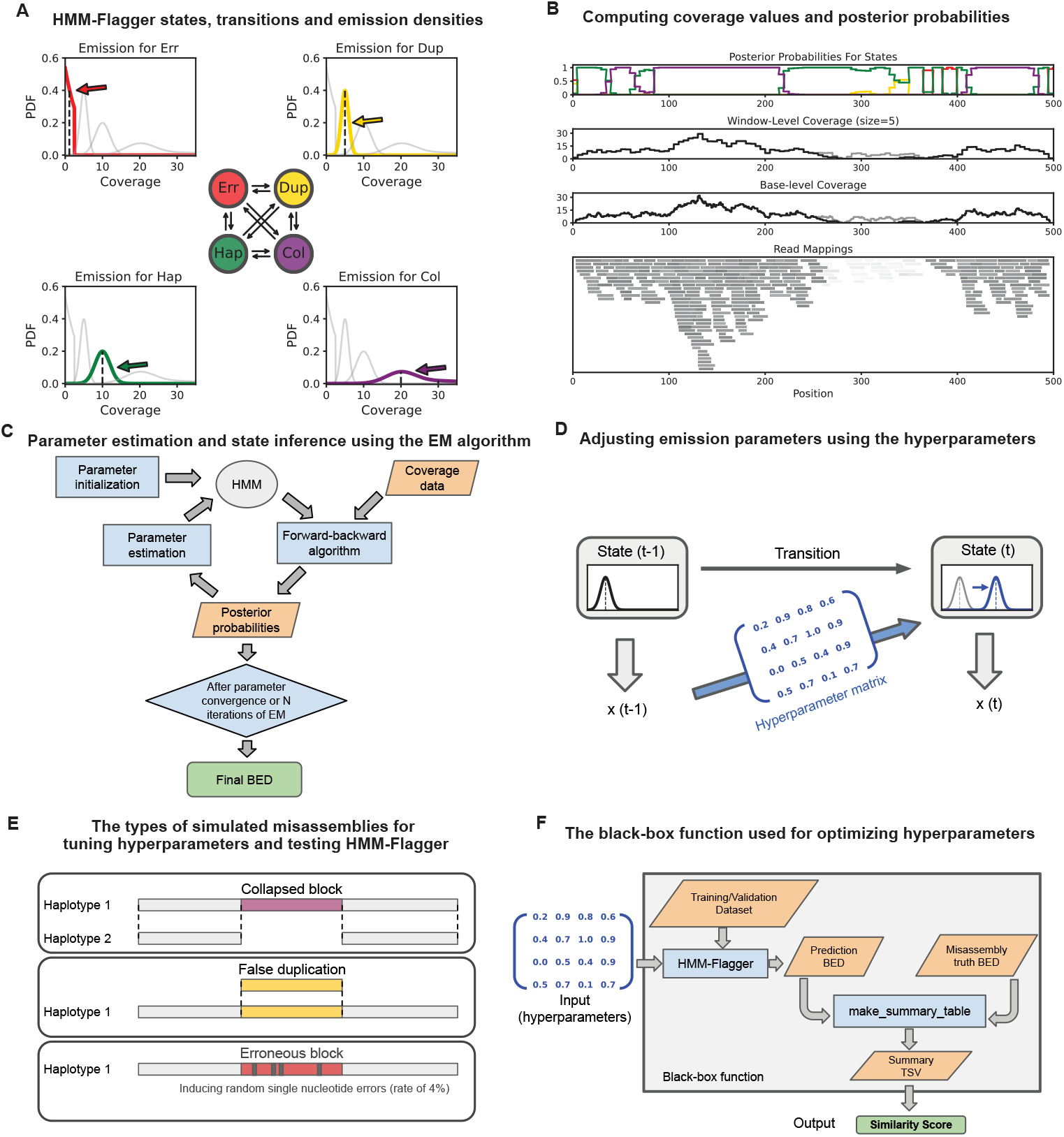
Overview of HMM-Flagger’s model, inputs/outputs, and hyperparameter tuning with simulated misassemblies. **A**) The states of HMM-Flagger are shown with colored circles in the center and the possible transitions between them are shown with black arrows. The emission density for each state is drawn in a separate panel with the corresponding color alongside the densities for the other states in gray for comparison. **B**) The bottom panel shows a toy example of read mappings given as input to HMM-Flagger. The reads with lower mapping quality are shown lighter. In the two middle panels for base-level and window-level coverages, the gray line shows the coverage of all reads and the black line shows the coverage of the reads with high mapping quality (MAPQ*>*10). The posterior probabilities for different states are shown in the top panel with colors matching (**A**). **C**) The workflow for estimating HMM parameters, computing posterior probabilities and creating the final BED file. **D**) Schematic of the hyperparameter matrix, which shifts the emission parameters for position *t* based on the previous observation (*x*_*t*−1_). **E**) The three types of misassemblies created by Falsifier for tuning hyperparameters and testing HMM-Flagger. **F**) Schematic of the black-box function used for tuning the hyperparameter matrix with a Bayesian optimization approach. The function consists of taking hyperparameters as input, running HMM-Flagger and outputting truth-prediction similarity in a single score.

An assumption in the HMM explained above is that given the states of two consecutive windows, their observed coverages are independent. However in reality this assumption is not satisfied since same reads might span consecutive windows therefore their coverages are not truly independent. To account for local dependence in coverage expectations we augmented the HMM with a Gaussian AutoRegressive Process (GARP), where emission parameters are determined using a linear combination of the previous observation and a constant term (**Fig 1D**)[16]. To optimize the factors of the linear combinations for GARP, represented as a hyperparameter matrix, we introduced synthetic misassemblies into the HG002 Telomere-to-Telomere (T2T) reference (v1.1) [1]. These misassemblies included deletions, insertions, and single-base errors (**Fig 1E**). Then we mapped reads from different platforms to the falsified assemblies, compared the HMM-Flagger predictions to the truth misassembly coordinates and derived a similarity score (**Methods**). We then applied a Bayesian optimization approach [17] to fit the GARP parameters, treating their relationship to the similarity score as a black box function (**Fig 1F**) (**Methods**).

To further improve the precision of HMM-Flagger we employed three strategies. First, we corrected for coverage drops at contig ends, caused by mappers discarding short alignments. To account for this we derived a factor that adjusts coverage expectations based on the minimum fraction of each read that must map to be retained. This factor is tuned separately for HiFi, ONT-R10, and ONT-R9 platforms (**Supp Fig 1**). Second, to account for platform-specific biases, particularly in pericentromeric satellites (**Supp Fig 2**), HMM-Flagger accepts BED files for such regions and estimates the HMM parameters for each region independently. Finally, we developed a filtering step that uses self-homology mappings of assembly contigs to identify sequences missing in one haplotype (confirming Col predictions) or redundantly represented multiple times (confirming Dup predictions), resulting in a conservative prediction set (**Methods**).

### 2.2 Benchmarking on synthetic misassemblies

To benchmark HMM-Flagger and compare it against other evaluation tools we created a set of synthetic misassemblies independent of those used for hyperparameter tuning. We developed a program called Falsifier to create two falsified assemblies with 0.87% and 3.32% misassembly rates, which fall within the range of unreliability rates observed in prior HPRC assemblies. Reads from three sequencing platforms; PacBio-HiFi, ONT-R10 and ONT-R9 were mapped to each falsified assembly, producing 6 BAM files. Next we ran HMM-Flagger, Flagger-v0.4.0 (the previous non-HMM version of Flagger) [5] and NucFlag-v0.3.6[18] on each BAM file and measured the accuracy of predictions (**Methods**). NucFlag was not benchmarked on ONT data since it was not optimized for this platform. Ignoring tool-specific labels we evaluated performance by counting the number of bases correctly identified as problematic (true positives), the misassembled bases that were missed (false negatives), and the correctly assembled bases flagged by mistake (false positives).

Using the assembly with the higher misassembly rate (3.32%) HMM-Flagger predictions (unfiltered) achieved the highest recall across all platforms with the rates of 82.8%, 72.3% and 69.3% for HiFi, ONT-R10 and ONT-R9 respectively (**Figs 2A and 2B**; **Supp Fig 3**; **Supp Table 1**). Meanwhile, the conservative set of HMM-Flagger achieved the highest precision. Based on F1 scores HMM-Flagger (78.4%) outperformed non-HMM Flagger (58.9%) and NucFlag (57.5%) using HiFi data. The same patterns were observed with the lower misassembly rate (0.87%) (**Supp Fig 4; Supp Table 2**). The unfiltered and conservative sets of HMM-Flagger performed comparably based on F1 score, with HiFi favoring the unfiltered set and ONT favoring the conservative set. Furthermore, we down-sampled reads from 40× to 20× and computed F1 scores. Using HiFi reads for the unfiltered set the score dropped only marginally from 78.4% to 75.1% showing that HMM-Flagger is highly robust to coverage (**Supp Figs 5 and 6; Supp Tables 3 and 4**).

**Fig. 2.**
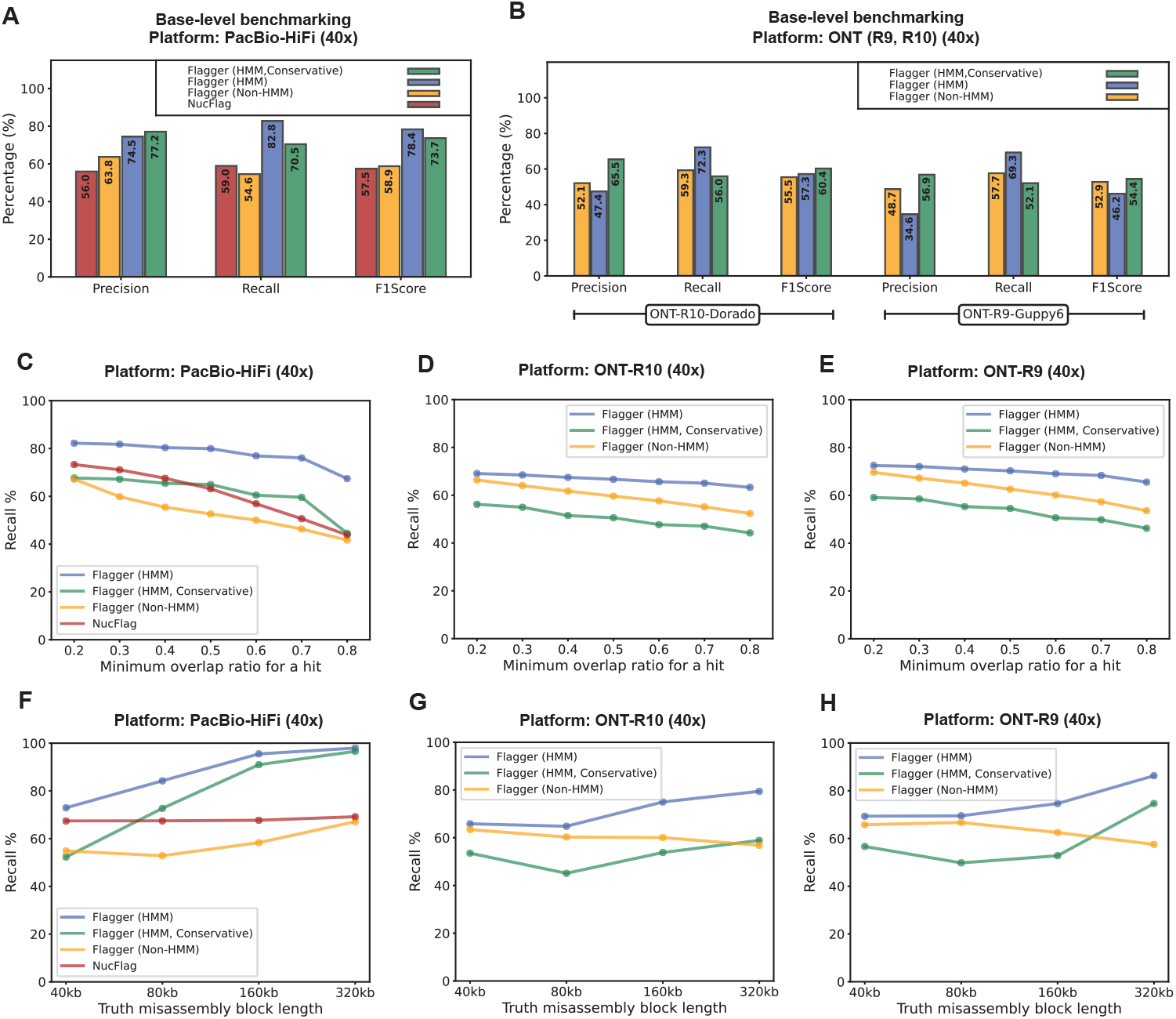
Benchmarking HMM-Flagger on synthetic misassemblies. **A, B)** Benchmarking on the falsified assembly with a misassembly rate of 3.32% using PacBio-HiFi, ONT-R9, and ONT-R10 reads. Metrics were computed by counting the number of TP (flagged misassembly), FP (correctly assembled but flagged), and FN (misassembly but not flagged) bases. **C, D, E)** Completeness of flagged misassembled blocks. The same assembly and reads were used as in (A) and (B). The recall rates (y-axis) were measured by counting the number of misassembled blocks covered by different minimum overlap ratios (x-axis). **F, G, H)** Impact of misassembly size on performance. The same assembly and reads were used as in (A) and (B). Recall rates were measured by counting misassembled blocks of each size that were covered at least 40% by flagged regions. The four misassembly sizes shown on the x-axis correspond to those used to generate the synthetic misassemblies.

Base-level benchmarking provides a coarse measure of accuracy, capturing only how many bases were flagged correctly. However, flagging only a part of a larger misassembled block can still provide a useful signal for further investigation. To account for this, we used an overlap-based metric that counts the number of misassembled blocks that were flagged by more than a minimum overlap threshold. Overlap-based recall rates were then computed by sweeping this threshold from 0.2 to 0.8. Using HiFi data, HMM-Flagger’s curve was above those of other tools, ranging from 82.24% at 0.2 to 68.34% at 0.8 (**Fig 2C; Supp Fig 7; Supp Tables 5 and 6**). A similar pattern was observed for ONT data (**Figs 2D and 2E; Supp Fig 8; Supp Tables 7-10**). Overlap-based precision, counting partially correct prediction blocks, also showed the better performance of HMM-Flagger compared to other tools. The conservative set was consistently more precise than the unfiltered one across different thresholds especially using ONT data (**Supp Fig 9**).

To measure the impact of misassembly size on performance we stratified the overlap-based recall rates by misassembly size, ranging from 40Kb to 320Kb, with the minimum overlap threshold set to 0.4. The results showed that in general recall rate increases with misassembly size (**Figs 2F, 2G, and 2H; Supp Tables 11-16**). For example, out of 146 misassemblies of 320 Kb, HMM-Flagger predictions covered 143 (96.58%) with an overlap ratio above 0.4 using HiFi. However for the smallest induced misassemblies (40Kb long) the recall rate was lower (72.92%) and stratifying by misassembly type revealed that false duplication was the hardest category to detect (**Supp Fig 10 and 11**).

### 2.3 Testing HMM-Flagger on six HG002 assemblies

HMM-Flagger was also tested on actual errors made by assemblers. For this purpose, we selected six recently published assemblies of the HG002 sample [1, 19–22], mapped HiFi and ONT-R10 reads to each one and used the four methods mentioned previously to detect misassemblies (**Supp Note 2**). For each combination of assembly, read platform and detection tool, we computed unreliability percentage, which is the number of flagged bases divided by the total size of the assembly.

With HiFi mappings, unreliability percentages ranged from 1.37% to 0.03% (rDNA array and gaps are ignored). We observed the highest unreliability percentage in the first scaffolded HG002 assembly that was created by the HPRC (HPRC-bakeoff) using Hifiasm (v0.14.1), PacBio HiFi reads and parental short reads[22]. The assembly with the lowest unreliability percentage was created by the Telomere-to-Telomere (T2T) Consortium using a wide range of sequencing platforms and by leveraging both automated pipelines and manual investigations (**Fig 3A**)[1]. A noticeable outlier was the output of NucFlag on the assembly created with the PECAT assembler and ONT-Duplex data. Because this assembly was built exclusively from ONT reads but evaluated using higher-accuracy HiFi mappings, the elevated unreliability likely reflects base-level discrepancies rather than structural errors. Most of the regions flagged in this assembly are structurally supported by both ONT and HiFi mappings (**Supp Fig 12**).

**Fig. 3.**
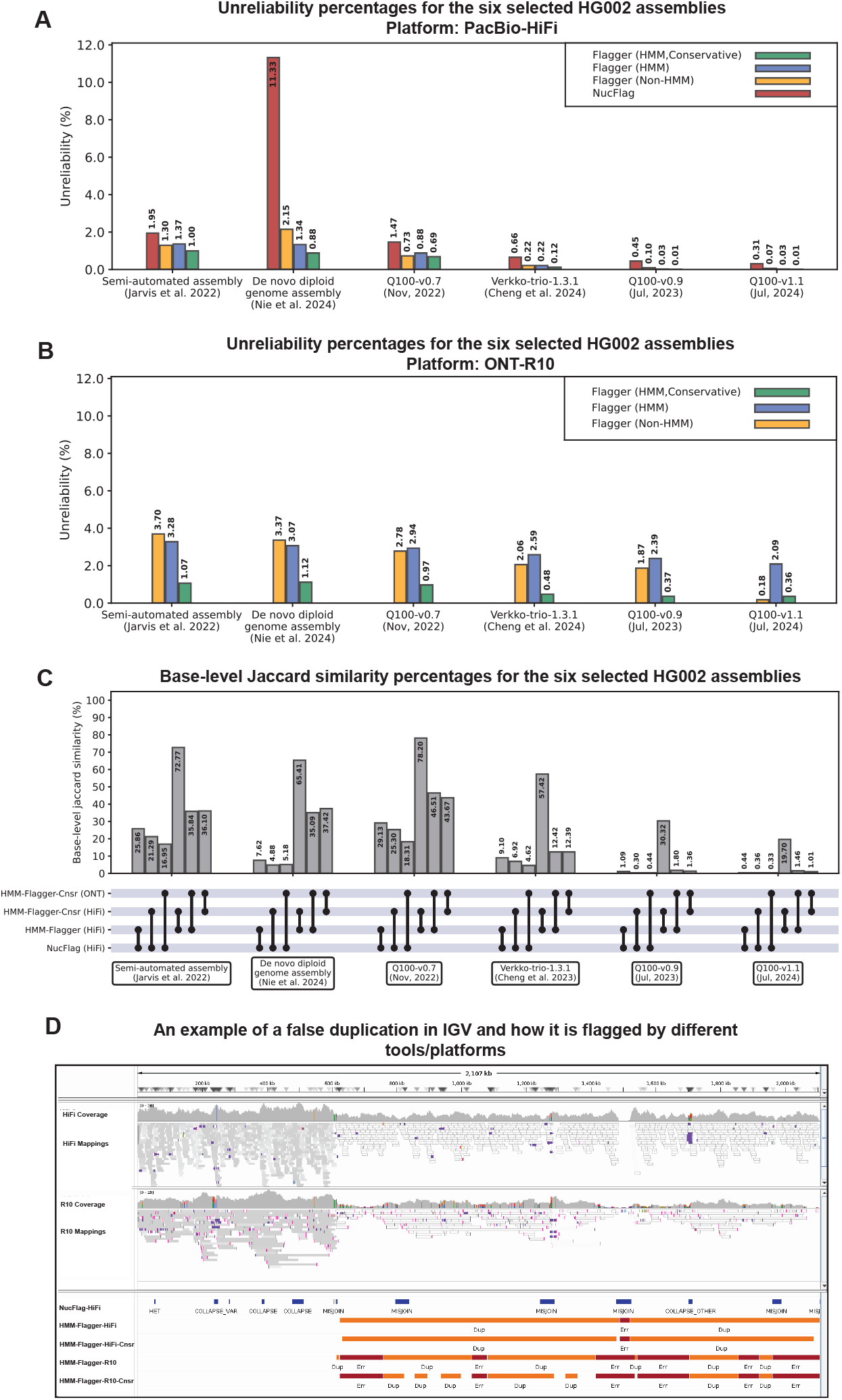
Testing HMM-Flagger on the six selected HG002 assemblies. **A)** Unreliability percentages for the six selected HG002 assemblies, computed using HiFi mappings. The rDNA arrays and gaps were ignored in unreliability calculation. The y axis shows unreliability percentage, which is calculated by dividing the number of flagged bases by the total size of the assembly. **B)** Same as (A) but computed using ONT-R10 mappings. The NucFlag results are not shown in this figure since it is not designed for ONT data. **C)** Computing base-level jaccard similarity percentages between different pairs of methods for the six selected HG002 assemblies. The dots at the end of each vertical black line show which methods are being compared. **D)** A 1.5 Mb false duplication detected at the end of the contig “ctg64” in the Duplex-based PECAT assembly (Nie et al, 2024)

We also computed unreliability percentages using ONT-R10 mappings. The numbers reported with the HMM-Flagger unfiltered sets ranged from 3.28% to 2.09%, while for the conservative sets they ranged from 1.07% to 0.36% (**Fig 3B**). These results indicate that ONT-based predictions are highly sensitive before filtering, which aligns with the previous base-level benchmarking results (**Fig 2B**). We then computed the base-level Jaccard similarities between the predictions made by different pairs of tools (**Fig 3C**). The lowest similarity scores were between NucFlag and conservative ONT-based HMM-Flagger. This may be explained partly by tool-specific mistakes and partly by platform-specific biases.

HMM-Flagger was able to detect large misassemblies in hard-to-assemble parts of the genome. In the Duplex-based PECAT assembly the 1.5Mb ends of two contigs were flagged as a false duplication (**Fig 3D and Supp Fig 13**). To further investigate this, we extracted the HiFi reads mapped to these regions and remapped them to the T2T-v1.1 assembly. They were mapped to an HSat3 array in chromosome 9 covering the large overlap between the mappings of the assembled contigs, which supports HMM-Flagger’s prediction (**Supp Fig 14**). In the T2T-v0.7 assembly HMM-Flagger detected a collapsed block (about 150 kb) in chromosome 13 upstream of an rDNA array. Mapping its HiFi reads to T2T-v1.1 showed nearly half aligned to the paternal copy of chromosome 22, indicating a region T2T-v0.7 failed to assemble (**Supp Figs 15, 16 and 17**). HMM-Flagger also detected false duplications (each about 500 kb long) in the middle of two active HOR arrays in the HPRC bake-off assembly[22] (**Supp Figs 18, 19, 20 and 21**).

### 2.4 Evaluating HPRC assemblies with HMM-Flagger

The Human Pangenome Reference Consortium (HPRC) generated 47 high-quality phased assemblies in their first release by using Hifiasm v0.14.1, Circular Consensus Sequencing (CCS) HiFi reads, and parental short reads[5]. The next release includes 231 assemblies, mostly produced with recent Hifiasm versions v0.19.7–v0.19.9 that incorporate ONT Ultra Long (UL) reads in addition to HiFi reads, which was essential for filling gaps and achieving higher contiguity compared to release 1[23]. The HiFi reads used for release 2 were corrected with DeepConsensus and the assemblies were further polished with DeepPolisher[24]. These improvements potentially make release 2 less prone to errors. To test this hypothesis, we applied HMM-Flagger to evaluate the two releases.

We computed unreliability percentages using HiFi and ONT mappings. HiFi-based results showed a decrease in average unreliability from 0.94% in release 1 to 0.38% in release 2, with false duplications showing the largest improvement (0.62% to 0.22%) (**Fig 4A**). The ONT-based conservative set also showed a drop in average unreliability from 0.66% to 0.16% (**Supp Fig 22**). To stratify results, we projected segmental duplication (SD) annotations from the CHM13-v2.0 reference [9] to each assembly (**Methods**) and used the AlphaAnnotation workflow[25] to identify centromeric and pericentromeric satellites (**Supp Note 3**) and then computed unreliability in the union of SDs and satellites, observing a significant improvement in release 2 (**Supp Fig 23 and 24**). The improvement in active HORs and highly homogeneous SDs (similarity *>* 99%) was more pronounced than in other SDs (**Supp Figs 25, 26 and 27**).

**Fig. 4.**
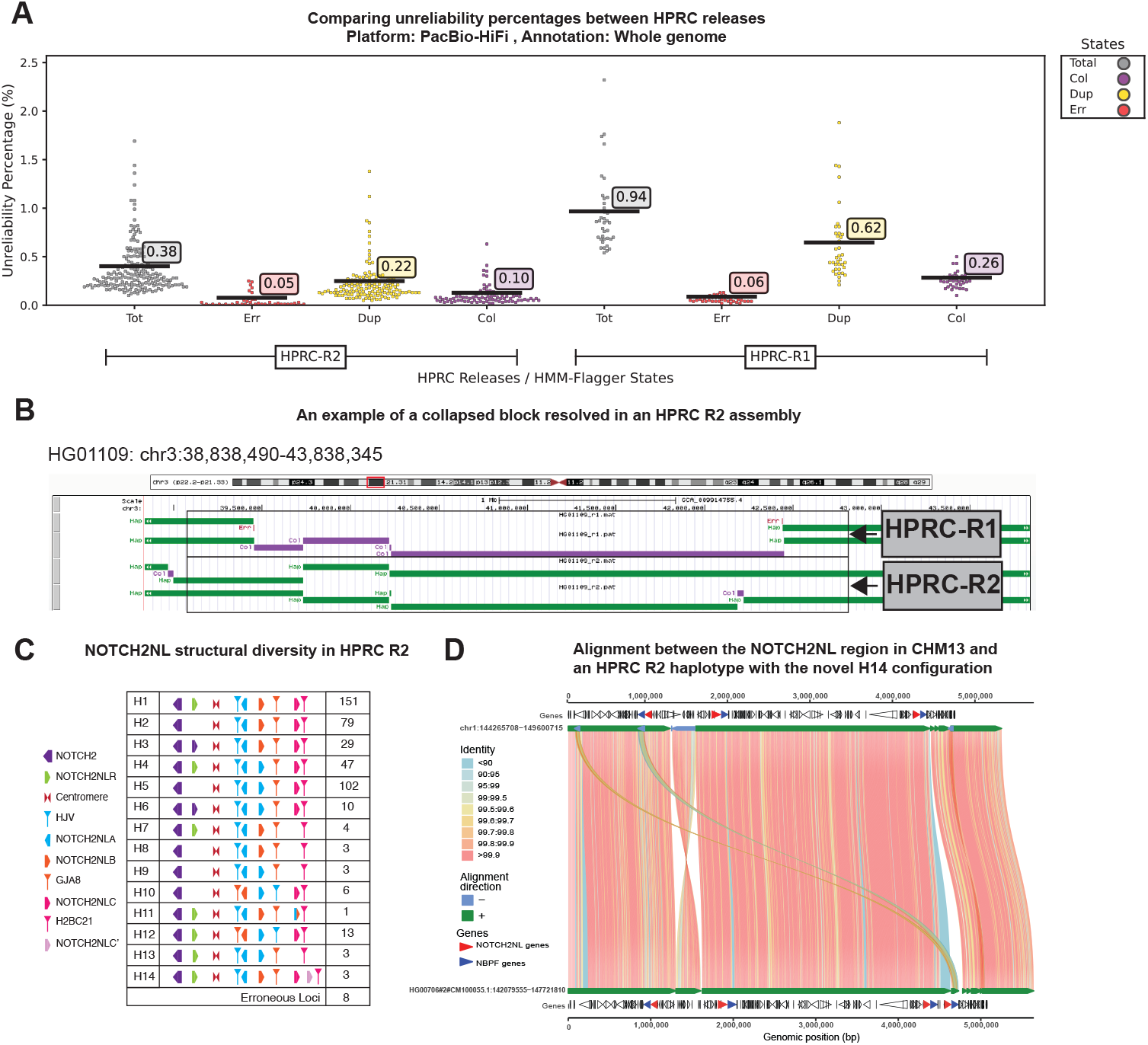
Evaluating HPRC assemblies and NOTCH2NL haplotypes with HMM-Flagger. **A)** Comparing unreliability percentages between HPRC releases across whole genome (predictions made with HiFi mappings). **B)** A 3 Mb maternal haplotype in chr3p22.1 is missing in HPRC-R1. It was detected as a collapsed block by HMM-Flagger. The maternal haplotype was correctly assembled in HPRC-R2. **C)** A summary table of various configurations of the NOTCH2NL region and their frequencies in the HPRC release 2 genome assemblies. **D)** An SVbyEye representation of the alignment between the NOTCH2NL regions of CHM13 and the maternal haplotype of HG00706 with an additional NOTCH2NLC’ copy (The rightmost red arrow). The NOTCH2NL genes are shown with red arrows and NBPF genes, which are known to always appear downstream of a NOTCH2NL gene, are in blue.

HMM-Flagger also revealed large misassemblies in release 1 that were corrected in release 2. Mapping assemblies to CHM13-v2.0 and projecting release 1 and release 2 flagged regions to its coordinates allowed a direct comparison. For the HG00673 sample, a 300kb false duplication upstream of the active HOR in chromosome 2 was resolved in release 2 (**Supp Fig 28**). For the HG01109 sample, a 3Mb maternal sequence in chromosome 3 p22.1 that was not assembled in release 1 caused a collapsed call. This missing sequence was assembled correctly in release 2 (**Fig 4B**).

### 2.5 Evaluating the NOTCH2NL gene in HPRC assemblies

To demonstrate that HMM-Flagger could be used to validate the assembly of complex regions, we conducted a comprehensive evaluation of the NOTCH2-N-terminus-like (NOTCH2NL) region in HPRC release 2 assemblies. The NOTCH2NL gene family arose from chromosome 1q21 segmental duplications and includes four genes: NOTCH2NLA, NOTCH2NLB, NOTCH2NLC, and NOTCH2NLR, which are implicated in human brain cortical expansion[26]. Like other complex segmental duplication regions, NOTCH2NL genes have been difficult to sequence and assemble with short-read technologies and have been incorrectly assembled in previous reference genomes[27].

We used an analysis framework similar to that described by a previous study for release 1[28]. Using the union of HiFi- and ONT-based HMM-Flagger results, our analysis predicts that 98% (454/462) of assemblies were accurately resolved in release 2, representing a substantial improvement over the 73% accuracy observed in release 1. Among these, 435 assemblies exhibited one of the 11 haplotype configurations previously characterized in the release 1 study, confirming the presence of known structural variants. Notably, this included the H8–H11 configurations, which were each observed in only a single assembly in the release 1 analysis (**Fig 4C**).

We identified three novel configurations. Configuration H12 represents a variant of H10, while configuration H13 represents a variant of H9; both are distinguished by the presence of an intact NOTCH2NLR copy. H12 was detected in thirteen haplotypes and H13 in three. Configuration H14 constitutes a modification of the canonical H1 configuration, characterized by an additional NOTCH2NLC’ copy on the positive strand downstream of NOTCH2NLC. This configuration was observed in three assemblies (two East Asian, one admixed American). The identity of the fourth copy was confirmed as NOTCH2NLC’ through SVbyEye analysis, protein/CDS multiple sequence alignment, and examination of surrounding duplicons (**Fig 4C, 4D** and **Supp Fig 29**).

HMM-Flagger successfully identified assembly errors in several samples. For instance, in HG03521, a misassembled contig resulted in false duplication in one haplotype, causing one haplotype to lack NOTCH2NLR while the other contained an additional copy. HMM-Flagger annotations prevented the misclassification of haplotype 1 as a valid H2 configuration (**Supp Figs 30 and 31**). In NA20870 haplotype 2, we detected two spurious copies of NOTCH2NLA and one spurious copy of NOTCH2NLB, all of which were flagged as false duplications (**Supp Fig 32**). Overall, HMM-Flagger facilitated rapid assessment of these structurally complex regions and enabled identification of 19 haplotypes with one of three novel configurations.

## 3 Discussion

In this study, we introduce HMM-Flagger a reference-free read mapping-based tool for evaluating the structural correctness of genome assemblies. A hidden Markov model (HMM) is at the core of its algorithm, which identifies coverage anomalies in the read mappings to the assembly and classifies them into three types of misassemblies: erroneous, falsely duplicated, and collapsed. The coverage values processed by HMM-Flagger are aggregated over fixed, pre-tuned windows, allowing the detection of structural errors at the resolution of those windows.

We showed that HMM-Flagger can identify a large number of synthetic misassemblies using either HiFi or ONT mappings (**Fig 2**). However two variables obviously influence the chances of accurate detection. First is length: longer events were easier to detect. The detection resolution was also limited by the window sizes tuned per platform (**Supp Figs 33-35**). Smaller windows improve resolution but increase false calls and runtime. The second correlating variable was misassembly type, with false duplications being hardest to flag (**Supp Fig 7**). This stems from two assumptions: that duplications lead to low MAPQ mappings, which is not always true when reads span non-duplicated regions, and that reads split evenly between copies, where bias toward one copy can mask duplications and cause them to be mislabeled as correct blocks or erroneous low-coverage blocks.

The misassemblies assessed in this study do not include all possible types of errors that can occur in a genome assembly. The current implementation of HMM-Flagger is mainly based on coverage and it is unlikely to detect balanced events that are not reflected in coverage, like inversions, unless there is a knock-on effect on read coverage. In the future, HMM-Flagger’s model could be improved in various ways to detect more types of errors. For example it could take into account more features such as apparent clustered read errors, the size of soft- or hard-clipped sequences and the frequency or presence of read k-mers versus assembly k-mers. The method also does not yet use split read mappings or attempt to detect breakpoints spanned by reads. The HMM itself could also be replaced with deep learning models, which have proven highly capable of learning context-specific patterns in read alignments [24, 29–31].

A major application of high-quality genome assemblies is providing accurate representations of gene families located in complex loci that in many cases have important clinical and medical implications[32]. As a case study, we used HMM-Flagger to examine NOTCH2NL genes assembled in HPRC release 2 and found three haplotypes with novel configurations, including duplication and gene conversion. Furthermore, HMM-Flagger successfully identified false duplications in some assemblies, preventing the false discovery of structural variants. Evaluation tools like HMM-Flagger can greatly aid in confirming the accuracy of assemblies in such loci and preventing false disease-risk associations of the assembled haplotypes.

## 4 Acknowledgments

We acknowledge Ivana Pačar (University of California, Santa Cruz) for valuable comments and feedback on the identification of the novel NOTCH2NL haplotypes. Research reported in this publication was supported, in part, by Weill Neurohub (to E.E.E.) and the National Human Genome Research Institute of the National Institutes of Health (NIH) under Award Number R01HG002385 (to E.E.E.). E.E.E. is an investigator of the Howard Hughes Medical Institute and Weill Neurohub. B.P. and UCSC personnel were in part supported by NIH grants. R01HG014490, U01HG010961, U24HG010262, OT2OD026682, and U24HG011853. The content is solely the responsibility of the authors and does not necessarily represent the official views of the NIH.

## 6 Author contributions

K.M. and B.P. devised the study. M.A., J.M.E., P.H., T.D.R., and B.P. drafted the manuscript. M.A. and J.M.E. developed the tool. M.A. performed data analysis. P.H., T.D.R., M.D., E.E.E., and S.S. conducted the NOTCH2NL analysis. J.L. and K.M. generated the HPRC assemblies and censat annotations, with H.L. contributing to censat annotation generation. A.C. collected the HG002 assemblies.

## 7 Methods

### 7.1 Input coverage data

The primary input for HMM-Flagger is the depth of read coverage along the assembly. Since BAM/SAM files contain more data than needed for this purpose, we defined a human-readable tab-delimited coverage format with header lines, which contains only relevant information for running HMM-Flagger. This format enables generating summary tables and stratifying final results using provided annotations. HMM-Flagger reads the coverage file and based on the input parameter `--windowLen`, it splits the genome into non-overlapping windows and computes the average coverage per window (window sizes for HiFi: 16kb, ONT-R9: 16kb, ONT-R10: 8kb). The window-level coverage values will be used as observations for parameter estimation and state inference. Files in this coverage format can be considered intermediate outputs between the input BAM and the final BED that contains HMM-Flagger predictions. Being at least two orders of magnitude smaller than BAM/SAM files, this format facilitates experimentation and reproducibility. More details about this format and the commands for creating it are provided in **Supp Note 1**.

### 7.2 HMM-Flagger model

One common way to evaluate a genome assembly is by mapping reads back to the assembly and investigating the regions where alignments appear abnormal. Depending on the types of misassemblies being examined, different metrics can be defined to measure alignment abnormalities. One metric is the depth of coverage which is particularly useful for detecting large structural misassemblies.

Collapsed blocks and false duplications are the two common types of misassemblies which can be detected by examining read coverage. Regions with extremely low or absent coverage may indicate highly erroneous assemblies, contig breakpoints, or structural misjoins. Based on the distinct coverage signatures of different assembly errors we developed HMM-Flagger. It employs a hidden Markov model (HMM) with four states: erroneous, falsely duplicated, haploid, and collapsed. which are shown with *S*_*E*_, *S*_*D*_, *S*_*H*_,and *S*_*C*_ respectively. Only regions assigned to the haploid state (*S*_*H*_) are considered reliably assembled. The emission probability densities of the states *S*_*D*_, *S*_*H*_, *S*_*C*_ are modeled by Gaussian Mixture Models (GMM) and for *S*_*E*_ by a truncated exponential distribution (**Fig 1A**). By performing inference over windows, HMM-Flagger provides posterior probabilities for each state.

Here we introduce the notation for formalizing the HMM. Let *T* denote the number of windows in a chunk (genome is split into multiple large chunks for parallelizing parameter estimation as explained in **Supp Note 4**) and in each window there is one observation shown with *x*_*t*_ which is a nonnegative integer. The whole observation sequence for one chunk is shown with ***x*** as below:

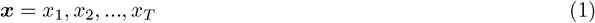

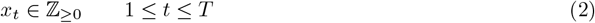

There are *N* states in the HMM (for HMM-Flagger *N* = 4). For each window *t*, the corresponding hidden state is denoted by *π*_*t*_, which can take any integer value from 1 to *N*. The whole state sequence for one chunk is represented by π as below:

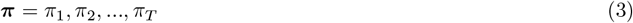

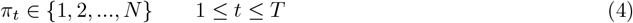

We define the state transition probability from state *i* to *j* as *a*_*ij*_. For the first window (*t* = 1), the initial state distribution is denoted by *a*_*i*_ showing the probability of starting from state *i*.

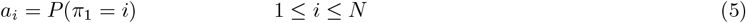

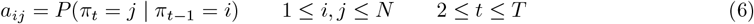

The probability of observing *x* at window *t*, given the hidden state *i*, is shown with *e*_*i*_(*x*):

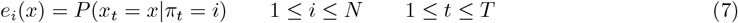

Under the assumption that reads are sampled uniformly and independently across the genome, the number of reads covering a given position follows a Poisson distribution, in which the mean and variance (*λ*) are equal. However, real data are usually overdispersed. Given the fact that a Poisson distribution with large *λ* is approximately Gaussian with mean and variance both equal to *λ*, using a Gaussian with decoupled mean and variance allows us to account for overdispersion. Therefore, emission densities for all states, except the erroneous state, are modeled using Gaussian distributions. This choice also simplified its implementation, since closed-form expressions exist for updating Gaussian parameters using the Expectation Maximization (EM) algorithm[16].

For *j ∈ {S*_*D*_, *S*_*H*_, *S*_*C*_*}, x* follows a Gaussian Mixture Model (GMM). The probability density function of the GMM is defined as:

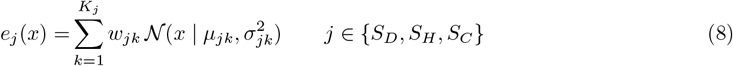

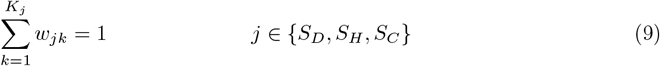

Here, *K*_*j*_ is the number of mixture components for state *j*. Each component is parameterized by a mixture weight (*w*_*jk*_), mean (*µ*_*jk*_) and variance (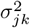). They will be estimated using the EM algorithm. For the falsely duplicated (*S*_*D*_) and haploid (*S*_*H*_) states there is only one mixture component (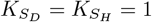).

For *i* = *S*_*E*_ which represent the regions with low or almost no coverage *x* follows a truncated exponential distribution. The probability density function of a truncated exponential distribution can be defined as below:

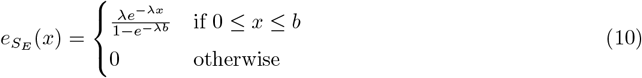

This distribution is non-zero only in the interval [0, *b*] and *λ* is the only parameter that has to be estimated.

### 7.3 Constraints on emission parameters

To determine the number of components for the collapsed state (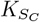), the genome-wide maximum and median coverage are used as below:

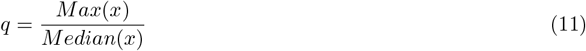

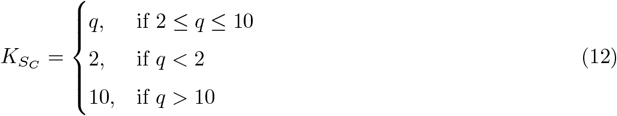

With equation (11) we ensure that 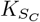 is neither too large nor too small. A very small value may underfit the coverage distribution of collapsed regions, while a large value increases the EM algorithm’s runtime, without significantly impacting the final inferred states.

With the model defined in the HMM-Flagger model section, there were some issues in estimating the parameters. The haploid state is usually predominant, especially when the assembly quality is high. As a result, in the estimation step the means of all states mutually converge to the median coverage, which is often close to the mean of the haploid state.

To ensure that each state represents the intended type of assembly region, we constrained the emission parameters of different states based on our prior expectations. These constraints are specified below with respect to the mean of the haploid state:

- As in equation (8) the collapsed state is modeled by a mixture of Gaussian distributions, whose mean values follow an arithmetic sequence with a step of 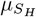 and a start point of 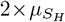. Although Gaussian approximations are used in our model, these ratios are justified by the facts that the coverage of a collapsed block is the sum of the coverages from the correctly resolved blocks, and that the sum of *k* independent Poisson variates with parameter 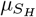 is a Poisson with parameter 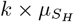.

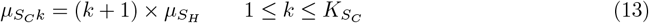
- The falsely duplicated state follows a Gaussian distribution, with its mean tied to half of the haploid mean. This ratio is justified by the fact that a duplicated sequence is most probably present in only one extra copy and the sum of two independent Poisson variates with parameter 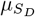 gives a Poisson with 2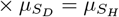.

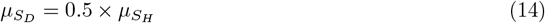
- As defined in equation (10) the erroneous state has two emission parameters: *λ* and *b* (truncation point). *λ* is an independent parameter, but the truncation point is set to 0.25 times the haploid mean.

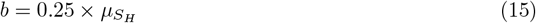

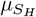 was used instead of 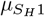 since *S*_*H*_ has only one mixture component.

The variances of the collapsed (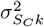), haploid (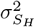) and falsely duplicated (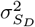) states are also related by the same factors and equations as in 13 and 14.

### 7.4 Constraints on transition parameters

The transition probabilities of the model are restricted based on the mapping quality of the read alignments. These constraints reduce the sensitivity of the model to minor fluctuations in coverage. For falsely duplicated regions the mapping quality is usually low especially if the two copies are highly similar and the mapper is not confident in choosing one copy to map reads to. To reflect this, we allow transition to the falsely duplicated state only when the ratio of the high mapping quality (MAPQ) coverage (alignments with MAPQ greater than 10) to the total coverage is lower than 0.25. In the following, we use the notation 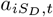 to show that the transition to *S*_*D*_ might not be allowed for all windows (*t* subscript shows the window index).

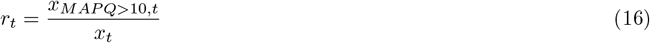

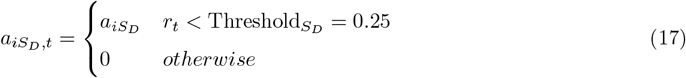

However in collapsed regions the mapping quality is usually high especially when one copy is completely missed so the mapper is falsely confident in mapping to the only assembled copy. Therefore, we allow transition to the collapsed state only when the ratio for high MAPQ alignment (defined in equation (16)) is greater than 0.75.

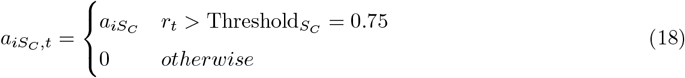

The default values mentioned earlier for the MAPQ-based thresholds are tuned manually. They can also be specified by the HMM-Flagger parameters `--maxHighMapqRatio` (for 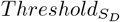) and `--minHighMapqRatio` (for 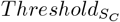).

### 7.5 Accounting for correlated coverage in nearby windows

With the HMM explained in the previous sections we observed that in some cases HMM-Flagger produces short false positive calls in which it flags correctly assembled regions as problematic. In order to increase its specificity, we revisited the assumptions of the initial model. One assumption was that given the state of the current position (*π*_*t*_) we can compute the probability of emitting the current observation (*x*_*t*_), which means that knowing the previous observation (*x*_*t−*1_) does not matter since given *π*_*t*_, the observations *x*_*t*_ and *x*_*t−*1_ are independent. However, this assumption is overly simplistic for coverage data. Reads can overlap multiple windows regardless of the underlying correctness of the sequence. This causes correlation between the coverage values that is not completely dependent on the correctness state, which violates the conditional independence of the HMM emissions.

The previous observation *x*_*t−*1_ in calculating the emission probability is necessary for developing a more accurate model. For instance, if the observation at the previous position *x*_*t−*1_ is known, and both the previous and current states are the same (*π*_*t−*1_ = *π*_*t*_) it is very likely that *x*_*t*_ emits a similar value. In other words, *x*_*t*_ and *x*_*t−*1_ are more correlated when *π*_*t−*1_ = *π*_*t*_ than when *π*_*t−*1_*≠ π*_*t*_. We can formulate is as below:

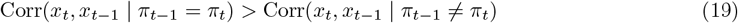

By considering such dependencies, we can develop a model that automatically adjusts coverage expectations to the local coverage fluctuations. To formalize this mathematically we augmented the HMM with Gaussian Autoregressive Process.

In a Gaussian Autoregressive Process we assume that each observation is a linear combination of previous observations and a probabilistic term that follows a Gaussian distribution. It can equivalently be formulated as one random variable whose mean value is a linear combination of previous observations and a constant. The number of previous observations that each observation depends on is the order of the process (shown with *P*). The order of the process we used for HMM-Flagger is one (*P* = 1). Therefore we can define the dynamic mean value for the states, *S*_*D*_, *S*_*H*_ and *S*_*C*_ as follows:

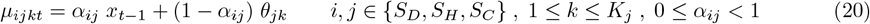

where *µ*_*ijkt*_ is the mean value of the *k*-th component of the *j*-th state given that the transition was from the *i*-th state (*π*_*t−*1_ = *i*). The *t* subscript indicates that the mean value can be variable between windows because of dependency to *x*_*t−*1_. The parameter *θ*_*jk*_ is not variable with respect to *t* and has to be estimated with EM algorithm. We can disable the autoregressive process by setting *α*_*ij*_ = 0 (in that case *µ*_*ijkt*_ would be constant and equal to *θ*_*jk*_ across all windows). As *α*_*ij*_ increases, the mean becomes more sensitive to local fluctuations in coverage.

We can also redefine equation (7) for emission probability to show the dependency on previous state and observation.

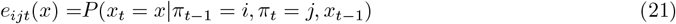

By combining equation (8) and equation (21) we can derive the new emission density function. As it is shown the variances of the Gaussian distributions 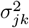 are not affected.

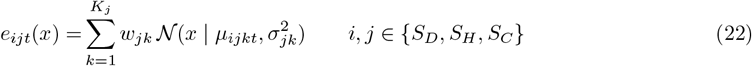

In **Supp Note 5** we prove that the forward-backward algorithm is still applicable with the new emission function.

### 7.6 Adjusting coverage expectation at contig ends

Genome assemblers cannot always assemble a chromosome into a single contig and might break it into multiple contigs. When reads are mapped back to the assembly, some that are mapped close to the contig ends might be dropped by the mapper. This can happen if the fraction of a read that could be mapped to the contig was smaller than a threshold. Therefore, even if a contig is error-free at its ends the read coverage might drop, which is misleading for HMM-Flagger.

In order to consider this coverage degradation, we need to adjust the coverage expectation based on closeness to the contig ends. We assume that a read will only be mapped to the contig if its alignment shares some fraction *η* (say, 1/2), then we can define lower *l*_*t*_ and upper *u*_*t*_ bounds for where the read alignment can start and end with respect to the contig ends. In these equations *Pos*(*t*) is the middle position of window *t*, G is the contig size and R is the average length of reads.

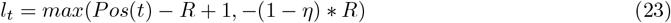

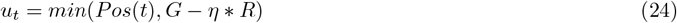

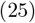

Then we can compute *β*_*t*_ which is a factor for adjusting the coverage expectation at contig ends.

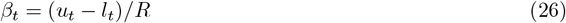

For a window that is far from the ends we expect *l*_*t*_ = *Pos*(*t*) *− R* and *u*_*t*_ = *Pos*(*t*) therefore 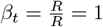 which means that there is no coverage degradation.

We will then use *β*_*t*_ to scale the emission parameters of the model. For states *S*_*D*_, *S*_*H*_ and *S*_*C*_ we scale the mean and variance in equation (22) and change it to the equation below:

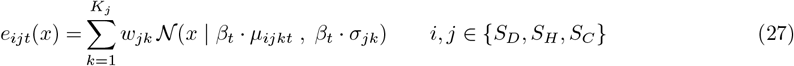

For the state *S*_*E*_ we scale the truncation point and *λ* as below:

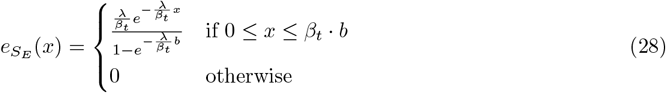

In order to tune *η* for different sequencing platforms we used simulated missamblies as explained in the section Simulating misassemblies with Falsifier (**Supp Fig 1**).

### 7.7 Incorporating coverage biases in HMM-Flagger

Based on the recent T2T-CHM13 paper, PacBio and ONT sequencing data exhibit coverage biases in some human satellites (HSats)[9]. More specifically, it was observed that PacBio data shows elevated coverage in HSats 2,3 and both PacBio and ONT data show reduced coverage in HSat1 with varying degrees (**Supp Fig 2**). Based on the investigations described in that paper it is very unlikely that these changes in read coverage are due to assembly errors because of multiple reasons. First, these biases were platform-specific for example while PacBio reads have excessive coverage in HSat2,3, the coverage level in ONT data was consistent with the genome-wide average. Second, the level of variation in coverage was consistent across the same type of arrays in different chromosomes. For example if there was 50% reduction of ONT coverage in HSat1 in one chromosome, it was around the same level in all HSat1 arrays across the genome. Although there was not one main reason for such platform-specific biases they were partially attributed to some factors like shorter or longer sequencing times and polymerase lengths. These biases mislead HMM-Flagger and impact the accuracy of its predictions especially for centromeric and pericentromeric satellites.

We added some features and parameters to HMM-Flagger to account for potential biases in different parts of the genome. The program bam2cov, which is for creating coverage files (**Supp Note 1**), can take a list of BED files for different annotations by passing a JSON file to the parameter `--annotationJson`. With the parameter, `--baselineAnnotation`, it takes the name of the annotation that should be used as the baseline to detect annotations with coverage biases. The parameter, `--runBiasDetection` will compute the median coverage for each annotation and compare it with the baseline annotation. If the median coverage of an annotation deviates from the baseline by greater than `--covDiffThreshold` then it will be reported as a biased annotation (e.g. HSat2 for PacBio HiFi). The blocks for each biased annotation will be parsed and assigned to a region index greater than 0. By default the region index of each block is 0 assuming that it does not have coverage bias. Region indices are saved in the 7th column of the output coverage file (**Supp Note 1**).

For the windows overlapping each region index we run the EM algorithm independently so that we have an independent set of HMM parameters for each biased annotation. We used the fixed probability of 0.25 for transitioning between adjacent windows with different region indices. This enabled us to use the original implementation of forward-backward algorithm without any need to break the iteration wherever we reach a window with a different region index. We only needed to compute the emission probability and intra-region transition probability based on region-specific parameters. This could be achieved easily by augmenting each observation with its pre-computed region index. To estimate region-specific parameters reliably we need to have a decent number of observations per region index. Therefore we only select an annotation as biased if its total length is greater than `--minBiasLength` (A parameter of bam2cov with the default value of 100K). For informing the model of potential coverage biases we need reliable annotations of human satellites.

It is more convenient to annotate the human satellites using an annotation tool designed for this aim (e.g. HumAS-HMMER)[33]. However it is also possible to do this by using a well annotated reference like T2T-CHM13 and the assembly alignments to this reference. We developed a multi-threaded Python script, project blocks multi thread.py, that takes the alignments in PAF format and also a bed file in the reference coordinates. It outputs the projections of the reference blocks into the assembly coordinates by iterating over the CIGAR string. This script is available in the HMM-Flagger Docker image and can also be useful in any other analysis that needs such projections.

### 7.8 Filtering predictions with self-homology mappings

As described earlier we were able to increase the accuracy of HMM-Flagger by modifying the model in several ways such as putting constraints on HMM parameters, augmenting the model with Gaussian Autoregressive Process, adjusting coverage expectation at contig ends and incorporating coverage biases in human satellites. However even with these improvements there were still some unrelsoved false positive predictions. Here we explain how self-homology mappings were used to filter predictions and create a conservative call set with higher specificity.

Based on the types of misassemblies described in the section, HMM-Flagger model, a false duplication results in having redundant copies of an assembled block, whereas a collapsed region typically leads to a complete absence of the homologous region. Therefore, we can map contigs against each other to obtain self-homology mappings and use them as a secondary check for the blocks flagged as falsely duplicated or collapsed by HMM-Flagger. For a region flagged as collapsed, there should be no mappings from other contigs. However for false duplications, there should be more than one mapping and at least one of them should originate from another block also flagged as falsely duplicated.

Based on the above criteria, we wrote a Python script, named filter hmm flagger calls.py, that takes a BAM/SAM file of self-homology mappings and also the HMM-Flagger BED file as input. After filtering, it will output a BED file with conservative predictions. For the filtering process, we associate each region of the genome with all blocks that map to it and apply the following four rules, in order:

- If a region has more than 2 associated blocks we assume that they do not represent true homology. Since it is hard to find true homology relations in such cases we try to be conservative and report these regions as haploid even if they were originally marked as collapsed or falsely duplicated.
- If a region was marked as falsely duplicated and at least one of the associated blocks was also flagged as falsely duplicated, we keep the original prediction; otherwise we assume that is a FP and report it as haploid.
- If a region was marked as collapsed and there was either no associated block (no mapping) or there was at least one associated block marked as erroneous we keep the original prediction; otherwise we assume that is a FP and report it as haploid. The reason for considering the case with erroneous label is that sometimes all copies of homologous blocks are assembled but one copy is highly erroneous or fragmented, so that we label the correct copy as collapsed.
- For all other cases we label as haploid whatever the original prediction was.

The self-homology mappings can be created by executing minimap2 with the parameters, `-D -ax asm5`, and using the diploid assembly both as reference and query.

The filtering step described above has some limitations. First, self-homology mappings created with minimap2 are not accurate enough for highly repetitive arrays like Higher Order Repeats in centromeres, which leads to filtering based on wrong homologies. Second, if there is a biologically real large deletion (at least at the order of a read length) in one haplotype and the other haplotype is marked as collapsed by mistake, we cannot fix this FP even after filtering with self-homology mappings.

### 7.9 Simulating misassemblies with Falsifier

In order to measure the sensitivity and specificity of HMM-Flagger predictions we need a set of assemblies with known misassembly types and coordinates. We developed a program named Falsifier that takes a diploid assembly along with the mappings of the two haplotypes against each other and uses them to simulate three types of misassemblies: erroneous blocks with high base-level error rate (4% by default), false duplications, and collapsed blocks. The locations of misassmblies are selected randomly however the desired numbers and lengths of misassembled blocks per misassembly type can be specified with the `--misAssemblyTsv` flag. Falsifier’s output includes a diploid falsified assembly in FASTA format and the coordinates of induced misassemblies in a BED file.

To ensure that the misassemblies induced with Falsifier are the only misassemblies in the output falsified assembly, there should not be any error in the input assembly. One of the most accurate diploid assemblies currently available for human is the HG002-T2T-v1.1 assembly released by the Telomere-to-Telomere (T2T) Consortium. Therefore we used this assembly for inducing misassemblies with Falsifer.

Falsifier takes the mappings of one haplotype (e.g. paternal) to the other (e.g. maternal) with the `--paf` parameter. For creating these mappings we used two mappers minimap2 and centrolign. Centrolign is a multiple (and pairwise) sequence aligner designed for long tandem repeats like alpha satellites [34]. For mapping centromeric and pericentromeric arrays we used centrolign-v0.2.1 (with default flags) and all other parts of the genome were mapped with minimap2-v2.28 (with `-x asm5 –eqx` flags) (**Supp Note 6**).

Falsifier takes a PAF file with haplotype mappings and performs the steps listed below:

- Parses the alignments of one haplotype versus the other for HG002-T2T-v1.1 assembly.
- Splits the alignments wherever there is a long InDel (`--maxGapLength` with 500 as the default)
- Ignores the short alignments (`--minAlignmentLength` with 50kb as the default)
- Ignores the regions with more than one alignment. It does not discard the whole alignment instead eliminates only the segments of each alignment with a coverage greater than 1. This step ensures that Falsifier keeps only the parts of the genome with a one-to-one homology between the haplotypes.
- Splits each contig into consecutive non-overlapping blocks based on the existence or absence of a one-to-one homology between the haplotypes. Therefore each contig can be represented by a sorted list of consecutive blocks each of which is augmented by at most one block from the other haplotype. In other words it is a list of homology relations. In the actual implementation, a homology relation will be empty if there is no valid one-to-one alignment for a block. Collapsed blocks are induced only in the blocks with non-empty homology relations since we need to remove a sequence from one haplotype and flag the homologous sequence in the other haplotype as collapsed.
- Uses the lists of homology relations to keep track of assembly coordinates after creating each misassembly. Misassemblies are created sequentially in Falsfier.

For creating each misassembly type Falsifier selects a random block with the desired misassembly length and performs one of these actions based on the misassembly type:

- **Collapsed block**: It can be created only within regions with non-empty homology relation. The randomly selected block is removed and the related contig is split into two smaller contigs at the end points of the removed block. Using the one-to-one mapping Falsifier finds the homologous block in the other haplotype and flag it as collapsed.
- **False duplication**: It can be created in any block with either empty or non-empty homology relation. The randomly selected block is duplicated and placed into a new contig. Both the original copy and the new copy are flagged as false duplication. We expect both copies to be flagged by HMM-Flagger.
- **Erroneous block**: Similar to false duplication this can occur in any block. Random SNP errors are induced in the randomly selected block. The rate of SNP errors can be specified by the `--singleBaseErrorRate` flag (We used a rate of 0.04). The current implementation does not induce InDel errors.

A JSON file listing annotation BED files can be provided to Falsifier with the `--annotationsJson` flag. The coordinates of annotations for the input assembly will be adjusted to match the output falsified assembly. The adjusted coordinates will be saved in new BED files. This enables stratifying the later benchmarking results by different annotations (e.g. seg dups or satellites). It is also possible to specify the number and lengths of misassemblies per annotation. This feature is useful for controlling the misassembly rate for hard-to-assemble parts of the genome for example, segmental duplications. The `--minOverlapRatio` flag sets the minimum fraction of a misassembled block that must overlap an annotation to be counted in that annotation (default: 0.75). If the requested misassembly lengths exceed what is possible within an annotation, Falsifier will induce as many as feasible and skip the rest. More details on the annotations that were used for misassembly generation and also the number and length of misassemblies can be found in **Supp Note 7**.

### 7.10 Tuning hyperparameters of the model

In the HMM augmentend with Gaussian Autoregressive Process the mean of the emission distribution depends not only on the current state but also on the previous state and the previous observation. This dependency applies only when the current state is one of the three states—*S*_*D*_, *S*_*H*_, and *S*_*C*_ (All except *S*_*E*_). As it is defined in Equation (20) the dependency hyperparameter *α*_*ij*_ quantifies the impact of the previous observation, *x*_*t−*1_, on the mean of the emission distribution given that the transition is from state *i* to *j*. Since there are 4 possible previous states (including *S*_*E*_) and 3 current states, this results in 4 *×* 3 = 12 distinct *α*_*ij*_ hyperparameters, which are shown in the matrix below:

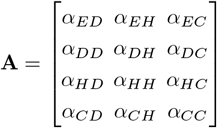

For tuning the hyperparameter matrix, **A**, we created a training data set containing assemblies with known misassembly coordinates. We could then optimize the matrix in a way that the HMM-Flagger’s predictions are as close as possible to the truth locations and labels of the misassemblies. We created such falsified assemblies by applying the program Falsifier on the HG002-T2T-v1.1 assembly. The Falsifier algorithm, along with its inputs and outputs, is described in the Methods section, simulating misassemblies with Falsifier.

The training data set contained the read mappings to two falsified assemblies; one with 0.87% misassembly rate and another with 3.32%. The reads from each sequencing platform were mapped to them at two coverage levels; 40x and 20x. Each mapping was performed using two different mappers, minimap2 and winnowmap, to avoid overfitting the hyperparameters to a specific mapper. This also allowed us to fairly choose one mapper for large-scale use of HMM-Flagger on HPRC assemblies (**Supp Figs 36, 37 and 38**). In total, the training data set comprised 8 BAM files—one for each combination of misassembly rate, coverage level, and read mapper.

HMM-Flagger was run on each training BAM file separately and the resulting prediction BED files were then compared against the related truth BED files. We used the three metrics described in **Supp Note 8**, to quantify the agreement between predictions and ground truth. The final performance score was computed as the average of these metrics across all eight BAM files.

Now we can consider the whole process of running HMM-Flagger and computing the performance score as an objective function by which we can optimize **A. Fig 1F** shows the objective function, *f* (**A**) that maps the hyperparameter matrix, **A** (blue matrix on the left side), to the performance score, *Y* (green rectangle at the bottom). Given the properties of the objective function and the structure of the problem, Bayesian Optimization (BO) is a suitable approach for finding the optimal value of **A**. Some of these properties are listed below [17, 35]:

- It is not feasible to find a close-form mathematical expression for *f* (*A*) so that it can be considered as a black-box function.
- Computing *f* (**A**) is time-consuming, as each evaluation requires running HMM-Flagger on eight BAM files. Thus, we aim to find an optimal (or near-optimal) value using as few evaluations as possible.
- We can only compute the value of *f* (*A*) and it is not possible to take first- or second-order derivatives. Therefore we cannot use methods like gradient descent.
- We want to perform a global optimization rather than local.

One of the widely used BO algorithms is Efficient Global Optimization (EGO) [17], which we used for optimizing HMM-Flagger hyperparameters. It works by taking some initial randomly selected points from the feasible space of **A** and then after evaluating f(A) for those points it fits a surrogate to approximate *f* (*A*) using Gaussian Process regression. This surrogate will provide an approximation of any unobserved point and also a confidence interval which shows the uncertainty of the model in each point. Then the EGO algorithm can select the next point to evaluate by maximizing an acquisition function on the surrogate (This function is also referred to as infill sampling criterion). One common acquisition function is the expected improvement which measures the expectation of improvement compared to the maximum observed value so far (shown with 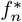).

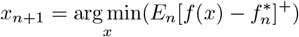

After multiple rounds of selecting points and updating the surrogate the EGO algorithm stops and outputs the optimum value observed among the performed evaluations (**Supp Fig 39**).

To estimate the optimal hyperparameters we wrote this script which uses SMT library, a python API for the EGO algorithm[36].

https://github.com/mobinasri/flagger/blob/v1.2.0/programs/src/tune_alpha_hmm_flagger.py

### 7.11 Evaluating the NOTCH2NL configurations

To validate the HPRC release 2 assembly set, we confirmed correct and contiguous assembly of sequences spanning the NOTCH2NL paralogs on both chromosomal arms, including non-NOTCH2NL gene markers. Assemblies containing large gaps in this region or where these regions were distributed across multiple contigs were excluded from analysis. HMM-Flagger was subsequently applied to identify assemblies containing structural errors in these regions; all assemblies passing this validation were included in the final set of 454 assemblies.

To determine NOTCH2NL locus configurations, we employed multiple complementary approaches in conjunction with CAT2 [37] gene annotations on HPRC release 2 assemblies. The NOTCH2NL transcript coding sequences (CDS) from the CHM13 reference were aligned against all assemblies using BLAT to identify the transcript that matched each paralog the best based on alignment scores. Maximum likelihood phylogenies were then constructed using intronic sequences from all NOTCH2NL transcripts in each assembly and CHM13; multiple sequence alignments were generated using MAFFT [38], and phylogenetic trees were inferred using IQ-TREE2 [39]. Cases exhibiting conflicts among the three methods were considered potential gene conversion candidates and were manually inspected using the IGV and CDS/protein multiple sequence alignment visualization.

