## Supplementary Notes for "Evaluating genome assemblies with HMM-Flagger"

### Supplementary Notes
### Evaluating genome assemblies with HMM-Flagger

Mobin Asri[1*], Jordan M. Eizenga[1], Prajna Hebbar[1], Taylor D. Real[2], Julian Lucas[1], Hailey Loucks[1], Alessandro Calicchio[1], Mark Diekhans[1], Evan E. Eichler[2,3], Sofie Salama[1], Karen H. Miga[1], Benedict Paten[1*]

[1]Genomics Institute, University of California Santa Cruz, Santa Cruz, CA, 95060, USA.
[2]Department of Genome Sciences, University of Washington School of Medicine, Seattle, WA 98195, USA.
[3]Howard Hughes Medical Institute, University of Washington, Seattle, WA 98195, USA.

Contributing authors:;

### 1 Human-readable and binary coverage formats

The coverage format begins with multiple header lines each starting with a hash sign (#). After each hash sign there is a header tag separated from its related value with a colon (:). These header lines provide details such as the number of regions with different coverage modes, the coverage mode values, the names and indices of the annotations represented in the file, the presence or absence of truth/prediction labels, and the average alignment length (**Supp Table 17**). The block lines for each contig are sorted by coordinate after a definition line that begins with a greater-than sign (>) showing the contig name and its total size. Each subsequent block line has at least 7 tab-delimited columns showing start and end coordinates, coverage values, the indices of overlapping annotations, and also the region index this block belongs to (**Supp Table 18**). Region indices are essential for tracking areas that may have coverage biases. It allows HMM-Flagger to estimate independent parameters for blocks with each region index, which prevents from being misled by platform-specific coverage biases (explained in **Methods: Incorporating coverage biases in HMM-Flagger**).

The 8th and 9th columns are optional and included only if the truth and prediction header tags are set to true. These columns contain truth and prediction labels, which are useful for generating summary tables with performance statistics. Combined with annotation columns, they allow for stratified summary tables. For example we can calculate what percentage of collapsed bases in segmental duplications were correctly predicted by HMM-Flagger.

Coverage files can be generated using a multi-threaded program named bam2cov available in the HMM-Flagger Docker image. Primary inputs for bam2cov are a BAM file containing read alignments and a JSON file listing annotation BED files. Leveraging multi-threading for I/O operations, bam2cov achieves high-speed performance, especially on storage disks with high bandwidth, such as Solid-State Drives (SSD) (**Supp Fig 40**). For instance, on an SSD storage disk, running bam2cov with `--threads 16` on a 165 GB BAM file took only 4 minutes and used 7.75 GB of memory with a CPU usage of 1101.0%. These are the commands used for creating a gz-compressed coverage file:

```
# Go to the working directory
# Put fasta, bam and bam index files in this directory
cd ${WORKING_DIR}

# Create fasta index assuming that file is not gz-compressed
samtools faidx ${FASTA_FILE}

# Create a bed file covering whole genome
cat ${FASTA_FILE}.fai | \
    awk '{print $1"\t0\t"$2}' > whole_genome.bed

# Put the path to the whole-genome bed file in a json file
echo "{" > annotations_path.json
echo \"whole_genome\" : \"${PWD}/whole_genome.bed\" >> annotations_path.json
echo "}" >> annotations_path.json

# Convert bam to cov.gz with bam2cov program
Docker run -it --rm -v${WORKING_DIR}:${WORKING_DIR} mobinasri/flagger:v1.2.0 \
  bam2cov --bam ${WORKING_DIR}/${BAM_FILE} \
          --output ${WORKING_DIR}/coverage_file.cov.gz \
          --annotationJson ${WORKING_DIR}/annotations_path.json \
          --threads 16 \
```

```
--baselineAnnotation whole_genome
```

The path to any additional annotation can be added to the input annotation JSON file. For visualizing in a software like Integrative Genome Browser (IGV), coverage format can be converted to BED or WIG formats using the coverage_format_converter program available in the HMM-Flagger Docker image (**Supp Table 19**).

HMM-Flagger can receive an input file in either BED or coverage format (either gz-compressed or uncompressed). It then computes the average of coverage values in windows whose size is specified by the `--windowLen` input parameter (the default size is 16kb for HiFi and ONT-R9 and 8kb for ONT-R10, )(**Supp Fig 33, 34, and 35**). Each window-level coverage value will be considered as an observation for HMM-Flagger (**Fig 1**). The annotation list for each window will be the union of all the annotations overlapping the window.

For computing the data related to each window, HMM-Flagger first creates an index of the coverage file. This index splits the genome into chunks of size 20Mb and keeps the file positions of where the first entry of each chunk starts in the input coverage file. It enables creating a thread pool and computing window-level data for each chunk in a separate thread. This multithreading accelerates the process of transforming base-level data into window-level data including coverage values, annotation lists, and region indices. HMM-Flagger will then use window-level data for estimating the HMM parameters and inferring misassembled regions (**Fig 1B**).

A human-readable coverage file can be converted to a binary coverage format containing window-level data. This conversion can be done using the coverage_format_converter program available in the HMM-Flagger Docker image. A binary coverage file can be given directly as input to HMM-Flagger, which leads to skipping the indexing and window averaging steps.

### 2 Selecting six HG002 assemblies from literature

To show that HMM-Flagger can detect actual assembly errors, we selected six HG002 haplotype-resolved assemblies that have been created recently using different methods and sequencing platforms.

The first assembly was taken from the bake-off paper accompanying the first release of the human pangenome project[1]. For creating this assembly, HG002 PacBio-HiFi reads (130x coverage) and parental Illumina reads (from HG003 and HG004) were given to Hifiasm (v0.14.1), a state-of-the-art assembler designed for PacBio-HiFi reads[2, 3]. It used HiFi reads for creating assembly graph and parental reads for phasing bubbles in the graph. The output of Hifiasm was scaffolded using Bionano and OmniC HiC reads. The gaps in the resulting scaffolds were then filled using the contigs generated with Flye assembler and ONT-UL reads.

The second assembly was generated with PECAT (Phased Error Correction and Assembly Tool) using ONT-R10 Duplex data [4]. PECAT first corrects errors in long, noisy reads with a POA-based method, then builds two string graphs in consecutive rounds: the first produces a haplotype-collapsed assembly, which is used to identify and remove inconsistent read overlaps, and the second generates haplotype-resolved sequences from the corrected overlaps. Although PECAT supports PacBio HiFi and ONT-UL reads, we specifically used the ONT-Duplex assembly to demonstrate that HMM-Flagger can detect errors in ONT-Duplex–based assemblies.

The third assembly was created using Verkko assembler (v1.3.1), which is a state-of-the-art assembler designed to generate haplotype-resolved assemblies using various combinations of sequencing data types[5]. The HG002 Verkko assembly evaluated in this study was generated using PacBio-HiFi ($>$ 10kb long, >99.9% accurate reads), ONT-UL (>100kb long, >90% accurate reads ) and parental Illumina reads (from HG003 and HG004). The initial assembly graph was created with PacBio HiFi reads, after which ONT-UL reads were mapped to the graph for phasing bubbles, filling gaps, and resolving loops and tangles. Parental markers from Illumina reads were then used to generate haplotype-specific contigs.

The last three assemblies were old (v0.7 and v0.9) and latest (v1.1) versions of the assemblies released by the Q100 project of the Telomere-to-Telomere (T2T) consortium. The v0.7 assembly, released on the Q100 GitHub page in November 2022, was created by combining two earlier Verkko assemblies followed by manual curation, resulting in a Quality Value (QV) of 60. The v0.9 assembly, released in July 2023, was created using improved HiFi and ONT data along with polishing and patching steps, which increased the QV to 71.8. The latest v1.1 release represents, to our knowledge, the most accurate diploid assembly generated to date [6].

### 3 Creating and using censat annotations

Due to platform-specific coverage biases across different pericentromeric and centromeric satellite classes, HMM-Flagger takes a cenSat annotation BED file as an input. The tool extracts tracks corresponding to individual satellite families and estimates a separate set of parameters for each satellite (e.g., HSat2). The cenSat annotation inputs for the HPRC assemblies and the six HG002 test assemblies were generated using the AlphaAnnotation workflows available at https://github.com/kmiga/alphaAnnotation. Specifically, we used the `centromereAnnotation.wdl` workflow (https://github.com/kmiga/alphaAnnotation/blob/main/cenSatAnnotation/centromereAnnotation.wdl), and the resulting per-haplotype `cenSatAnnotations` BED files were provided to the `cntrBed` argument of the HMM-Flagger WDL, with `enableDecomposingCntrBed` set to `true`.

### 4 Implementing the EM algorithm

We explained earlier that converting a BAM file to a coverage file and also summarizing base-level data into window-level data are multi-threaded (**Supp Note 1**). In addition to that, parameter estimation and state inference in HMM-Flagger are also parallelized. After loading window-level data into memory they are stored into chunks of size 20Mb. For example for a window size of 8 Kb there will be 2,500 windows in a single chunk (or equivalently 2,500 observations). HMM-Flagger will then create a thread pool in which each thread will run the forward-backward algorithm for one chunk of windows. It enables running the EM algorithm for multiple parts of the genome simultaneously and therefore accelerating the whole process in multi-core environments. The chunk size and window size can be modified with the parameters `--chunkLen` and `--windowLen`.

After each iteration of the forward-backward algorithm, we calculate the posterior probability of each observation being emitted from a particular mixture component of a state (The E-step of the EM algorithm). By using these posterior probabilities we can update the model parameters including the means, variances, and weights of the emission distributions, as well as the transition probabilities between states (The M step of the EM algorithm). In the next iteration we repeat the same process using the updated parameters (**Fig 1C**). The outer loop of this EM algorithm is the same as the Baum-Welch algorithm; however, after each iteration, the GMM parameters are estimated using an inner EM algorithm for which we used a closed-form expression from Rabiner's tutorial on HMM[7].

After each iteration we calculate the relative difference between the current values of the model parameters and those from the previous iteration. The algorithm stops once this difference is smaller than the convergence tolerance (`--convergenceTol 0.001` as the default value) or the number of iterations exceeded a predefined threshold (`--iterations 100` as the default value). Once the EM algorithm stopped, we calculate the posterior probabilities for the last time and use them to find which state (out of the 4 states) obtains the highest probability in each window. The inferred state for each window will be reported in the output BED file.

### 5 Forward-Backward algorithm for the HMM with Gaussian Autoregressive Process

We will prove here that with the model developed in the Methods section (Gaussian Autoregressive Process), we can still use the forward-backward algorithm for computing posterior probabilities.

The original forward recursion without autoregressive process works as follows:

$$f_j(t+1) = e_j(x_{t+1}) \sum_i f_i(t)\, a_{ij} \tag{1}$$

With the autoregressive process we have to replace $e_j(x_{t+1})$ by $e_{ij(t+1)}(x_{t+1})$ (Defined in equation (22) in Methods). Therefore, the recursive expression can be written as follows:

$$f_j(t+1) = e_{ij(t+1)}(x_{t+1}) \sum_i f_i(t)\, a_{ij} \tag{2}$$

The forward algorithm works with the recursively defined function $f_i(t)$, which is the probability of the observed sequence up to and including $x_t$ requiring that $\pi_t = i$.

$$f_i(t) = P(x_1, \ldots, x_t, \pi_t = i) \tag{3}$$

Now we show that with the recursive expression (2), the inductive premise defined in (3) also holds for $f_j(t+1)$. Instead of $e_{ij(t+1)}(x_{t+1})$ we used the notation $e_{ij}(x_{t+1} \mid x_t)$ to show that the emission probability is conditional on $x_t$.

$$\begin{aligned} f_j(t+1) &= \sum_i e_{ij}(x_{t+1} \mid x_t)\, f_i(t)\, a_{ij} \\ &= \sum_i e_{ij}(x_{t+1} \mid x_t) \cdot P(x_1, \ldots, x_t, \pi_t = i) \cdot P(\pi_{t+1} = j \mid \pi_t = i) \\ &= \sum_i e_{ij}(x_{t+1} \mid x_t) \cdot P(x_1, \ldots, x_t, \pi_t = i, \pi_{t+1} = j) \\ &= \sum_i P(x_1, \ldots, x_t, x_{t+1}, \pi_{t+1} = j, \pi_t = i) \\ &= P(x_1, \ldots, x_{t+1}, \pi_{t+1} = j) \end{aligned} \tag{4}$$

Next we repeat a similar procedure for the backward algorithm.

The original backward recursion without autoregressive process works as follows:

$$b_i(t-1) = \sum_j a_{ij}\, e_j(x_t)\, b_j(t) \tag{5}$$

Similar to the forward algorithm, because of using autoregressive process, we rewrite the recursive expression with $e_{ij(t+1)}(x_{t+1})$:

$$b_i(t-1) = \sum_j a_{ij}\, e_{ijt}(x_t)\, b_j(t) \tag{6}$$

The backward recursion works with the recursively defined function $b_i(t)$:

$$b_i(t) = P(x_{t+1} \dots x_T \mid \pi_t = i, x_t) \tag{7}$$

In contrast to the usual definition, we explicitly mention $x_t$ in the conditional part of equation (7) because of using the autoregressive process.

Now we show that with the recursive expression (7), the inductive premise defined in (6) also holds for $b_i(t-1)$.

$$\begin{aligned}
b_i(t-1) &= \sum_j a_{ij}\, e_{ij}(x_t \mid x_{t-1})\, b_j(t) \\
&= \sum_j a_{ij}\, P(x_t \mid \pi_t = j, \pi_{t-1} = i, x_{t-1}) \cdot P(x_{t+1} \dots x_T \mid \pi_t = j, x_t) \\
&= \sum_j a_{ij}\, P(x_t x_{t+1} \dots x_T \mid \pi_t = j, \pi_{t-1} = i, x_{t-1}) \\
&= \sum_j P(\pi_t = j \mid \pi_{t-1} = i)\, P(x_t \dots x_T \mid \pi_t = j, \pi_{t-1} = i, x_{t-1}) \\
&= \sum_j P(x_t \dots x_T \mid \pi_t = j, \pi_{t-1} = i, x_{t-1}) \cdot P(\pi_t = j \mid \pi_{t-1} = i) \\
&= P(x_t \dots x_T \mid \pi_{t-1} = i, x_{t-1})
\end{aligned} \tag{8}$$

### 6 Mapping haplotypes for Falsifier

In order to simulate misassemblies with Falsifier we need the mappings between the haplotypes of a reference-quality assembly like HG002-T2T-v1.1. To create a PAF file with such mappings we wrote a script named montage_mapper.py. This script takes a 7-column mapping BED file that specifies which parts of the genome should be mapped with minimap2 or centrolign. For each row, the first 6 columns specify the coordinates of a pair of blocks and the 7th column should be either "minimap2" or "centrolign" showing which mapper to use. We created such a mapping BED file by taking the satellite coordinates from the censat annotation of HG002-T2T-v1.1 and using the commands mentioned in this Github page:

https://github.com/mobinasri/research_notes/tree/main/Flagger_HG002_T2T/Montage_Mapper_BED_HG002_v1.1.

The BED file is available through this link: https://s3-us-west-2.amazonaws.com/human-pangenomics/submissions/e093fd72-e31a-11ee-b020-27964ee37032--flagger_test_files/misassembly_simulation_oct_2024/all_block_pairs.bed.

This BED file along with the HG002-T2T-v1.1 haplotypes is provided to `montage_mapper.py` to create a PAF file.

Here is the command we ran for mapping haplotypes with `montage_mapper.py`:

```
Docker run -v${WORKING_DIR}:${WORKING_DIR} --rm mobinasri/flagger:v1.2.0 \
        python3 /home/programs/src/montage_mapper.py \
        --bed ${INPUT_MAPPING_BED} \
```

```
    --hap1 ${PAT_T2T_FASTA} \
    --hap2 ${MAT_T2T_FASTA} \
    --outDir ${OUTPUT_DIR} \
    --prefix "HG002_v1.1_mat_to_pat" \
    --threads ${THREADS} \
    --minimap2Path "${MINIMAP2_BINARY_PATH}" \
    --centrolignPath "${CENTROLIGN_BINARY_PATH}" \
    --minimap2Params "-x asm5 --eqx" \
    --centrolignParams " "
```

#### 7 Misassembly datasets for training and testing

Misassembly rate is defined as the total length of misassembly blocks divided by the total length of diploid assembly which is approximately 6.2 Gb for human. To benchmark HMM-Flagger on assemblies with different misassembly rates we designed two tables specifying the counts and lengths of misassembly blocks. Using Table 1 and Table 2 we could generate falsified assemblies with misassembly rates of 3.32% and 0.87% respectively. These tables also show that our simulation generated misassemblies of varying length (from 40Kb to 320 Kb) overlapping with different annotations. We used 4 annotations specified in these tables:

- **active_hors**: The Higher Order Repeats of alpha satellites taken from the censat annotation
- **hsats**: The human satellites 1, 2 and 3 taken from the censat annotation
- **sd_no_hor_or_hsat**: Segmental duplications lifted from HG002-T2T-v1.0.1 to HG002-T2T-v1.1. All satellites included in active_hors and hsats were excluded from this annotation.
- **other**: The rest of the genome excluding the other 3 annotationsm mentioned above.

Since the lengths of one-to-one alignments in HORs and HSats were limited, we were not able to induce misassemblies longer than 80Kb in those regions. This is partly due to the high rate of divergence between the haplotypes in centromeric and pericentromeric regions and partly due to the inability of centrolign to provide contiguous alignments.

We created two sets of falsified assemblies; one set for tuning the hyperparameters of HMM-Flagger (the training data set) and the other one for testing HMM-Flagger with tuned hyperparameters (the test data set). The training and test data sets should be generated independently to show that the hyperparameters are not overtrained to the training data set. Since Falsifier selects misassembly locations randomly, each run produces a new, independent assembly. Therefore, we ran Falsifier four times to create training and test data sets, with each data set including one assembly at a 0.87% misassembly rate and one at a 3.32% rate.

Here is the command we ran for creating falsified assemblies with Falsifier:

```
Docker run -it -v${WORKING_DIR}:${WORKING_DIR} --rm  mobinasri/flagger:v1.1.0 \
    python3 /home/programs/src/falsifier.py \
    --hap1 ${PAT_T2T_FASTA} \
    --hap2 ${MAT_T2T_FASTA} \
    --paf ${MONTAGE_MAPPER_PAF} \
    --outputDir ${OUTPUT_DIRECTORY} \
    --minAlignmentLength 20000 \
    --misAssemblyTsv ${MISASSEMBLY_SIZE_TABLE_TSV} \
    --misjoinJson ${MISJOIN_COUNT_JSON}  \
    --marginLength 5000 \
    --annotationsJson ${ANNOTATIONS_PATH_JSON} \
    --singleBaseErrorRate 0.04 \
```

```
--maxGapLength 500 \
--outPrefix ${OUTPUT_PREFIX}
```

The input files for running this command with misassembly rates of 3.32% and 0.87% are available in this s3 area:

https://s3-us-west-2.amazonaws.com/human-pangenomics/index.html?prefix=submissions/e093fd72-e31a-11ee-b020-27964ee37032--flagger_test_files/misassembly_simulation_oct_2024/

The four falsified assemblies created for training and test data sets are available in this s3 area:

https://s3-us-west-2.amazonaws.com/human-pangenomics/index.html?prefix=submissions/e093fd72-e31a-11ee-b020-27964ee37032--flagger_test_files/misassembly_simulation_oct_2024/output_falsified_assemblies/

| **length_kb** | **active_hor** | | | **hsats** | | | **sd_no_hor_or_hsat** | | | **other** | | |
|---|---|---|---|---|---|---|---|---|---|---|---|---|
| | Err | Dup | Col | Err | Dup | Col | Err | Dup | Col | Err | Dup | Col |
| 40 | 4 | 4 | 4 | 4 | 4 | 4 | 30 | 30 | 30 | 30 | 30 | 30 |
| 80 | 4 | 4 | 0 | 4 | 4 | 4 | 15 | 15 | 15 | 15 | 15 | 15 |
| 160 | 0 | 0 | 0 | 0 | 0 | 0 | 4 | 4 | 4 | 8 | 8 | 8 |
| 320 | 0 | 0 | 0 | 0 | 0 | 0 | 4 | 4 | 4 | 8 | 8 | 8 |

**Table 1** Misassembly counts per annotation, block length and misassembly type for creating assemblies with 0.87% misassembly rate

| **length_kb** | **active_hor** | | | **hsats** | | | **sd_no_hor_or_hsat** | | | **other** | | |
|---|---|---|---|---|---|---|---|---|---|---|---|---|
| | Err | Dup | Col | Err | Dup | Col | Err | Dup | Col | Err | Dup | Col |
| 40 | 15 | 15 | 5 | 15 | 15 | 15 | 120 | 120 | 120 | 120 | 120 | 120 |
| 80 | 15 | 15 | 0 | 15 | 15 | 15 | 60 | 60 | 60 | 60 | 60 | 60 |
| 160 | 0 | 0 | 0 | 0 | 0 | 0 | 30 | 30 | 30 | 30 | 30 | 30 |
| 320 | 0 | 0 | 0 | 0 | 0 | 0 | 15 | 15 | 15 | 15 | 15 | 15 |

**Table 2** Misassembly counts per annotation, block length and misassembly type for creating assemblies with 3.32% misassembly rate

### 8 Measuring HMM-Flagger performance using three types of metric

To compare HMM-Flagger predictions against the truth labels, we used three different types of metrics, which are named base-level, overlap-based and auN-based metrics.

For computing the base-level metric we fill a confusion matrix by counting the number of bases with any pair of prediction and truth labels (**Supp Fig 41**). This confusion matrix is then used to compute precision and recall rates per label.

A limitation of the base-level metric is that it is more influenced by longer misassemblies than shorter ones and does not account for the number of distinct events detected by HMM-Flagger (**Supp Fig 42**). To address the issue with the base-level metric we used an overlap-based metric, for which we take the contiguous blocks with the same truth label and compute what percentage of each block is predicted as any of the four labels. If the percentage was higher than a threshold (40% as default) that counts as a hit and we will add one unit to the related entry in the confusion matrix (**Supp Fig 43**). Using this metric we created two confusion matrices; one by counting hits with respect to the contiguous blocks with the same truth label and one with respect to the blocks with the same prediction label (**Supp Fig 44**). These two matrices are not necessarily the same. We used the truth-based matrix for computing recall rates and the prediction-based one for precision.

The overlap-based and base-level metrics do not consider how the lengths of the events detected by HMM-Flagger correlate with the lengths of the truth blocks (**Supp Fig 45**). We utilized the auN score, developed originally for assessing assembly contiguity, in our third metric to measure the contiguity of the correctly predicted blocks. For the auN-based metric we compute auN using the blocks with correct predictions and then normalize it by the auN score of the truth blocks. This provides a score between 0 and 1, where values closer to 1 indicate that the contiguity of the truth blocks is better preserved in the predicted blocks (**Supp Fig 46**).

### 9 Code and data availability

#### 9.1 HMM-Flagger code and workflows

All HMM-Flagger analyses in this study were performed using the Docker image `mobinasri/flagger:v1.2.0`. Most analyses were executed via a WDL-based workflow on the SLURM cluster at the University of California, Santa Cruz, using the Toil workflow engine [8]. The primary workflow used is available at: https://github.com/mobinasri/flagger/blob/main/wdls/workflows/hmm_flagger_end_to_end_with_mapping.wdl.

Bash scripts, Jupyter notebooks, and data tables used to run minimap2 and HMM-Flagger on HPRC assemblies were executed in multiple batches and are available at the following locations:

- https://github.com/human-pangenomics/hprc_intermediate_assembly/tree/main/assembly_qc/batch1/hmm_flagger
- https://github.com/human-pangenomics/hprc_intermediate_assembly/tree/main/assembly_qc/batch1_jan_12_2025/hmm_flagger
- https://github.com/human-pangenomics/hprc_intermediate_assembly/tree/main/assembly_qc/batch2/hmm_flagger
- https://github.com/human-pangenomics/hprc_intermediate_assembly/tree/main/assembly_qc/batch3/hmm_flagger

Additional documentation and instructions for running HMM-Flagger workflows are available at: https://github.com/mobinasri/flagger.

#### 9.2 Genome assemblies for HPRC release 2

Links to all genome assemblies included in HPRC release 2 are provided in the following index table: https://github.com/human-pangenomics/hprc_intermediate_assembly/blob/main/data_tables/assemblies_release2_v1.0.index.csv.

#### 9.3 Read alignments for HPRC release 2

HiFi and Oxford Nanopore Technologies (ONT) read alignments used for HMM-Flagger analyses of HPRC release 2 assemblies are available from the following S3 repository: https://s3-us-west-2.amazonaws.com/human-pangenomics/index.html?prefix=submissions/ca366a13-5bad-487b-8a57-97344e9aa0e4--HPRC_RELEASE_2_SUPPLEMENTARY_ASSEMBLY_QC/.

For each sample, alignments are organized by sequencing technology (HiFi or ONT). For example, the HiFi and ONT BAM files for sample HG00097 are located under:

- `HG00097/hprc_r2/assembly_qc/read_alignments/hifi/`
- `HG00097/hprc_r2/assembly_qc/read_alignments/ont/`

ONT data for some samples were generated using R9 chemistry, whereas others were generated using R10 chemistry.

##### 9.4 CenSat annotations

Centromeric satellite (CenSat) BED files for HPRC release 2 were generated using the AlphaAnnotation workflow and are indexed at: https://github.com/human-pangenomics/hprc_intermediate_assembly/blob/main/data_tables/annotation/censat/censat_hprc_r2_v1.0.index.csv.

##### 9.5 HMM-Flagger output BED files

HMM-Flagger (v1.2.0) output BED files for HPRC release 2 assemblies are available in the same S3 repository as the read alignments: https://s3-us-west-2.amazonaws.com/human-pangenomics/index.html?prefix=submissions/ca366a13-5bad-487b-8a57-97344e9aa0e4--HPRC_RELEASE_2_SUPPLEMENTARY_ASSEMBLY_QC/.

For each sample, unfiltered and conservative BED files are provided separately for HiFi- and ONT-based analyses. For example, the HMM-Flagger output files for sample HG00097 are located under:

- `HG00097/hprc_r2/assembly_qc/hmm_flagger/v1.2.0_hifi/`
- `HG00097/hprc_r2/assembly_qc/hmm_flagger/v1.2.0_ont/`
