## Supplementary Figures for "Evaluating genome assemblies with HMM-Flagger"

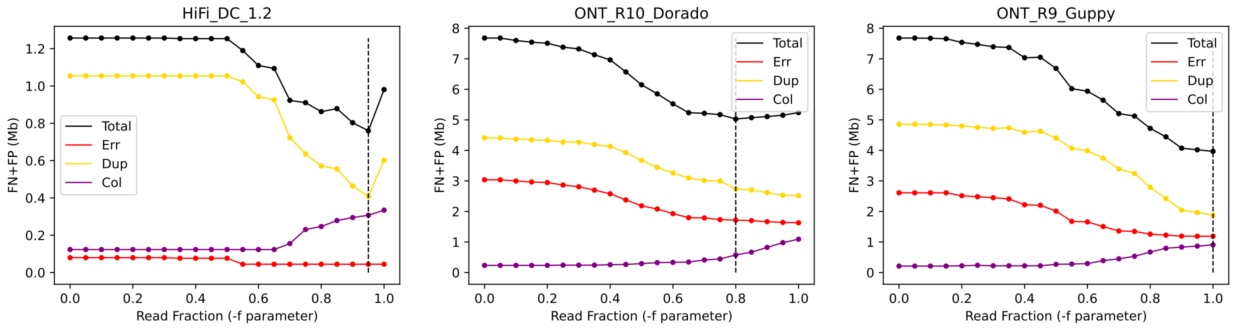


**Supplementary Fig 1 Tuning minimum read fraction for adjusting coverage expectation at contig ends.** Only the 50kb ends of contigs from the falsified assembly (3.32% misassembly rate) were included for tuning. The total number of misclassified bases were counted by sweeping the parameter. The tuned values for HiFi, ONT-R10 and ONT-R9 platforms were 0.95, 0.8 and 1.0 respectively.

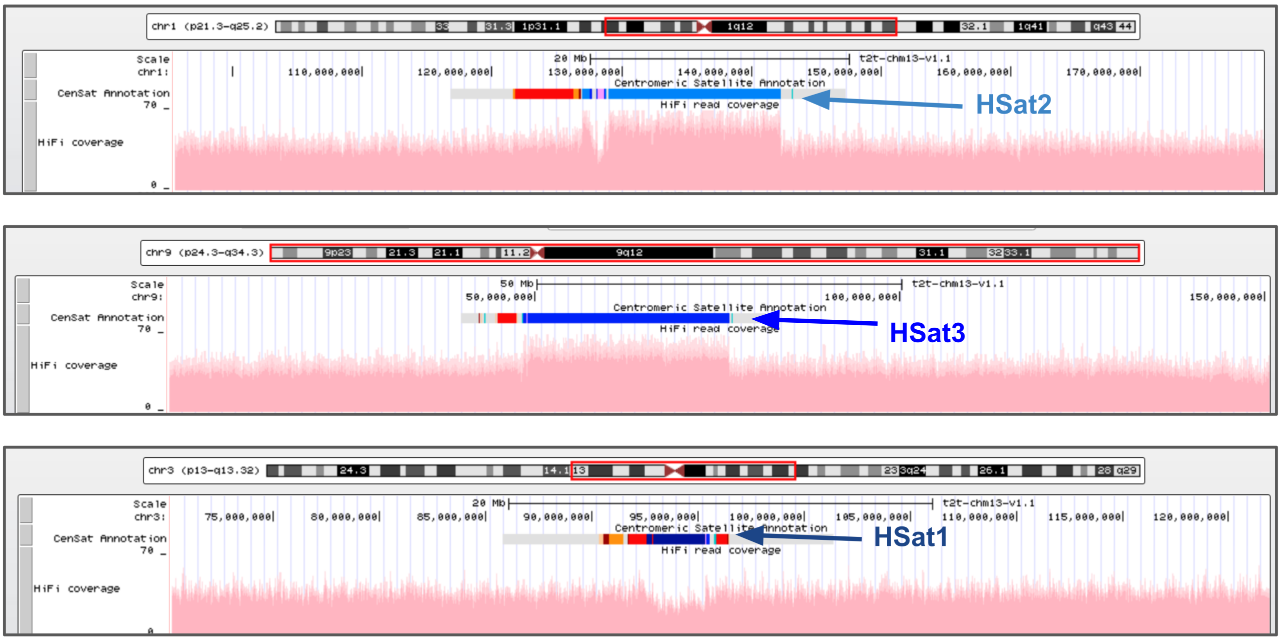


**Supplementary Fig 2 HiFi coverage biases in HSat regions:** The coverage of the HiFi reads aligned to T2T-CHM13v1.1 is available on the UCSC genome browser. The coverage in human satellites may systematically increase or decrease. For example, HSat2 in chr1 and HSat3 in chr9 (a and b) have about 1.25× higher coverage (25% increase), whereas HSat1 in chr3 (c) has about 0.75× the coverage (25% decrease) compared to the baseline.

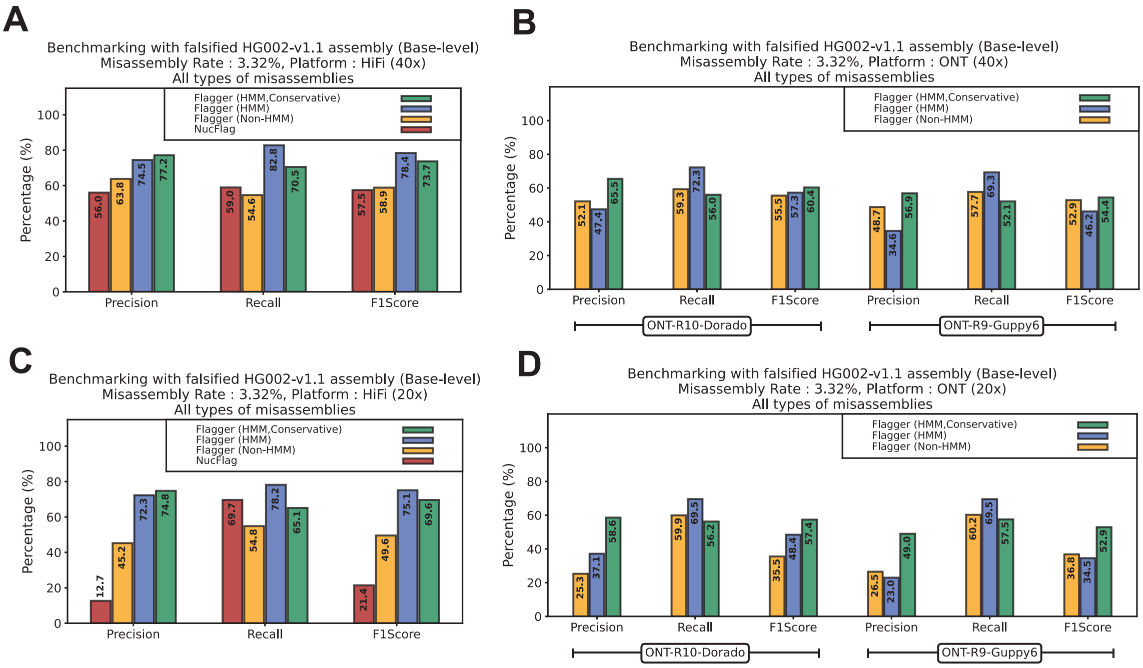


**Supplementary Fig 3 Benchmarking on falsified assemblies using base-level metric (3.32% misassembly rate): A)** For HiFi with 40x coverage **B)** For ONT R9 and R10 with 40x coverage **C)** For HiFi with 20x coverage **D)** For ONT R9 and R10 with 20x coverage

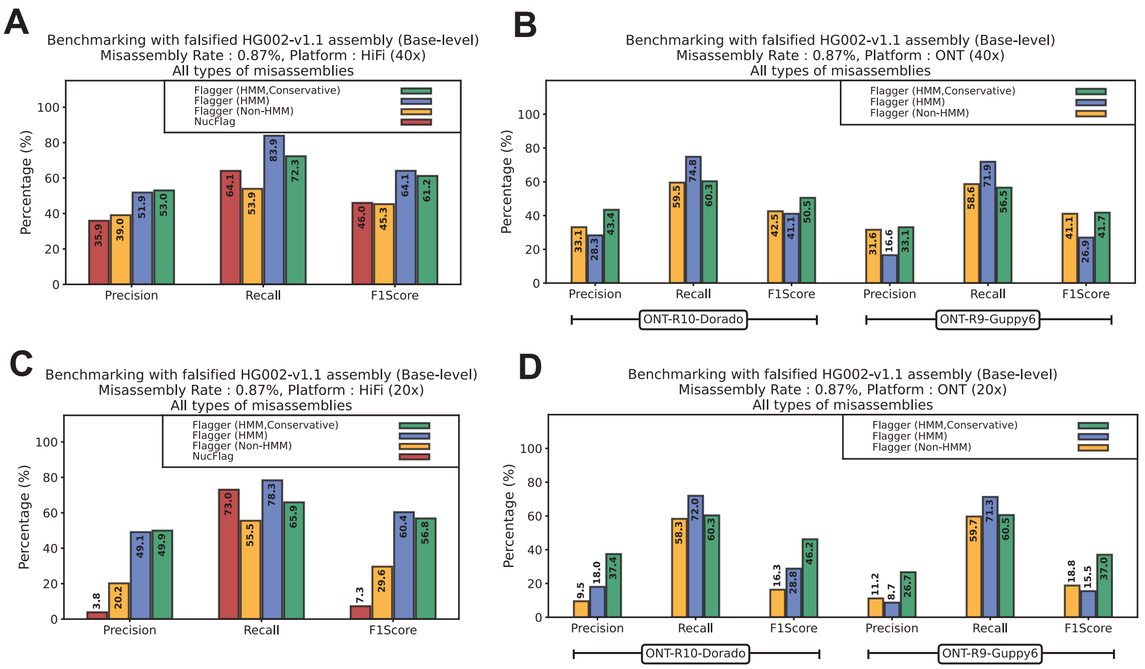


**Supplementary Fig 4 Benchmarking on falsified assemblies using base-level metric (0.87% misassembly rate): A)** For HiFi with 40x coverage **B)** For ONT R9 and R10 with 40x coverage **C)** For HiFi with 20x coverage **D)** For ONT R9 and R10 with 20x coverage

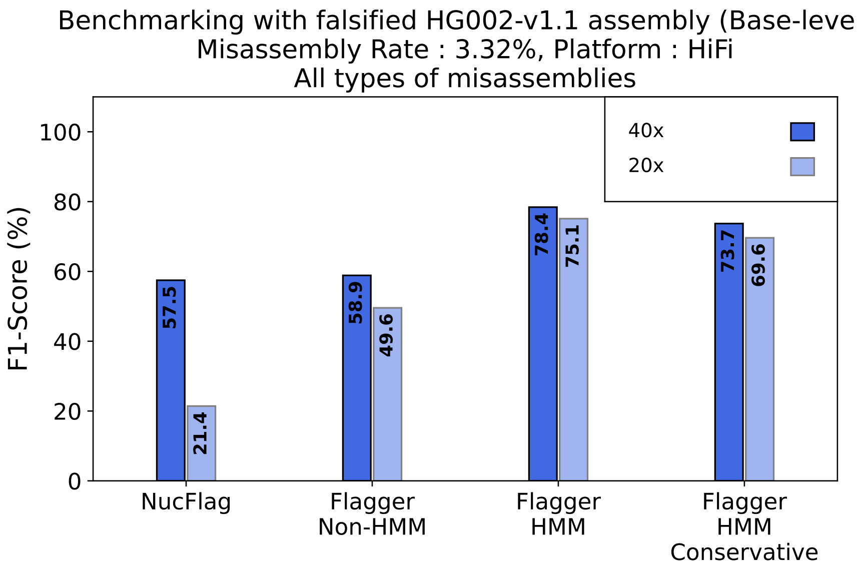


**Supplementary Fig 5 Impact of HiFi sequencing coverage on HMM-Flagger's performance**

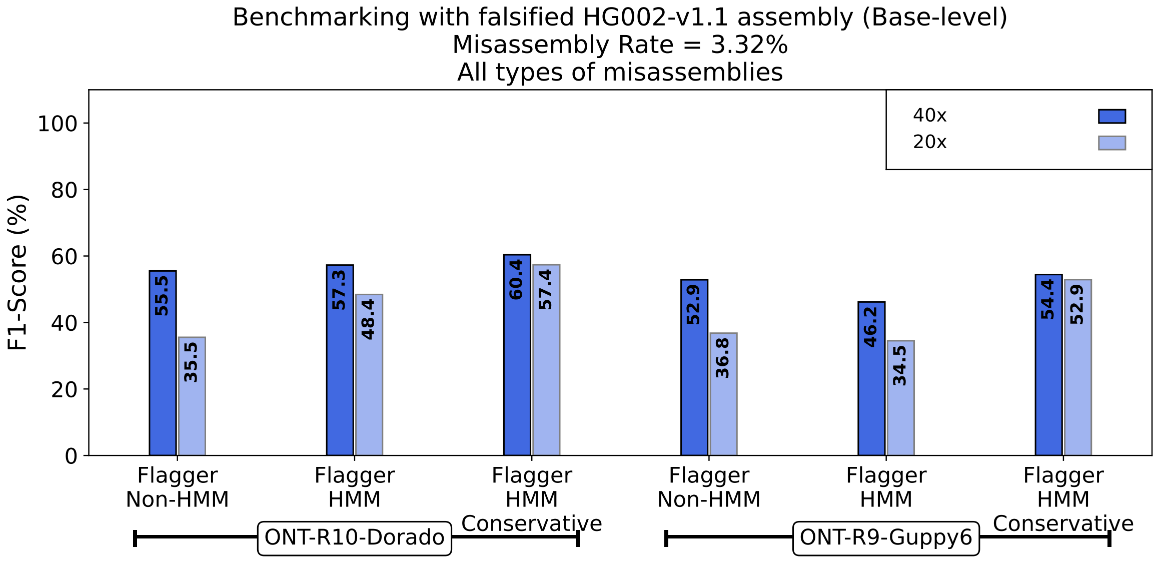


**Supplementary Fig 6 Impact of ONT sequencing coverage on HMM-Flagger's performance**

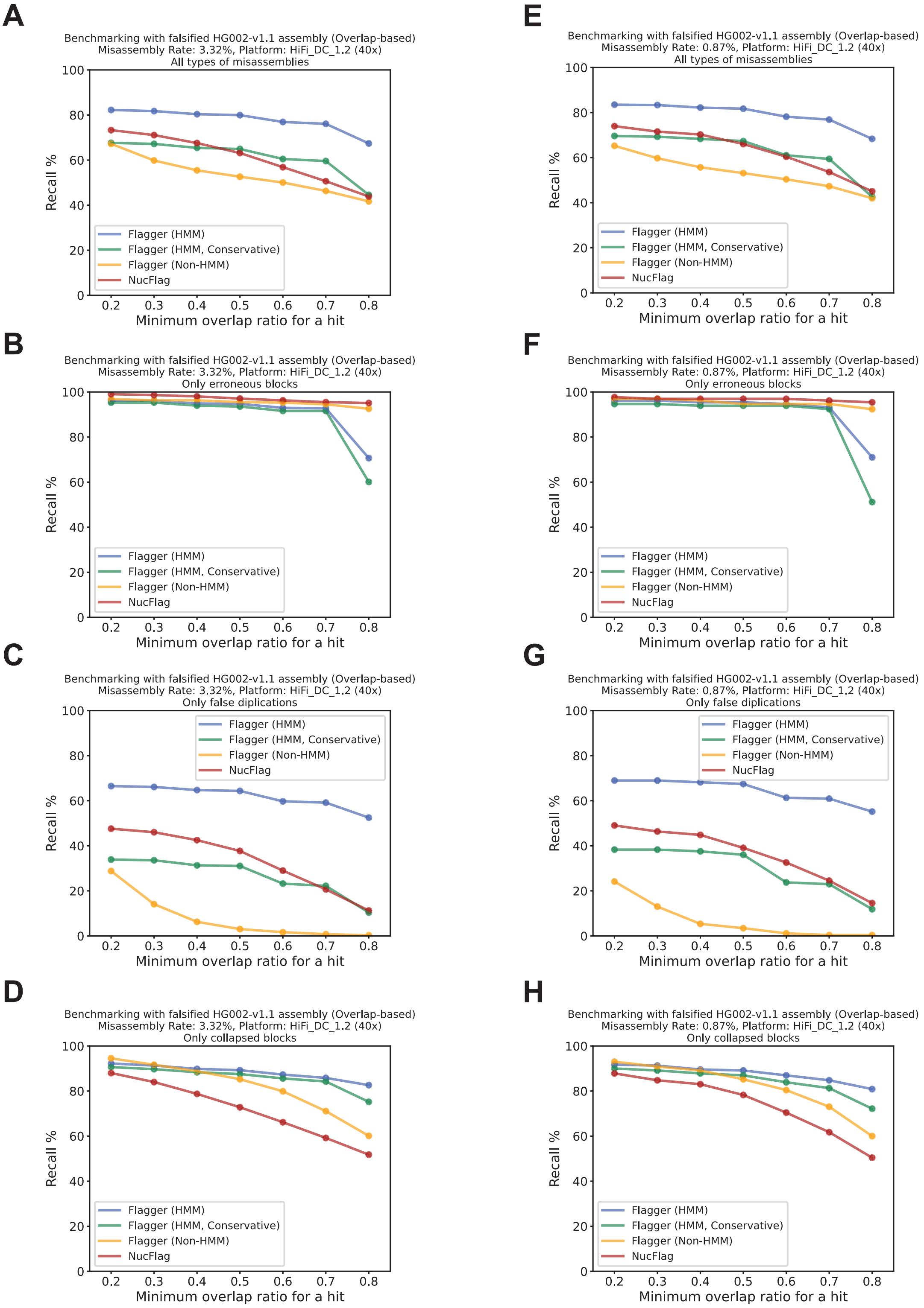


**Supplementary Fig 7 Benchmarking on falsified assemblies using HiFi data.** The y-axis shows the overlap-based recall rate, while the x-axis represents the minimum overlap ratio. **A–D)** Results for assemblies with a misassembly rate of 3.32%: (A) all misassembly types, (B) erroneous blocks, (C) false duplications, and (D) collapsed blocks. **E–H)** Results for assemblies with a misassembly rate of 0.87%: (E) all misassembly types, (F) erroneous blocks, (G) false duplications, and (H) collapsed blocks.

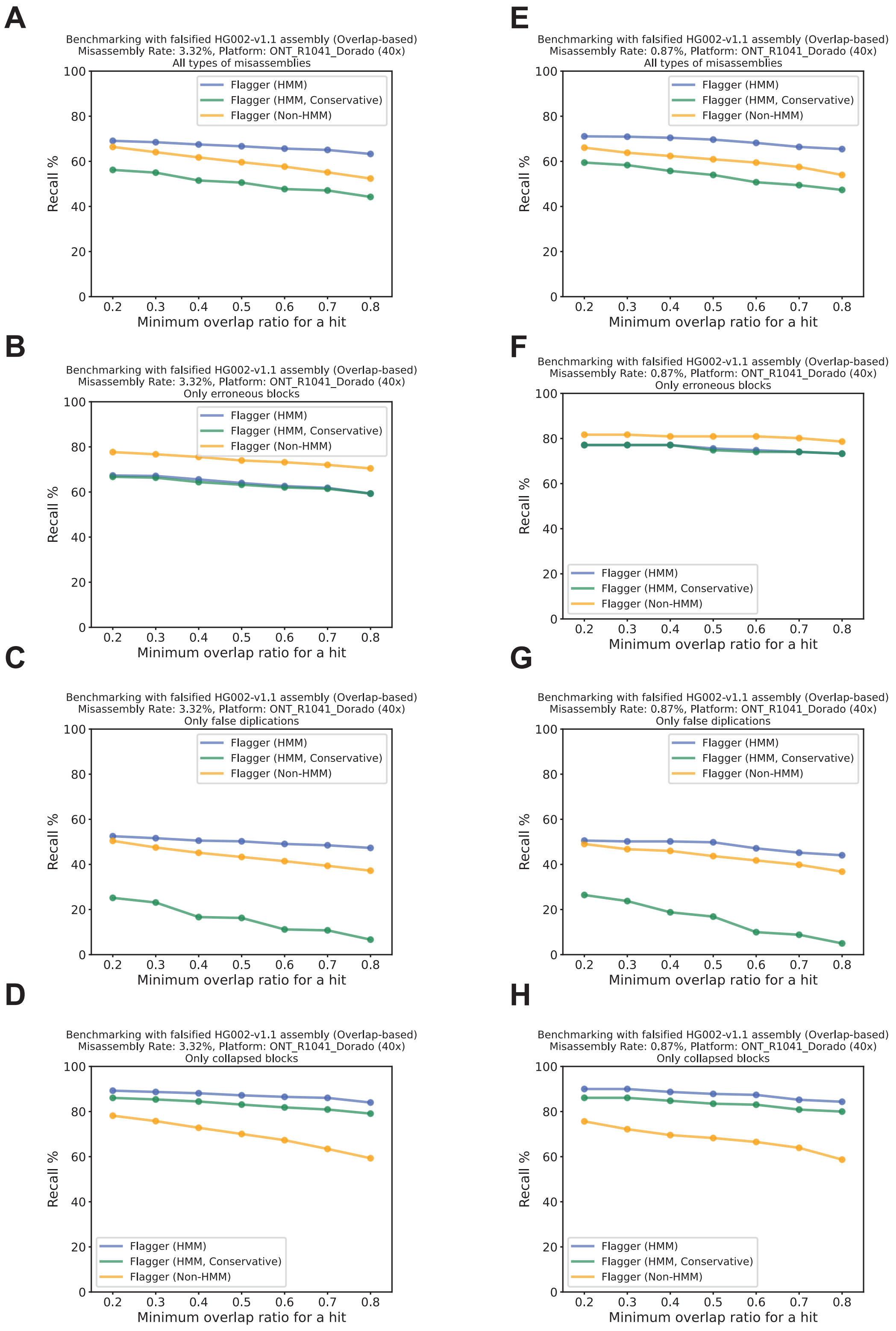


**Supplementary Fig 8 Benchmarking on falsified assemblies using ONT-R10 data.** The y-axis shows the overlap-based recall rate, while the x-axis represents the minimum overlap ratio. **A–D)** Results for assemblies with a misassembly rate of 3.32%: (A) all misassembly types, (B) erroneous blocks, (C) false duplications, and (D) collapsed blocks. **E–H)** Results for assemblies with a misassembly rate of 0.87%: (E) all misassembly types, (F) erroneous blocks, (G) false duplications, and (H) collapsed blocks.

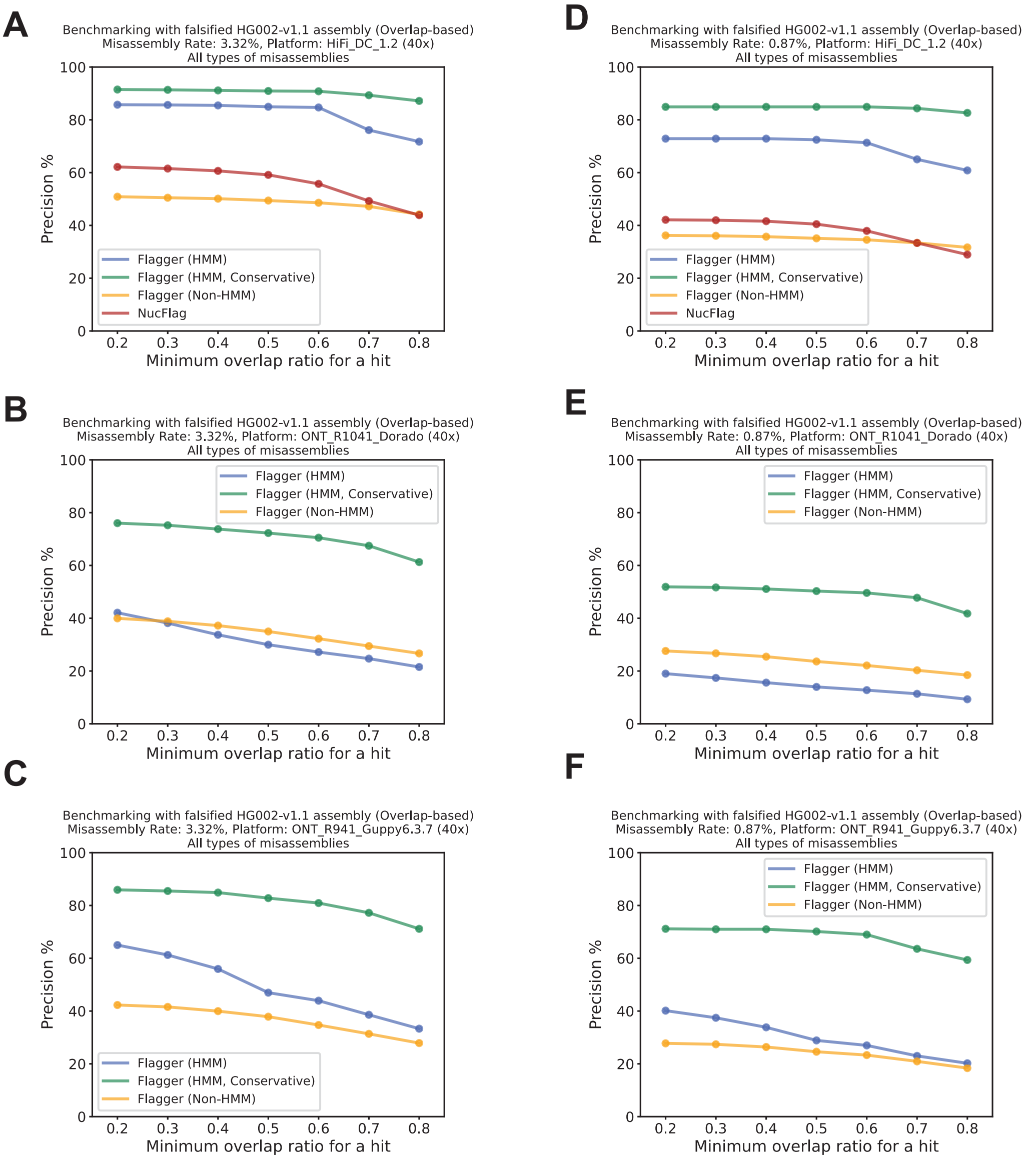


**Supplementary Fig 9 Overlap-based precision rates on different platforms and misassembly rates** The y-axis shows the overlap-based precision rate, while the x-axis represents the minimum overlap ratio. **A–C)** Results for assemblies with a misassembly rate of 3.32%: (A) HiFi, (B) ONT-R10, and (C) ONT-R9, **D–F)** Results for assemblies with a misassembly rate of 0.87%: (D) HiFi, (E) ONT-R10, and (F) ONT-R9.

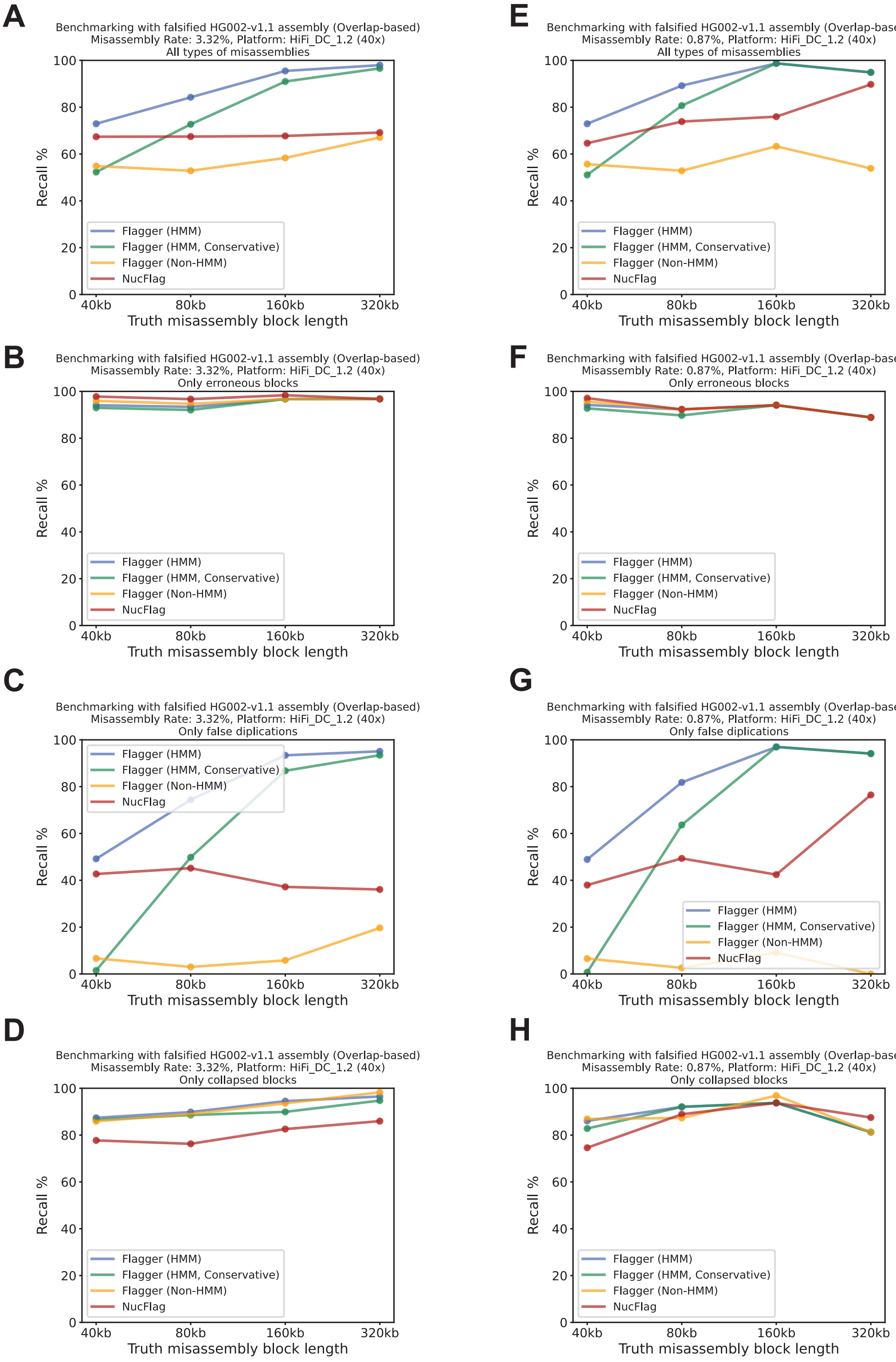


**Supplementary Fig 10 Impact of misassembly size on recall rate using HiFi data.** The y-axis shows the overlap-based recall rate, while the x-axis represents the misassembly size. **A–D)** Results for assemblies with a misassembly rate of 3.32%: (A) all misassembly types, (B) erroneous blocks, (C) false duplications, and (D) collapsed blocks. **E–H)** Results for assemblies with a misassembly rate of 0.87%: (E) all misassembly types, (F) erroneous blocks, (G) false duplications, and (H) collapsed blocks.

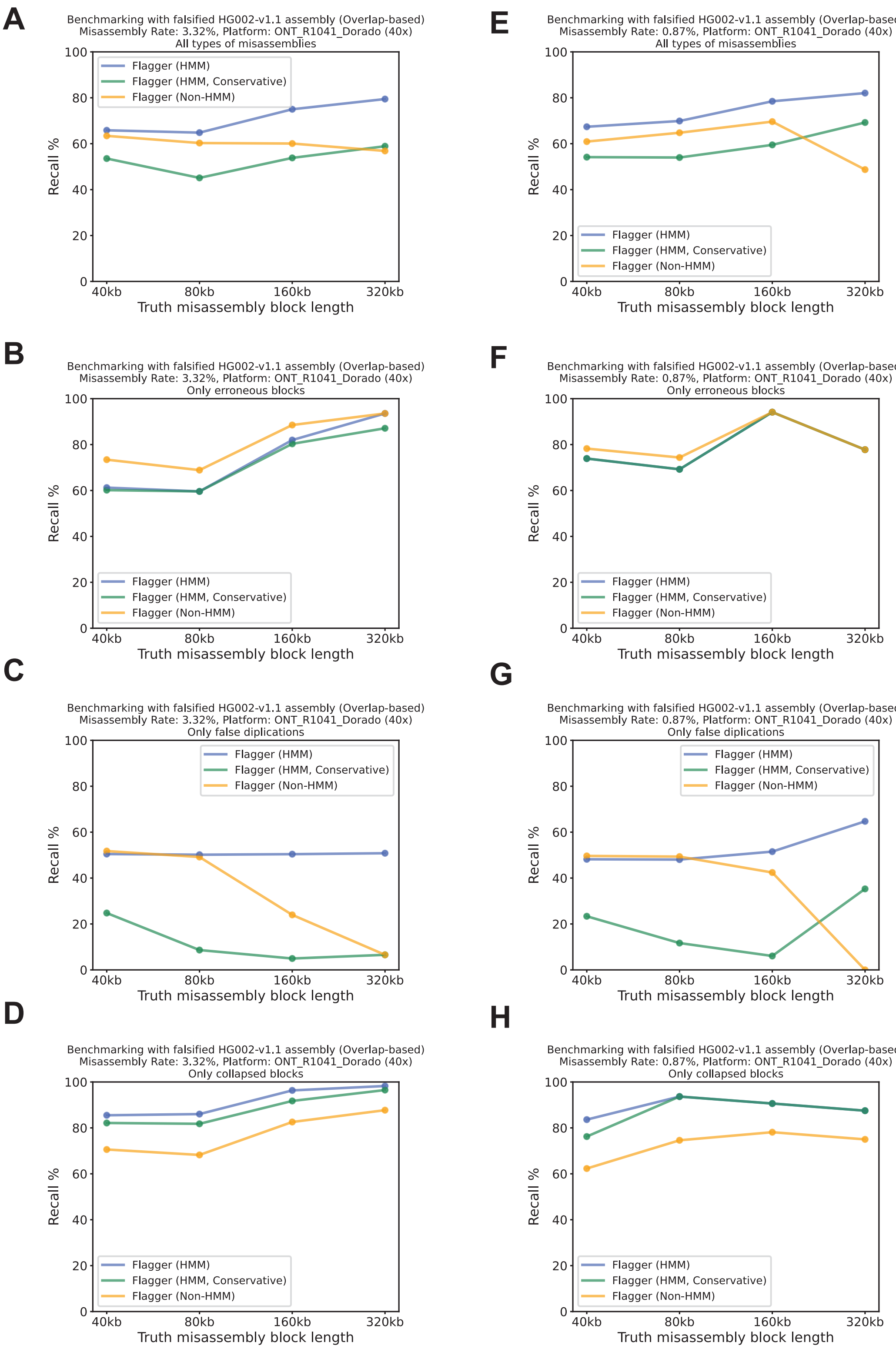


**Supplementary Fig 11 Impact of misassembly size on recall rate using ONT-R10 data.** The y-axis shows the overlap-based recall rate, while the x-axis represents the misassembly size. **A–D)** Results for assemblies with a misassembly rate of 3.32%: (A) all misassembly types, (B) erroneous blocks, (C) false duplications, and (D) collapsed blocks. **E–H)** Results for assemblies with a misassembly rate of 0.87%: (E) all misassembly types, (F) erroneous blocks, (G) false duplications, and (H) collapsed blocks.

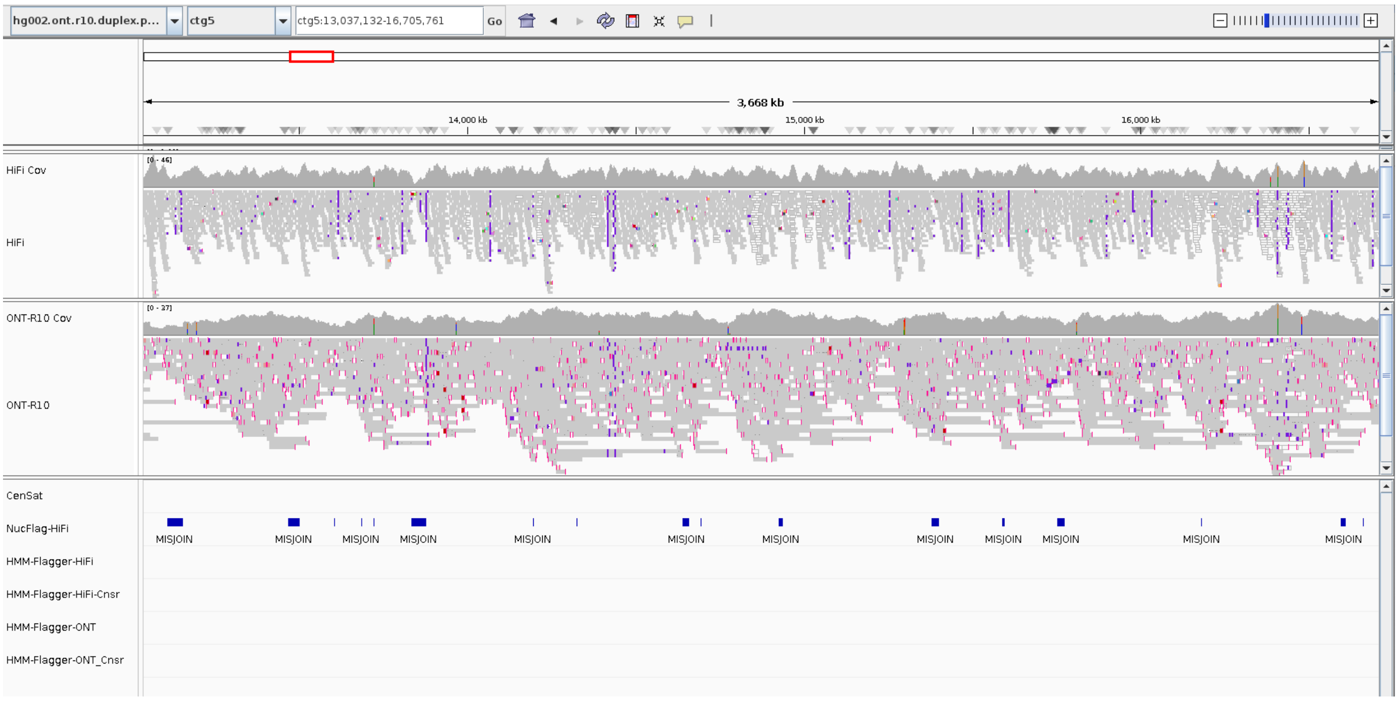


**Supplementary Fig 12 False Positive flags in NucFlag's output for Duplex-based PECAT assembly**

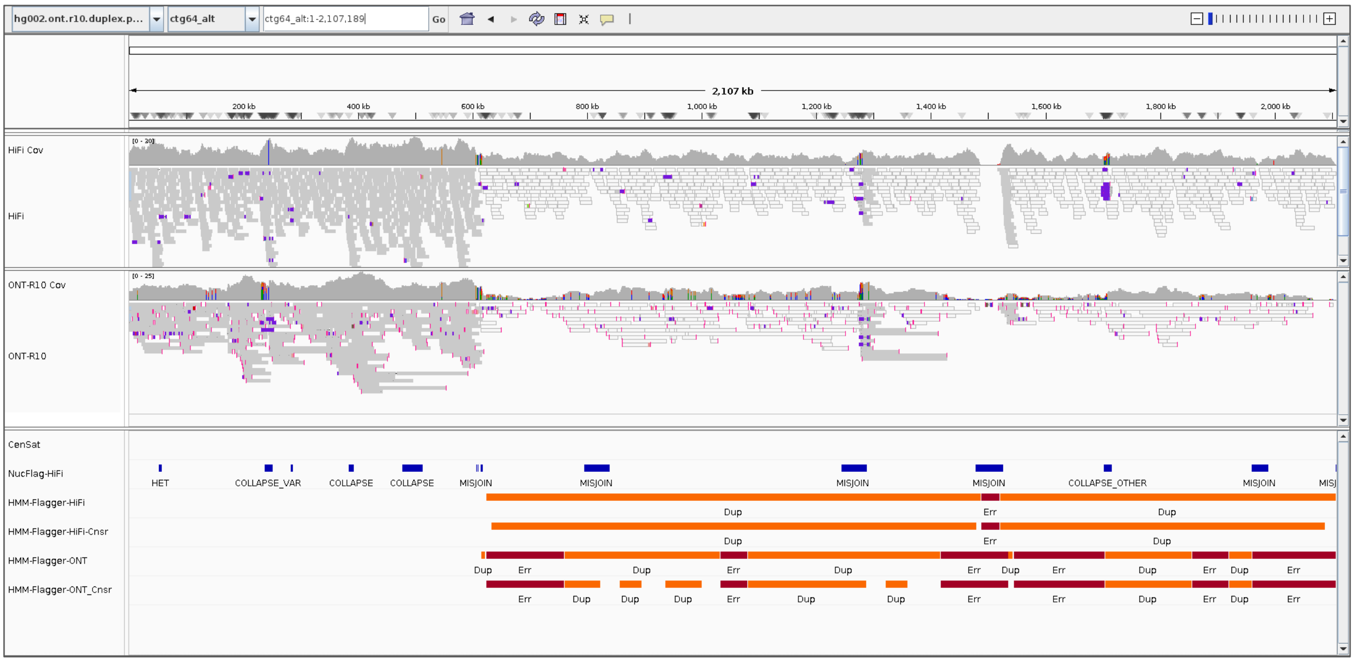


**Supplementary Fig 13 False duplication detected in the Duplex-based PECAT assembly:** A 1.5Mb region at the end of the contig ctg64_alt located in chr9-HSat3 array is detected as false duplication by HMM-Flagger. The mappings to T2T-v1.1 shown in **Supp Fig 14** confirmed this misassembly.

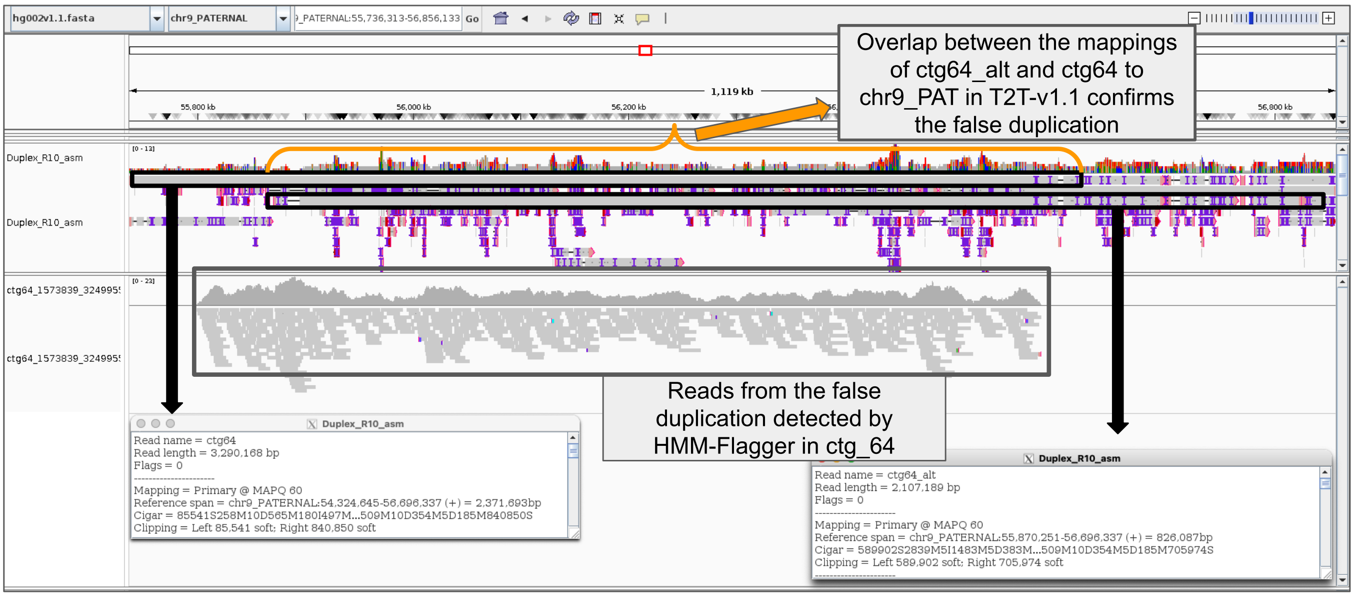


**Supplementary Fig 14 False duplications in the PECAT assembly mapped to T2T-v1.1:** Regions of about equal length (1.5Mb) at the ends of the contigs ctg64 and ctg64_alt were detected as false duplication by HMM-Flagger. The large overlap in the assembly mappings to chr9-PATERNAL of T2T-v1.1 confirmed this false duplication.

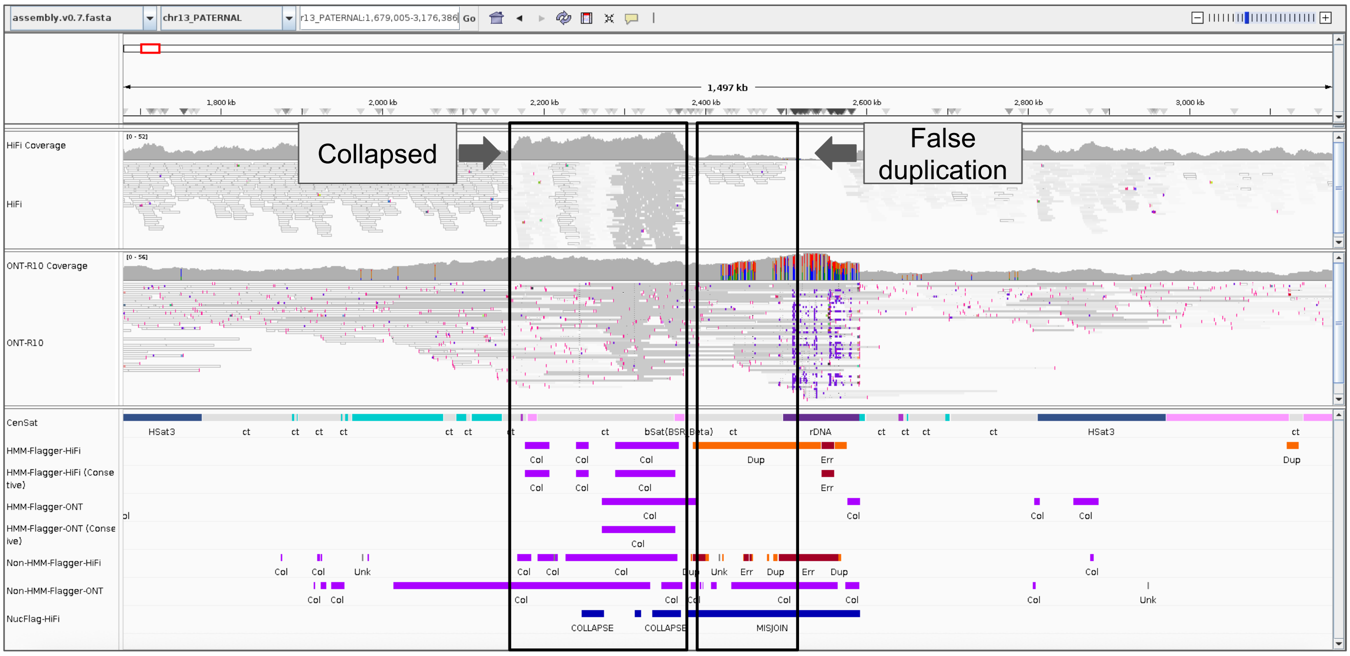


**Supplementary Fig 15 Two misassemblies next to each other detected in T2T-v0.7 assembly:** A region in chr13 PATERNAL (about 150 kb long) located downstream of rDNA array is detected as collapsed by HMM-Flagger using both HiFi and ONT-R10 data. It indirectly shows a sequence not assembled in chr22 PATERNAL. There is also a false duplication of about 150 kb right after the collapsed sequence that extends a bit into the rDNA array. The falsely duplicated copies of this block are assembled as small contigs. This false duplication (about 150 kb) was flagged only by using HiFi data.

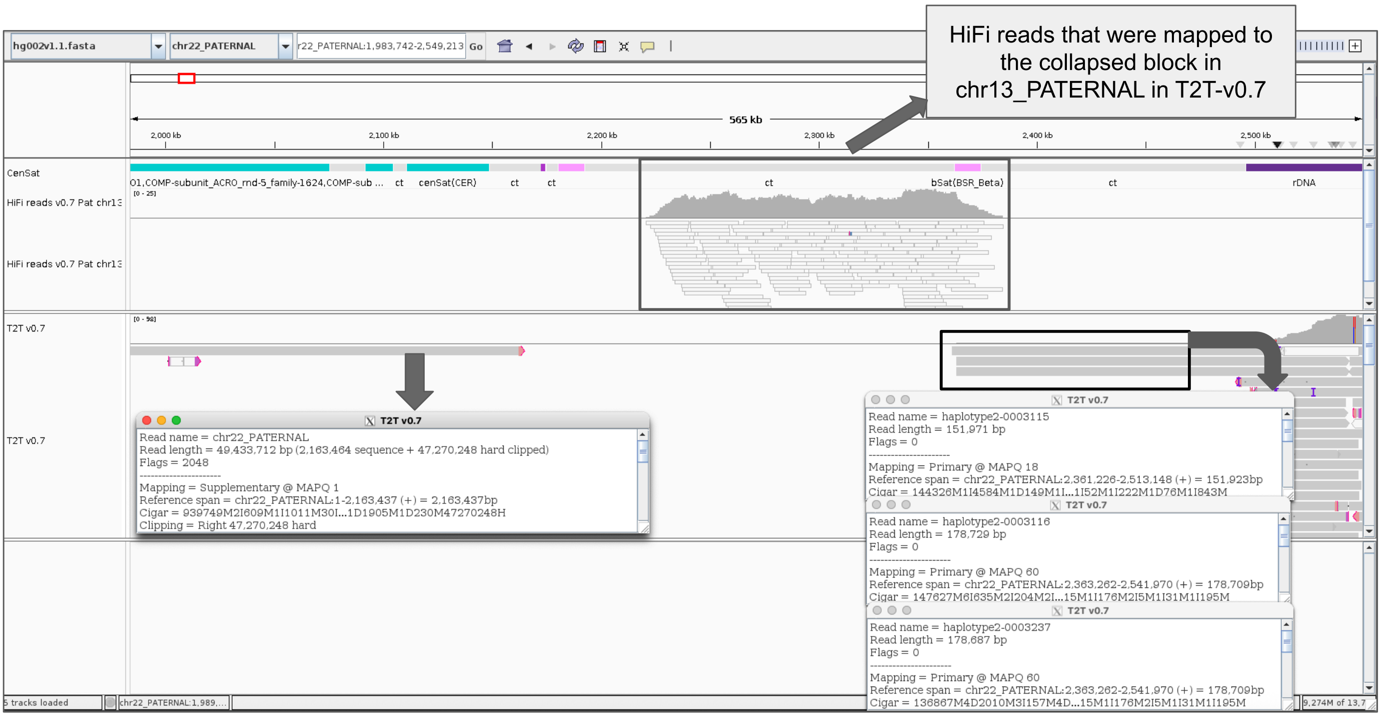


**Supplementary Fig 16 Mappings of T2T-v0.7 to T2T-v1.1 chr22 PATERNAL:** There is a missing sequence in T2T-v0.7 chr22 PATERNAL and that region does not have any mapping from T2T-v0.7 to T2T-v1.1. Nearly half of the HiFi reads that were mapped to the collapsed block in chr13 in T2T-v0.7 are correctly mapped to chr22 in T2T-v1.1. At the right side of the missing sequence we have multiple contigs all representing the same part of the genome redundantly. This false duplication was detected by HMM-Flagger in **Supp Fig 15**

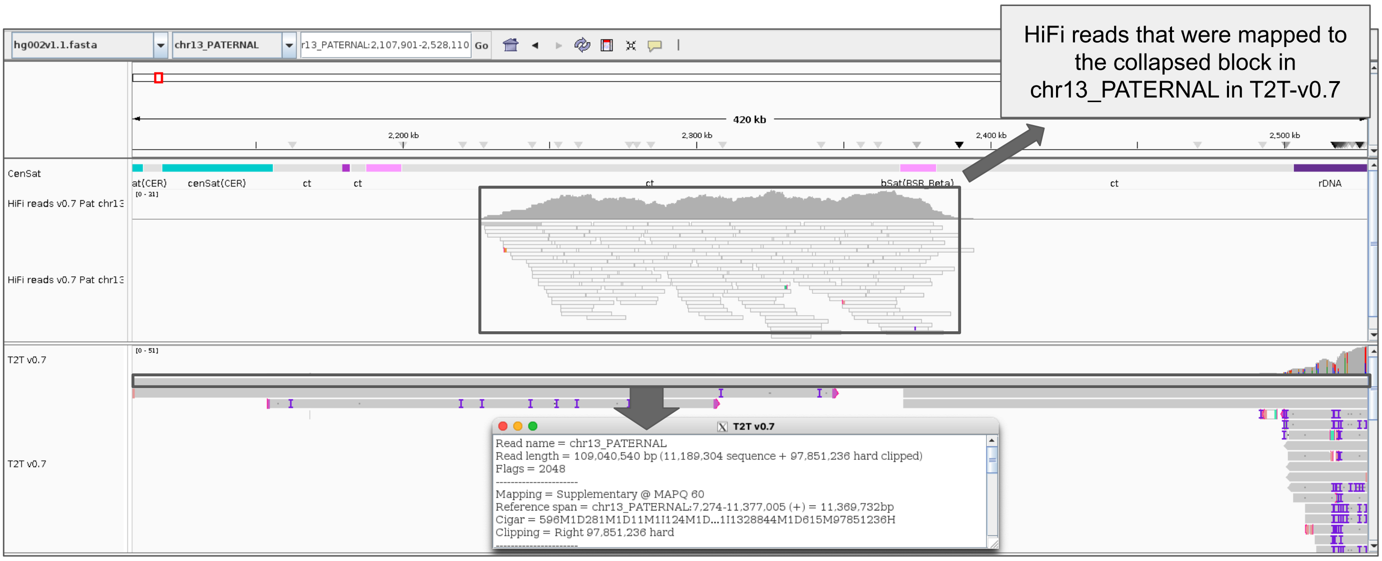


**Supplementary Fig 17 Mappings of T2T-v0.7 to T2T-v1.1 chr13 PATERNAL:** The region flagged as collapsed in chr13 PATERNAL of T2T-v0.7 assembly was mapped to T2T-v1.1. Nearly half of the HiFi reads from that collapsed block are mapped here. The other half of the reads are mapped to chr22 PATERNAL in T2T-v1.1 (Look at **Supp Fig 16**)

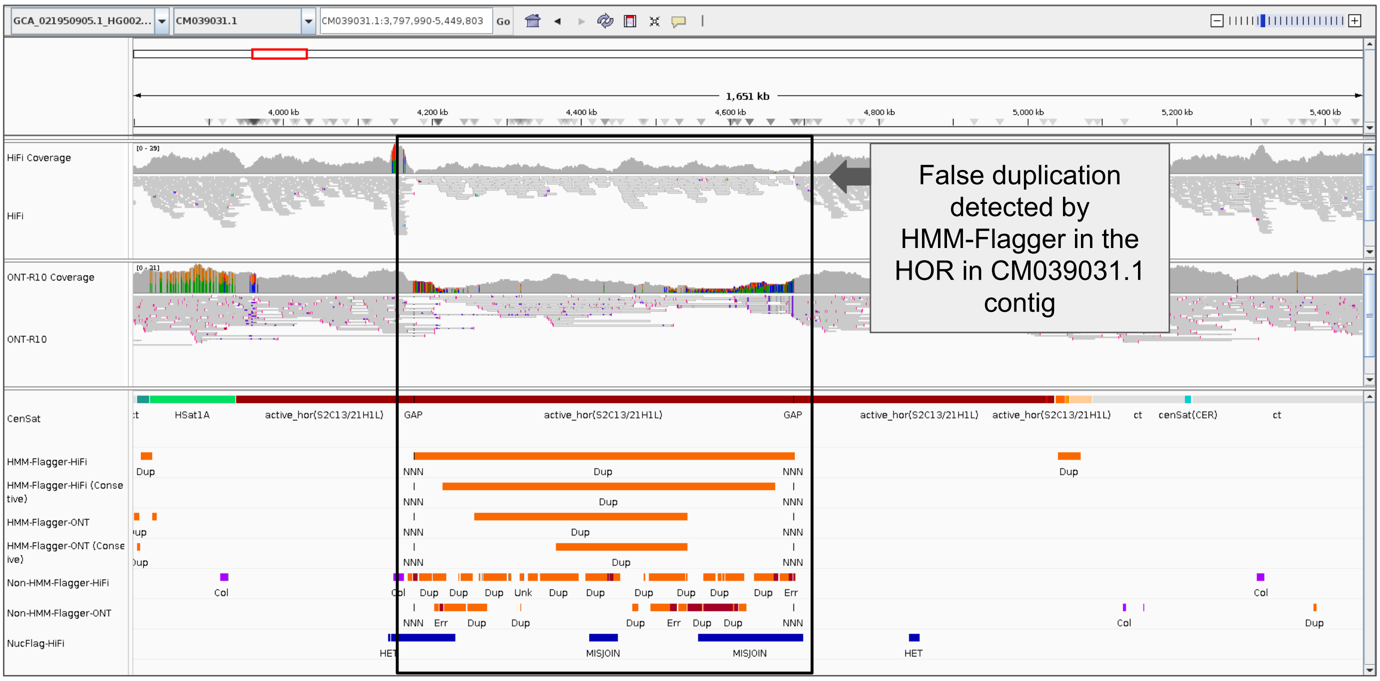


**Supplementary Fig 18  False duplication detected in an active HOR in the semi-automated assembly (Jarvis et al):** Further investigation showed that the other copy of the false duplication is located in an active HOR in the CM039046.1 contig **Supp Fig** 19

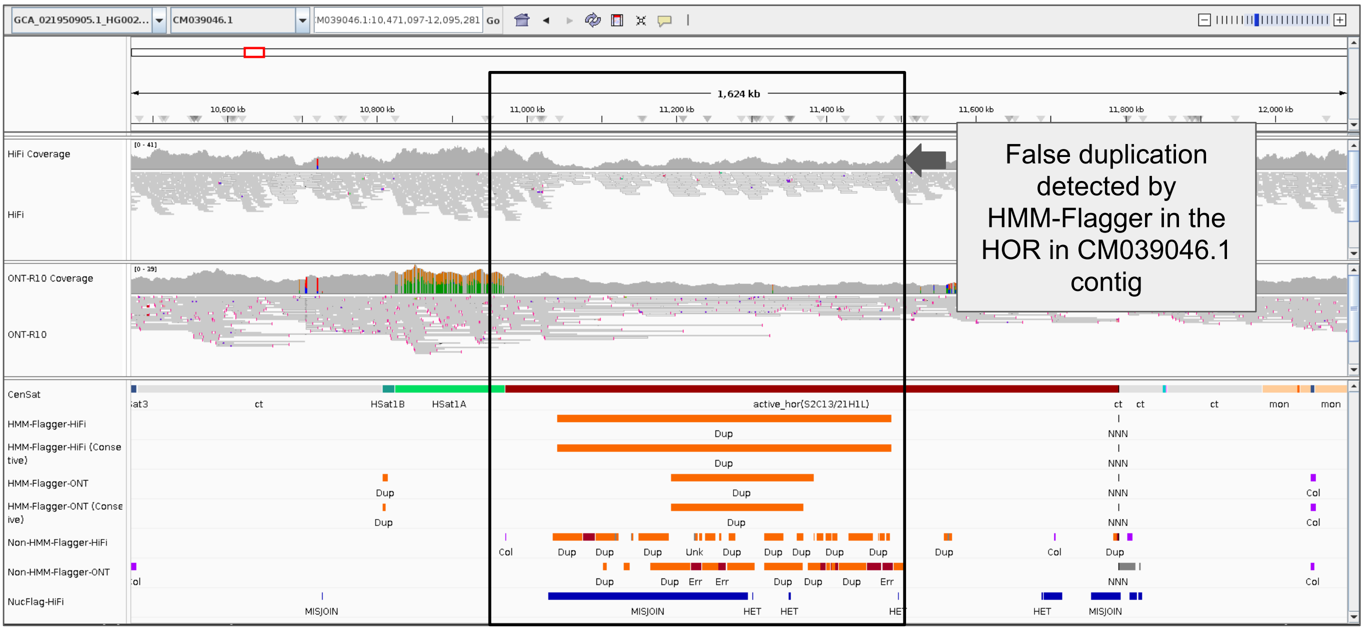


**Supplementary Fig 19  False duplication detected in an active HOR in the semi-automated assembly (Jarvis et al):** The other copy of the false duplication is located in an active HOR in the CM039031.1 contig. This is the redundant HOR assembled in the CM039046.1 contig as it is shown in **Supp Fig** 20. In the correct assembly this HOR belongs to chr21_PATERNAL (**Supp Fig** 21).

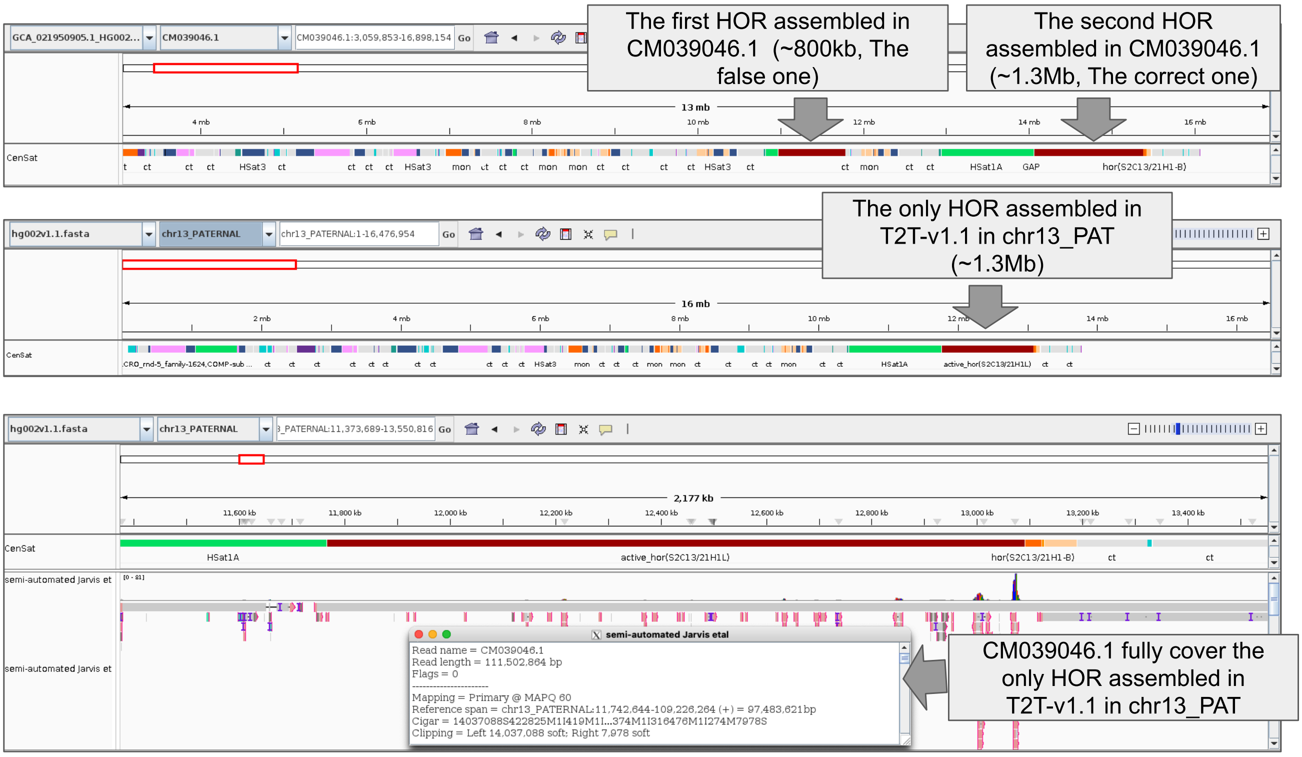


**Supplementary Fig 20 A redundant active HOR assembled in the CM039046.1 contig of the semi-automated assembly (Jarvis et al):** The top panel shows that in the CM039046.1 contig we have 2 assembled active HORs. This contig is mapped to chr13_PATERNAL in T2T-v1.1 and as it is shown in the middle panel this haplotype has only one active HOR. Only the second HOR (about 1.3 Mb) in CM039046.1 fully spans the correct array in T2T-v1.1. The first HOR (about 800 kb) was correctly flagged as problematic by HMM-Flagger as shown in **Supp Fig 19**

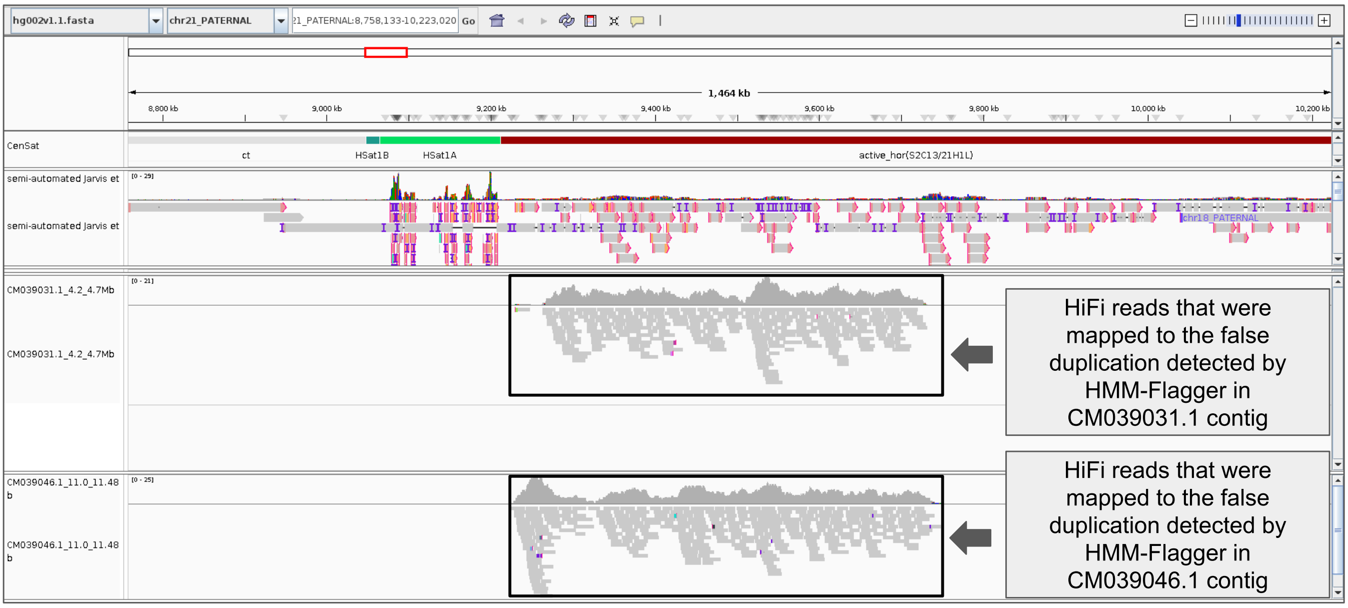


**Supplementary Fig 21 HiFi reads from the false duplications in the semi-automated assembly (Jarvis et al) are mapped to T2T-v1.1** The HiFi reads from the false duplications shown in **Supp Fig 18** and **Supp Fig 19** are mapped to T2T-v1.1, showing the origin of the duplicated copies in the correct assembly.

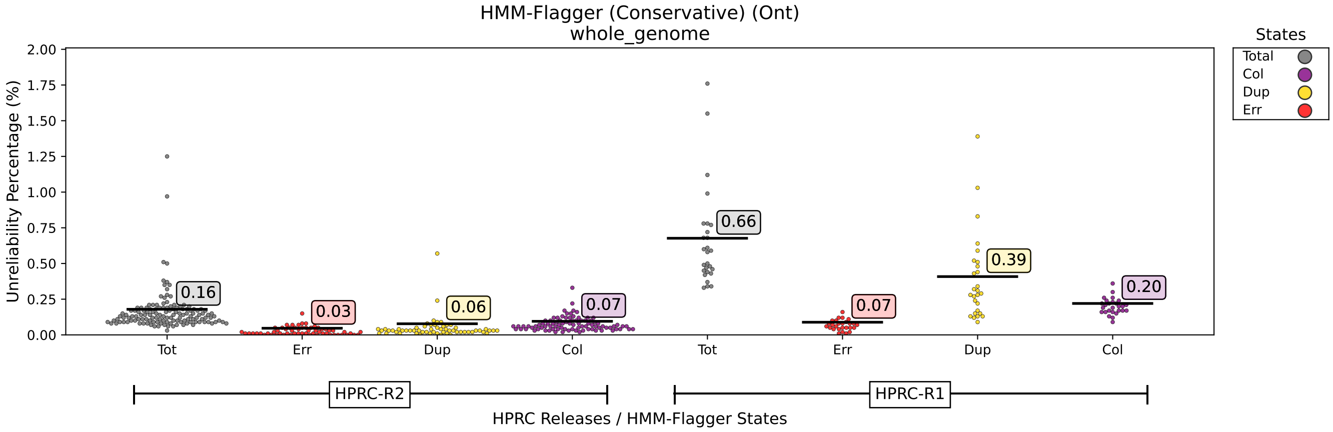


**Supplementary Fig 22 Comparing HPRC releases using ONT mappings and HMM-Flagger conservative sets (Whole genome)**

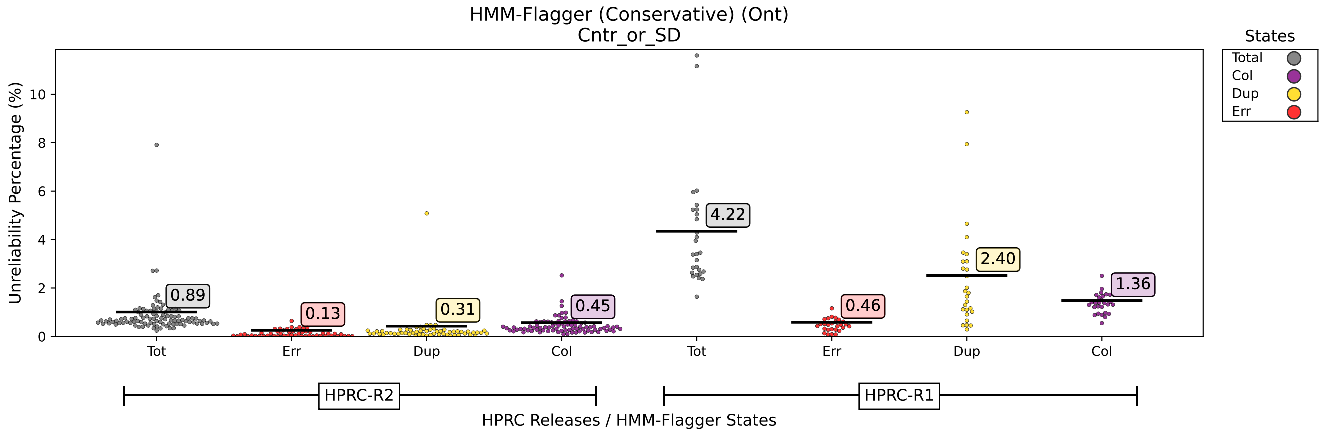


**Supplementary Fig 23** **Comparing HPRC releases using ONT mappings and HMM-Flagger conservative sets (Union of CenSat and SD)**

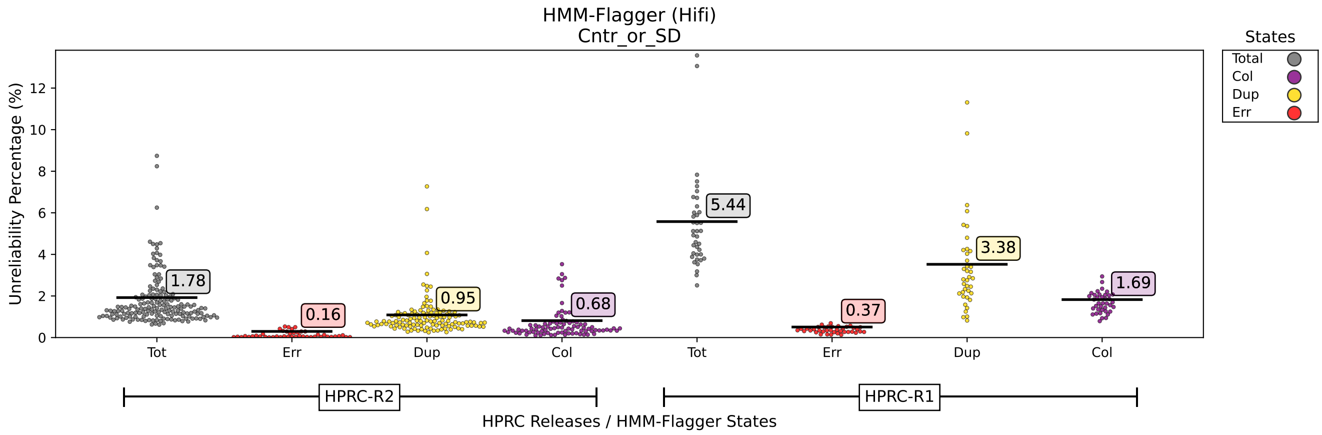


**Supplementary Fig 24** **Comparing HPRC releases using HiFi mappings and HMM-Flagger (Union of CenSat and SD)**

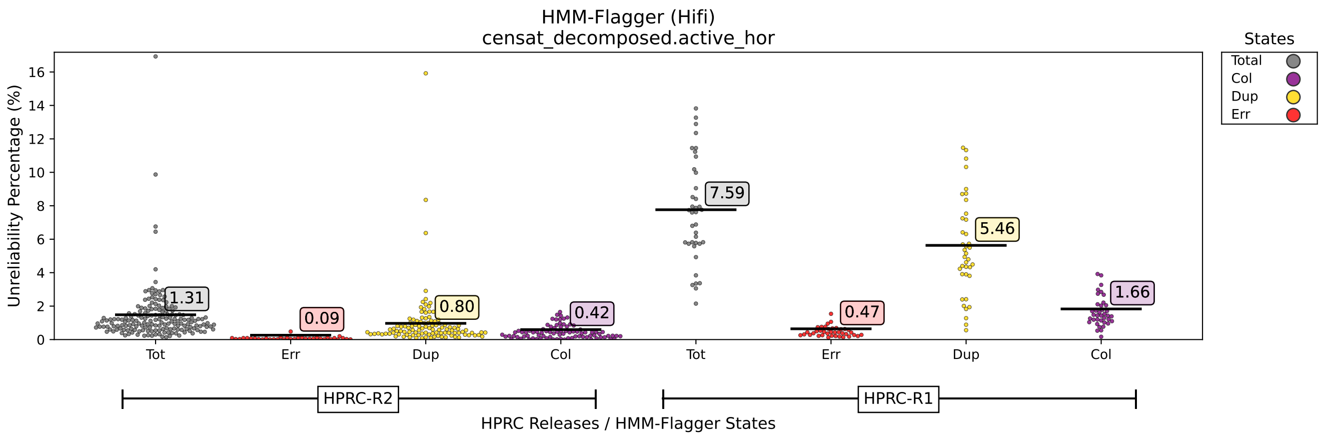


**Supplementary Fig 25** **Comparing HPRC releases using HiFi mappings and HMM-Flagger (active HOR)**

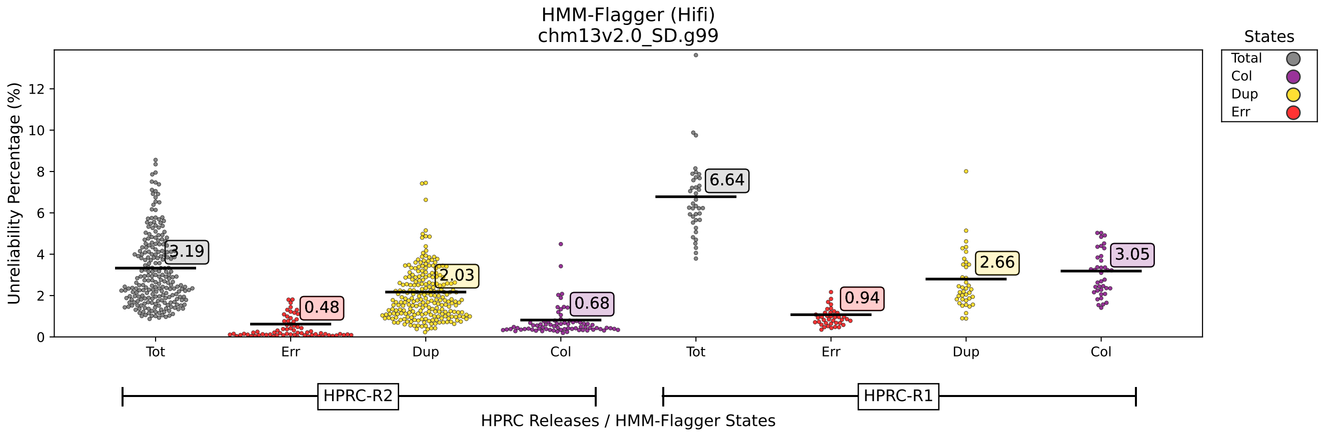


**Supplementary Fig 26** **Comparing HPRC releases using HiFi mappings and HMM-Flagger (SDs with similarity greater than 99%)**

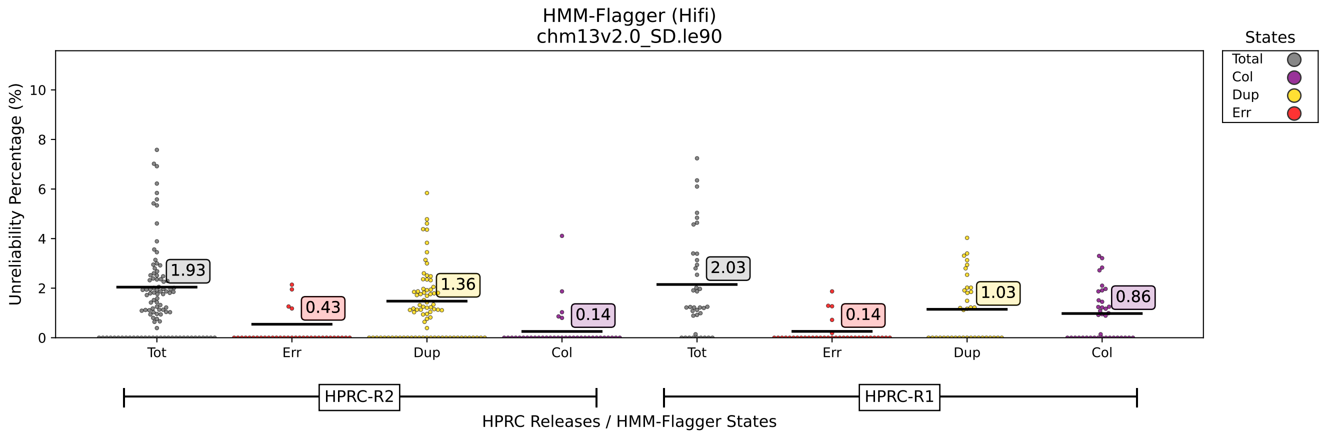


**Supplementary Fig 27** Comparing HPRC releases using HiFi mappings and HMM-Flagger (SDs with similarity lower than or equal to 90%)

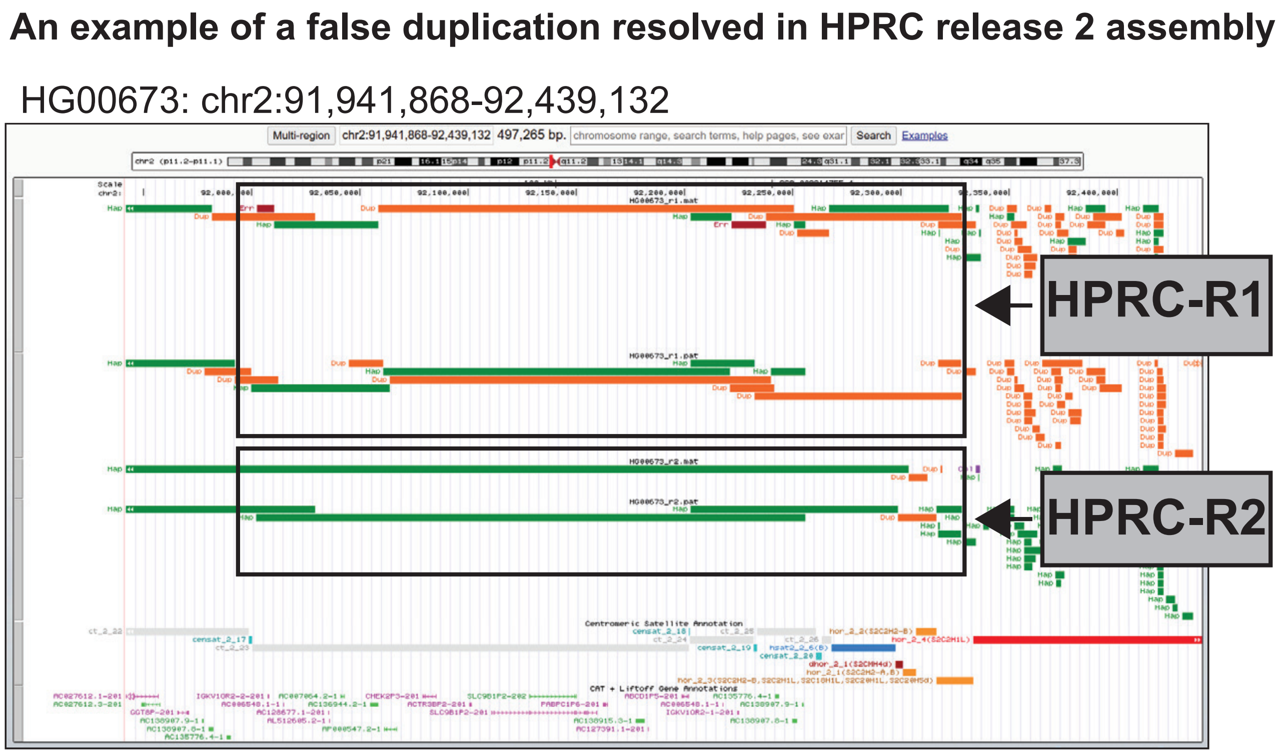


**Supplementary Fig 28** An example of a ∼300 kb false duplication detected upstream of the active HOR in chromosome 2. This false duplication was removed in HPRC-R2.

```
..
#
# Aligned_sequences: 2
# 1: NOTCH2NLC
# 2: NOTCH2NLC'
# Matrix: EDNAFULL
# Gap_penalty: 16
# Extend_penalty: 4
#
# Length: 819
# Identity:     818/819 (99.9%)
# Similarity:   818/819 (99.9%)
# Gaps:           0/819 ( 0.0%)
# Score: 4086
#
#
#=======================================

NOTCH2NLC          1 ATGCCCGCCCTGCGCCGCTCTGCTGTGGGCGCTGCTGGCGCTCTGGCTGT     50
                     ||||||||||||||||||||||||||||||||||||||||||||||||||
NOTCH2NLC'         1 ATGCCCGCCCTGCGCCGCTCTGCTGTGGGCGCTGCTGGCGCTCTGGCTGT     50

NOTCH2NLC         51 GCTGCGCGACCCCCGCGCATGGTGTCGAGATGGCTATGAACCCTGTGTAA    100
                     |||||||||||||||||||||.||||||||||||||||||||||||||||
NOTCH2NLC'        51 GCTGCGCGACCCCCGCGCATGTTGTCGAGATGGCTATGAACCCTGTGTAA    100

NOTCH2NLC        101 ATGAAGGAATGTGTGTTACCTACCACAATGGCACAGGATACTGCAAATGT    150
                     ||||||||||||||||||||||||||||||||||||||||||||||||||
NOTCH2NLC'       101 ATGAAGGAATGTGTGTTACCTACCACAATGGCACAGGATACTGCAAATGT    150

NOTCH2NLC        151 CCAGAAGGCTTCTTGGGGGAATATTGTCAACATCGAGACCCCTGTGAGAA    200
                     ||||||||||||||||||||||||||||||||||||||||||||||||||
NOTCH2NLC'       151 CCAGAAGGCTTCTTGGGGGAATATTGTCAACATCGAGACCCCTGTGAGAA    200

NOTCH2NLC        201 GAACCGCTGCCAGAATGGTGGGACTTGTGTGGCCCAGGCCATGCTGGGGA    250
                     ||||||||||||||||||||||||||||||||||||||||||||||||||
NOTCH2NLC'       201 GAACCGCTGCCAGAATGGTGGGACTTGTGTGGCCCAGGCCATGCTGGGGA    250

NOTCH2NLC        251 AAGCCACGTGCCGATGTGCCTCAGGGTTTACAGGAGAGGACTGCCAGTAC    300
                     ||||||||||||||||||||||||||||||||||||||||||||||||||
NOTCH2NLC'       251 AAGCCACGTGCCGATGTGCCTCAGGGTTTACAGGAGAGGACTGCCAGTAC    300

NOTCH2NLC        301 TCGACATCTCATCCATGCTTTGTGTCTCGACCTTGCCTGAATGGCGGCAC    350
                     ||||||||||||||||||||||||||||||||||||||||||||||||||
NOTCH2NLC'       301 TCGACATCTCATCCATGCTTTGTGTCTCGACCTTGCCTGAATGGCGGCAC    350

NOTCH2NLC        351 ATGCCATATGCTCAGCCGGGATACCTATGAGTGCACCTGTCAAGTCGGGT    400
                     ||||||||||||||||||||||||||||||||||||||||||||||||||
NOTCH2NLC'       351 ATGCCATATGCTCAGCCGGGATACCTATGAGTGCACCTGTCAAGTCGGGT    400

NOTCH2NLC        401 TTACAGGTAAGGAGTGCCAATGGACCGATGCCTGCCTGTCTCATCCCTGT    450
                     ||||||||||||||||||||||||||||||||||||||||||||||||||
NOTCH2NLC'       401 TTACAGGTAAGGAGTGCCAATGGACCGATGCCTGCCTGTCTCATCCCTGT    450

NOTCH2NLC        451 GCAAATGGAAGTACCTGTACCACTGTGGCCAACCAGTTCTCCTGCAAATG    500
                     ||||||||||||||||||||||||||||||||||||||||||||||||||
NOTCH2NLC'       451 GCAAATGGAAGTACCTGTACCACTGTGGCCAACCAGTTCTCCTGCAAATG    500

NOTCH2NLC        501 CCTCACAGGCTTCACAGGGCAGAAGTGTGAGACTGATGTCAATGAGTGTG    550
                     ||||||||||||||||||||||||||||||||||||||||||||||||||
NOTCH2NLC'       501 CCTCACAGGCTTCACAGGGCAGAAGTGTGAGACTGATGTCAATGAGTGTG    550

NOTCH2NLC        551 ACATTCCAGGACACTGCCAGCATGGTGGCACCTGCCTCAACCTGCCTGGT    600
                     ||||||||||||||||||||||||||||||||||||||||||||||||||
NOTCH2NLC'       551 ACATTCCAGGACACTGCCAGCATGGTGGCACCTGCCTCAACCTGCCTGGT    600

NOTCH2NLC        601 TCCTACCAGTGCCAGTGCCTTCAGGGCTTCACAGGCCAGTACTGTGACAG    650
                     ||||||||||||||||||||||||||||||||||||||||||||||||||
NOTCH2NLC'       601 TCCTACCAGTGCCAGTGCCTTCAGGGCTTCACAGGCCAGTACTGTGACAG    650

NOTCH2NLC        651 CCTGTATGTGCCCTGTGCACCCTCGCCTTGTGTCAATGGAGGCACCTGTC    700
                     ||||||||||||||||||||||||||||||||||||||||||||||||||
NOTCH2NLC'       651 CCTGTATGTGCCCTGTGCACCCTCGCCTTGTGTCAATGGAGGCACCTGTC    700

NOTCH2NLC        701 GGCAGACTGGTGACTTCACTTTTGAGTGCAACTGCCTTCCAGAAACAGTG    750
                     ||||||||||||||||||||||||||||||||||||||||||||||||||
NOTCH2NLC'       701 GGCAGACTGGTGACTTCACTTTTGAGTGCAACTGCCTTCCAGAAACAGTG    750

NOTCH2NLC        751 AGAAGAGGAACAGAGCTCTGGGAAAGAGACAGGGAAGTCTGGAATGGAAA    800
                     ||||||||||||||||||||||||||||||||||||||||||||||||||
NOTCH2NLC'       751 AGAAGAGGAACAGAGCTCTGGGAAAGAGACAGGGAAGTCTGGAATGGAAA    800

NOTCH2NLC        801 AGAACACGATGAGAATTAG    819
                     |||||||||||||||||||
NOTCH2NLC'       801 AGAACACGATGAGAATTAG    819
```

**Supplementary Fig 29 A visualization of the pairwise alignment between the coding sequences of the NOTCH2NLC and NOTCH2NLC' genes on the maternal haplotype of HG00706.**

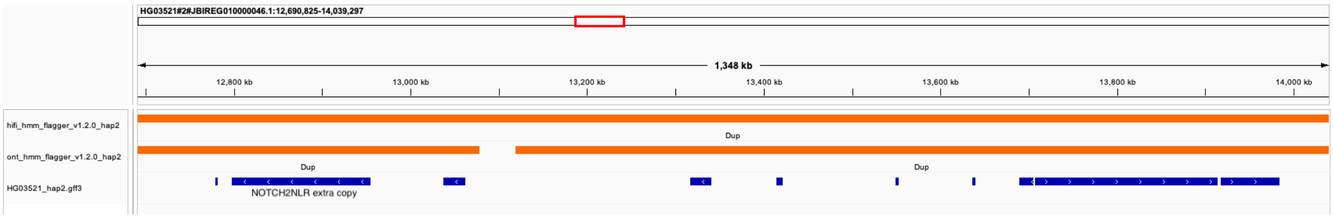


**Supplementary Fig 30 An IGV snapshot of the NOTCH2NL region in HG03521 haplotype 2 with gene annotations and HMM-Flagger annotations displayed, showing the false duplication of NOTCH2NLR. Both ONT- and HiFi-based HMM-Flagger tracks agree on the misassembly.**

| Assembly | Chromosome | Start Position | End Position | Strand | Gene Name | Notes | Flagger Annotations |
|---|---|---|---|---|---|---|---|
| HG03521_hap1 (AFR) | HG03521#1#CM094410.1 | 120566942 | 120756121 | - | NOTCH2 | | Dup |
| | HG03521#1#CM094410.1 | 148217135 | 148300021 | - | NOTCH2NLA | | |
| | HG03521#1#CM094410.1 | 149068582 | 149148347 | + | NOTCH2NLB | | |
| | HG03521#1#CM094410.1 | 151523743 | 151604957 | + | NOTCH2NLC | Missing R | |
| HG03521_hap2 (AFR) | HG03521#2#JBIREG010000050.1 | 120216071 | 120405270 | - | NOTCH2 | | |
| | HG03521#2#JBIREG010000050.1 | 121021651 | 121092627 | + | NOTCH2NLR | | |
| | HG03521#2#JBIREG010000050.1 | 151704690 | 151787552 | - | NOTCH2NLA | | |
| | HG03521#2#JBIREG010000050.1 | 152559591 | 152639290 | + | NOTCH2NLB | | |
| | HG03521#2#JBIREG010000050.1 | 155060590 | 155141808 | + | NOTCH2NLC | | |
| | HG03521#2#JBIREG010000046.1 | 12801125 | 12955216 | - | NOTCH2NLR | Extra R | Dup |

**Supplementary Fig 31 A table showing the presence and location of the NOTCH2 and NOTCH2NL genes on either haplotypes of HG03521. Highlighted in grey are the regions flagged by HMM-Flagger.**

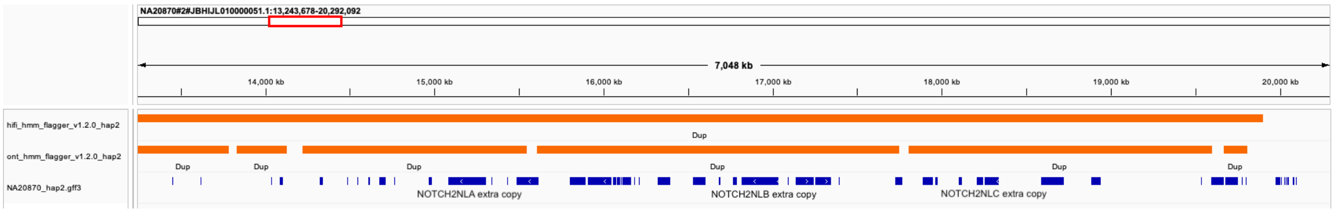


**Supplementary Fig 32 An IGV snapshot of the NOTCH2NLA-C region in NA20870 haplotype 2 with gene annotations and HMM-Flagger annotations displayed, showing the false duplication. Both ONT- and HiFi-based HMM-Flagger tracks agree on the misassembly.**

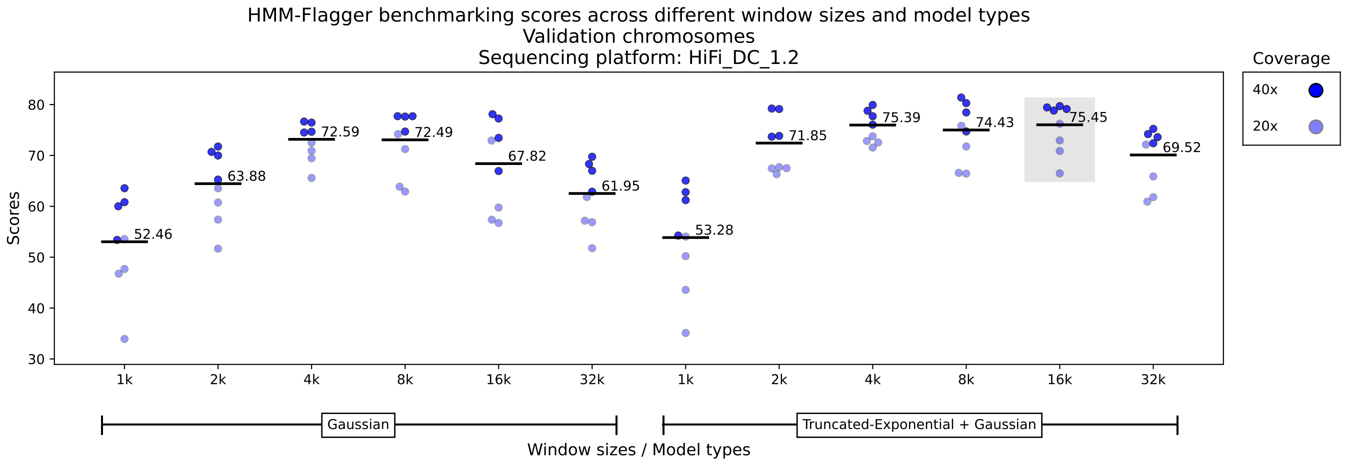


**Supplementary Fig 33 Selecting model type and window size for HiFi reads** The y-axis shows the average of the scores (in percentage) for base-level, overlap-based and auN-based metrics on the validation chromosomes (chr 17 to 22).For each model type and window size 8 points are shown each showing a specific combination of coverage (either 40x or 20x), mapper (winnowmap or minimap2), misassembly rate (either 3.32% or 0.87%). The emission density for the Err state performs best when it is modeled by a truncated exponential distribution. The selected window size is 16k.

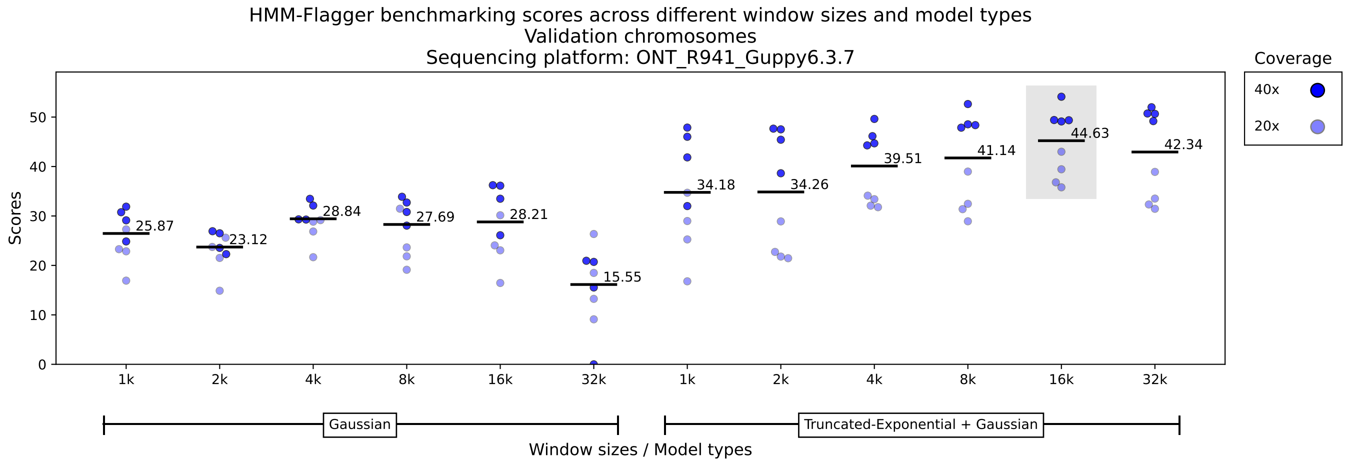


**Supplementary Fig 34 Selecting model type and window size for ONT-R9 reads** The y-axis shows the average of the scores (in percentage) for base-level, overlap-based and auN-based metrics on the validation chromosomes (chr 17 to 22).For each model type and window size 8 points are shown each showing a specific combination of coverage (either 40x or 20x), mapper (winnowmap or minimap2), misassembly rate (either 3.32% or 0.87%). The emission density for the Err state performs best when it is modeled by a truncated exponential distribution. The selected window size is 16k.

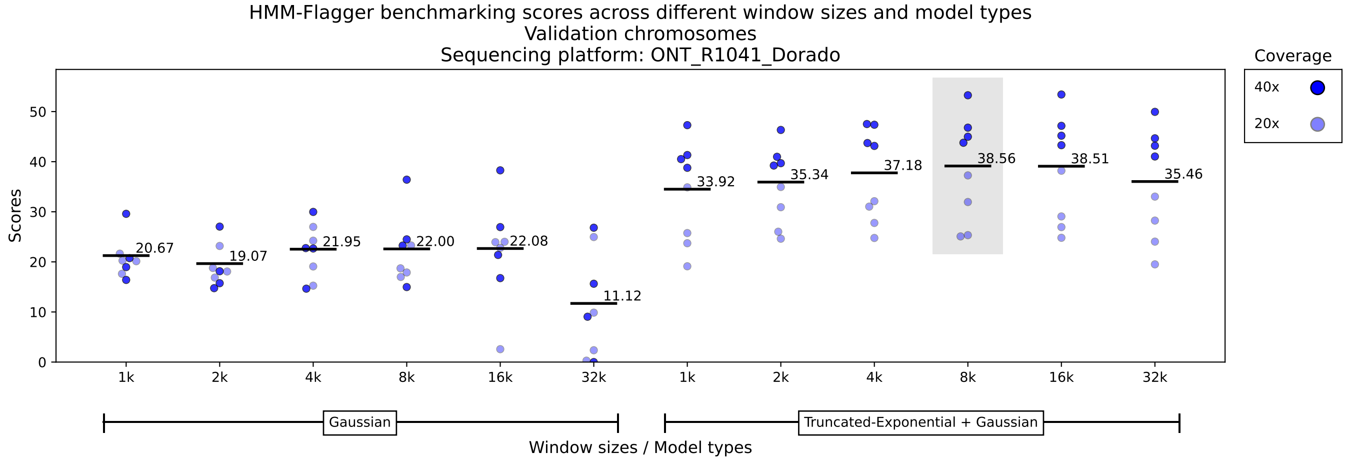


**Supplementary Fig 35 Selecting model type and window size for ONT-R10 reads** The y-axis shows the average of the scores (in percentage) for base-level, overlap-based and auN-based metrics on the validation chromosomes (chr 17 to 22).For each model type and window size 8 points are shown each showing a specific combination of coverage (either 40x or 20x), mapper (winnowmap or minimap2), misassembly rate (either 3.32% or 0.87%). The emission density for the Err state performs best when it is modeled by a truncated exponential distribution. The selected window size is 8k.

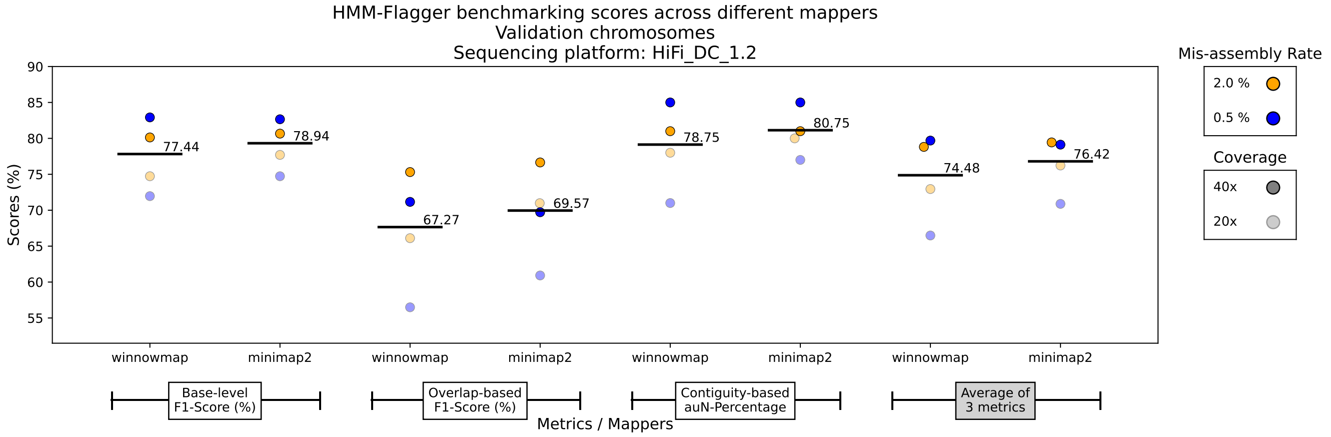


**Supplementary Fig 36 Benchmarking HMM-Flagger across different mappers for HiFi reads** The y-axis shows the scores (in percentage) for base-level, overlap-based, auN-based metrics and their average on the validation chromosomes (chr 17 to 22). For each mapper and metric 4 points are shown each showing a specific combination of coverage (either 40x or 20x) and misassembly rate (either 3.32% or 0.87%). Given HiFi data HMM-Flagger shows comparable performance with minimap2 and winnowmap.

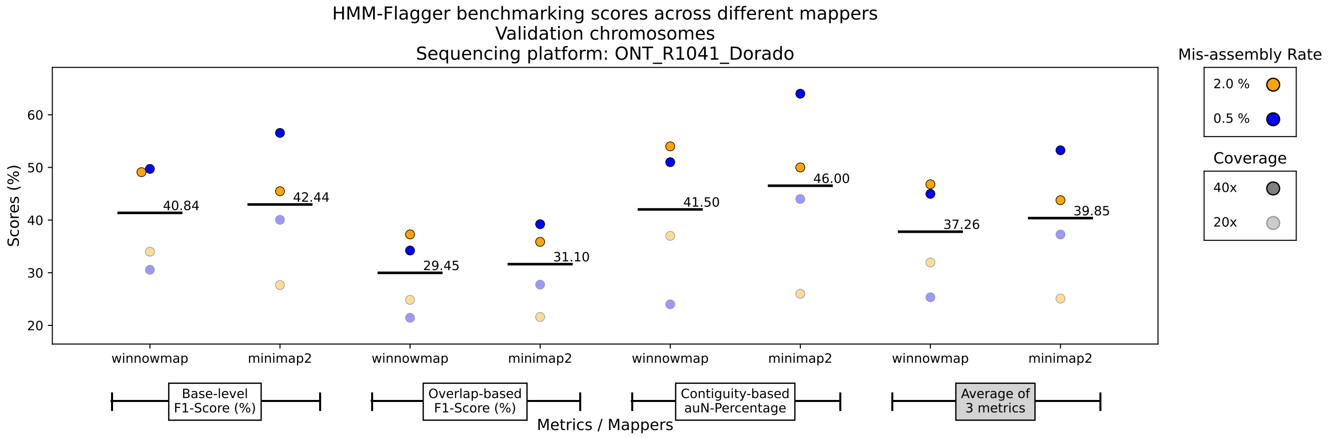


**Supplementary Fig 37 Benchmarking HMM-Flagger across different mappers for ONT-R10 reads** The y-axis shows the scores (in percentage) for base-level, overlap-based, auN-based metrics and their average on the validation chromosomes (chr 17 to 22). For each mapper and metric 4 points are shown each showing a specific combination of coverage (either 40x or 20x) and misassembly rate (either 3.32% or 0.87%). Given ONT-R10 data HMM-Flagger shows comparable performance with minimap2 and winnowmap.

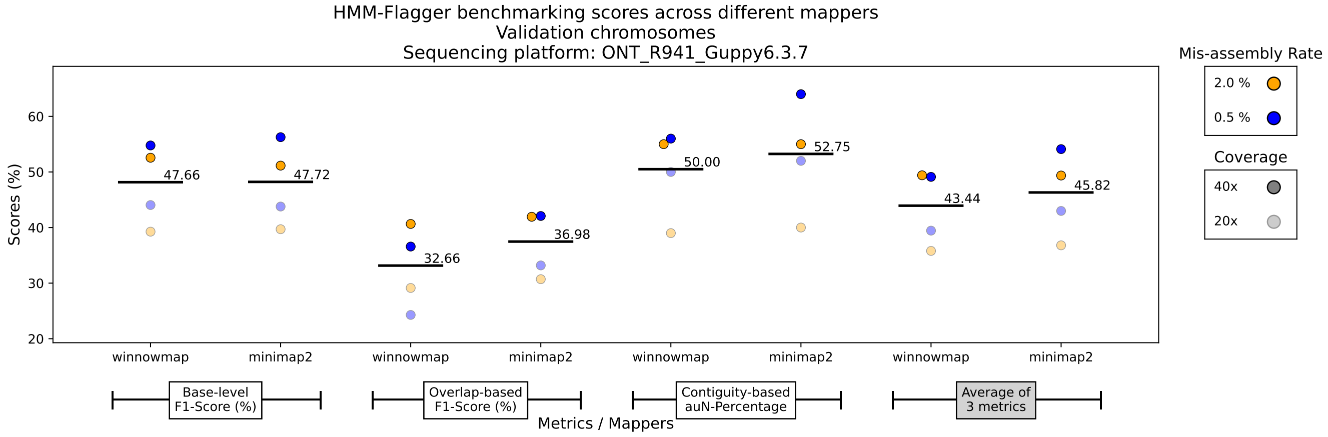


**Supplementary Fig 38 Benchmarking HMM-Flagger across different mappers for ONT-R9 reads** The y-axis shows the scores (in percentage) for base-level, overlap-based, auN-based metrics and their average on the validation chromosomes (chr 17 to 22). For each mapper and metric 4 points are shown each showing a specific combination of coverage (either 40x or 20x) and misassembly rate (either 3.32% or 0.87%). Given ONT-R9 data HMM-Flagger shows comparable performance with minimap2 and winnowmap.

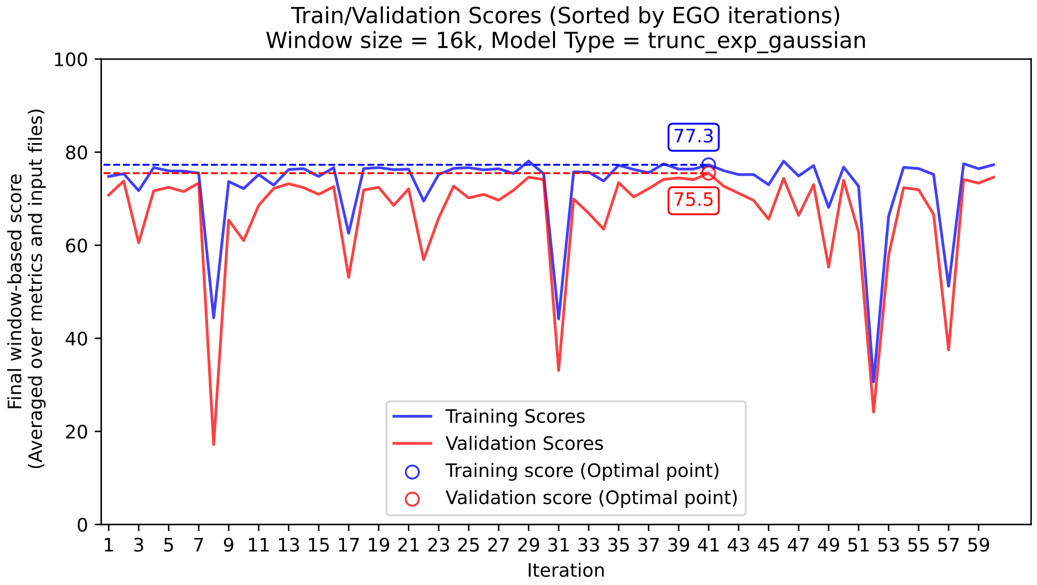


**Supplementary Fig 39 Efficient Global Optimization (EGO) iterations for tuning hyperparameters with HiFi mappings** The EGO algorithm was run for 60 iterations on both training (blue line) and validation (red line) chromosomes. The hyperparameter matrix with the highest score on validation chromosomes was selected (here happened at iteration 41 and shown with circles).

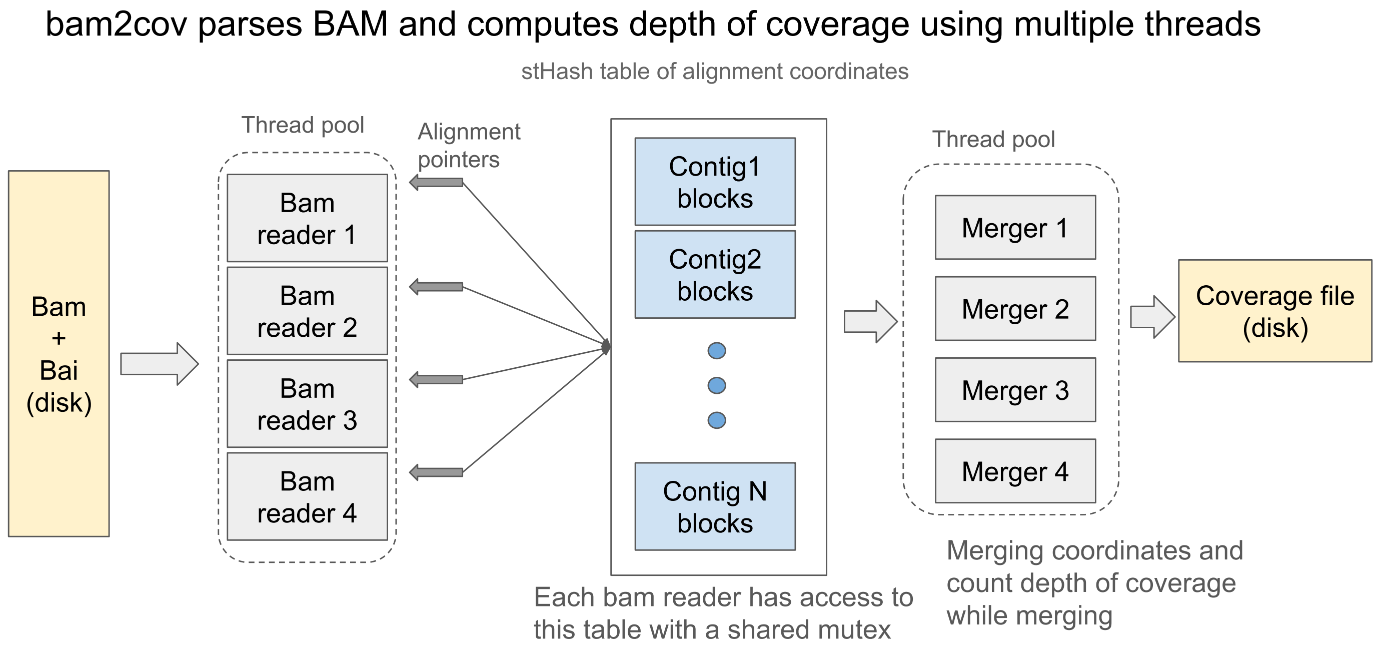


**Supplementary Fig 40 Generating coverage file from BAM:** bam2cov parses BAM and computes depth of coverage using multiple threads. Each BAM reader writes the coordinates of the parsed alignment to the contig table. A multi-threaded merging step merges the coordinates with one contig per thread and counts the depth of coverage while merging.

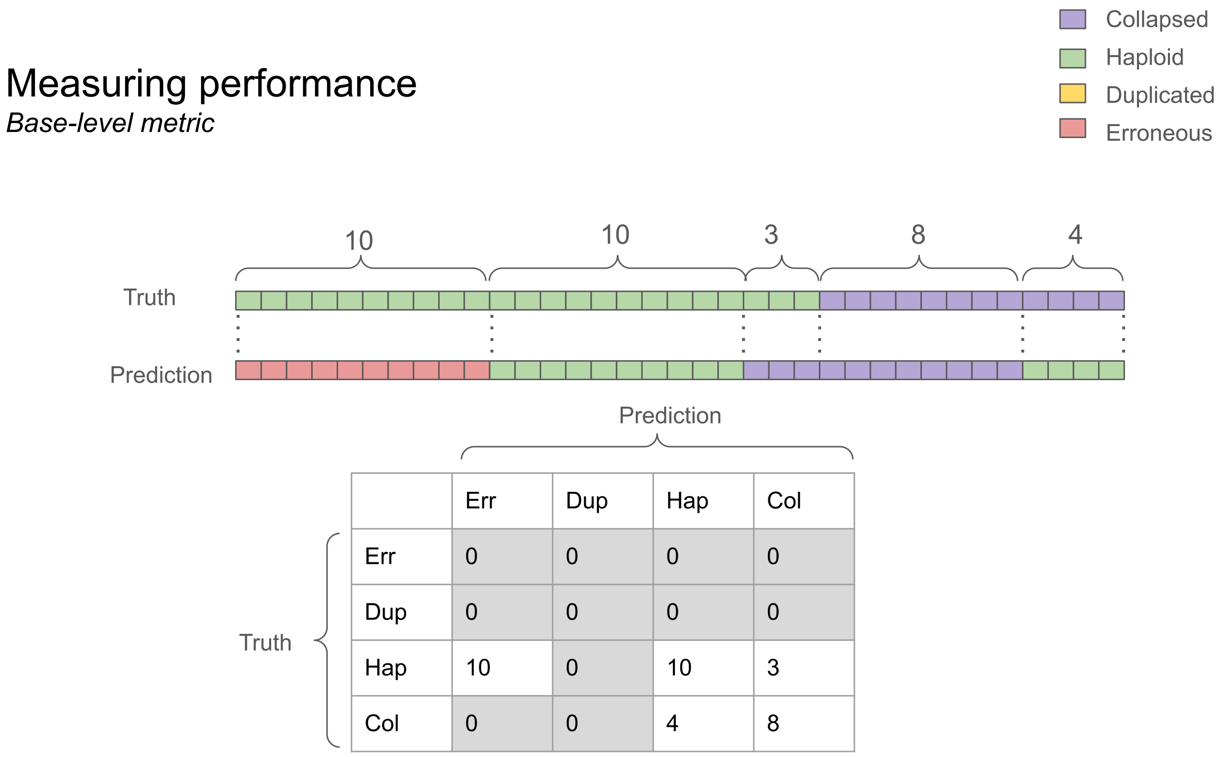


| | Err | Dup | Hap | Col |
|---|---|---|---|---|
| Err | 0 | 0 | 0 | 0 |
| Dup | 0 | 0 | 0 | 0 |
| Hap | 10 | 0 | 10 | 3 |
| Col | 0 | 0 | 4 | 8 |

**Supplementary Fig 41 Measuring performance with base-level metric:** For the base-level metric we count the number of individual bases that fall into each combination of prediction and truth labels and then compute precision, recall and F1-score for each label. For example here we have 10 bases with haploid label misclassified as erroneous.

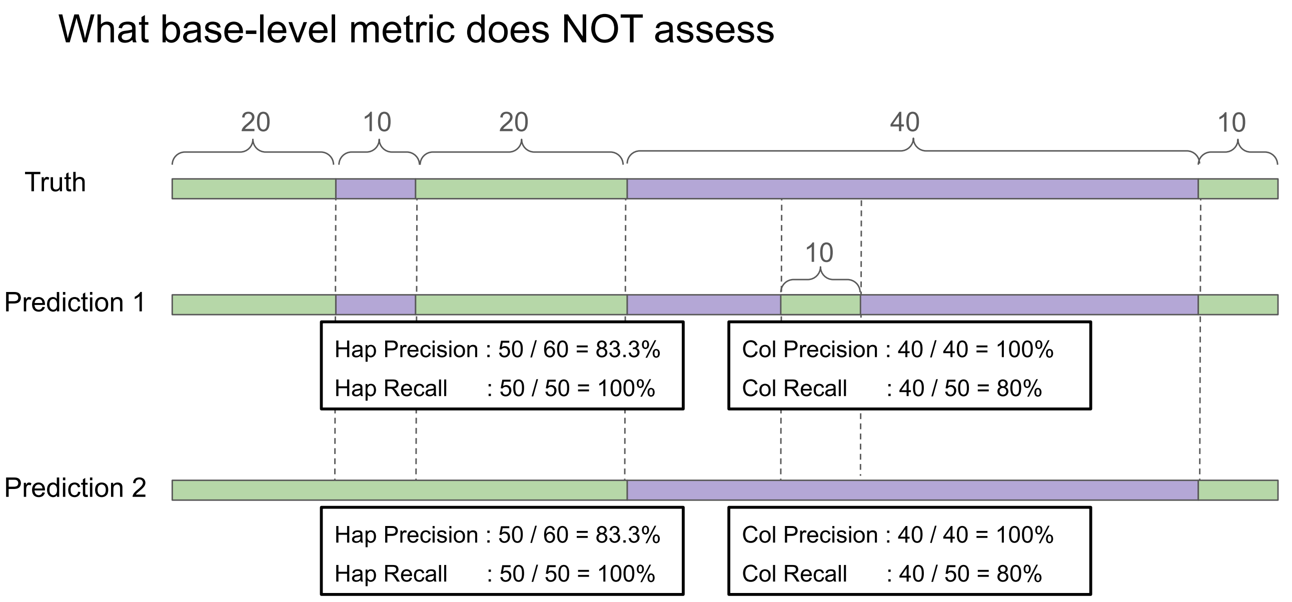


**Supplementary Fig 42 Base-level metric's issue:** Using the base-level metric we have exactly the same values for predictions 1 and 2. However prediction 2 has one disadvantage and that is skipping the whole smaller collapsed event in its entirety.

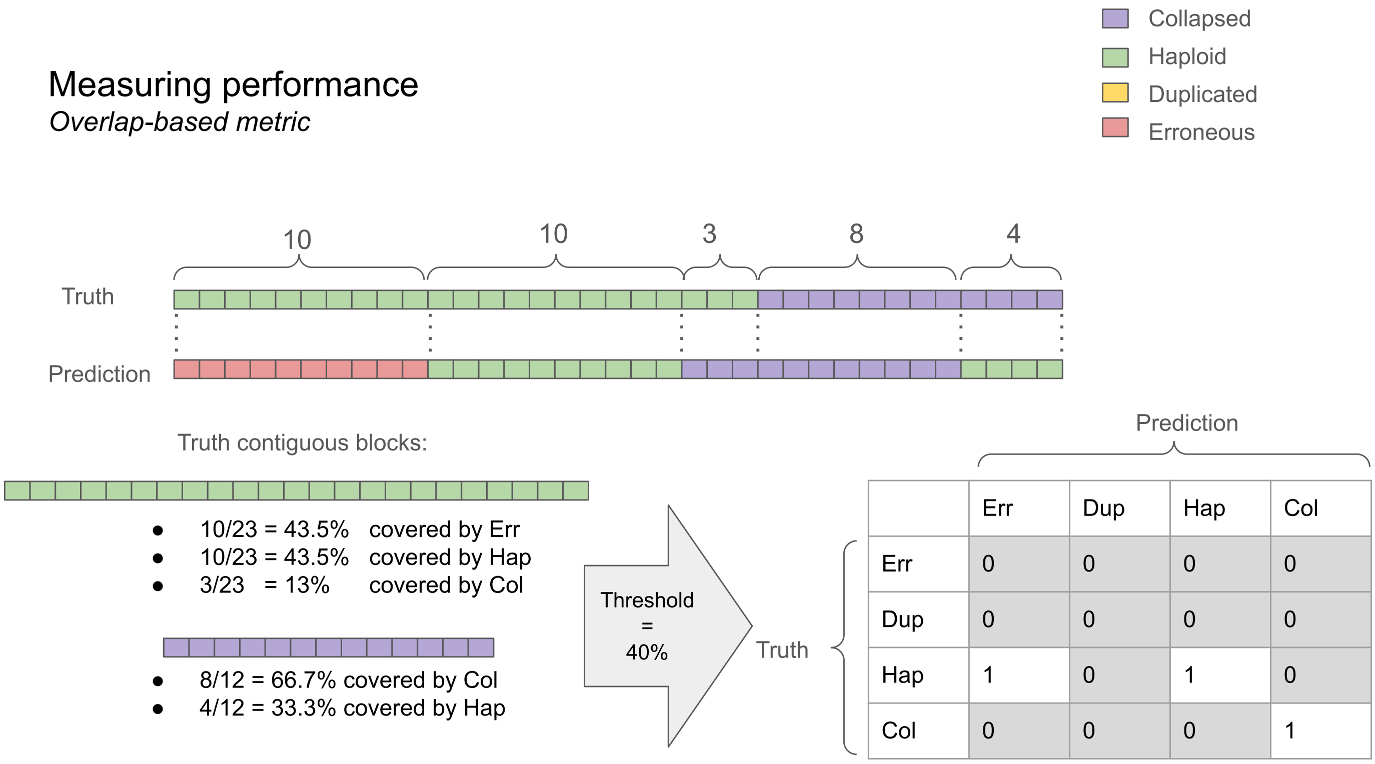


**Supplementary Fig 43 Measuring performance with overlap-based metric:** For the overlap-based metric we take each contiguous block with a single label and measure what percentage of the block is classified as any of the four labels. If the percentage with any combination of labels exceeded the given threshold (here 40%) that counts as a hit for the confusion matrix. This process is done for prediction blocks and truth blocks separately therefore two confusion matrices will be generated in the end.

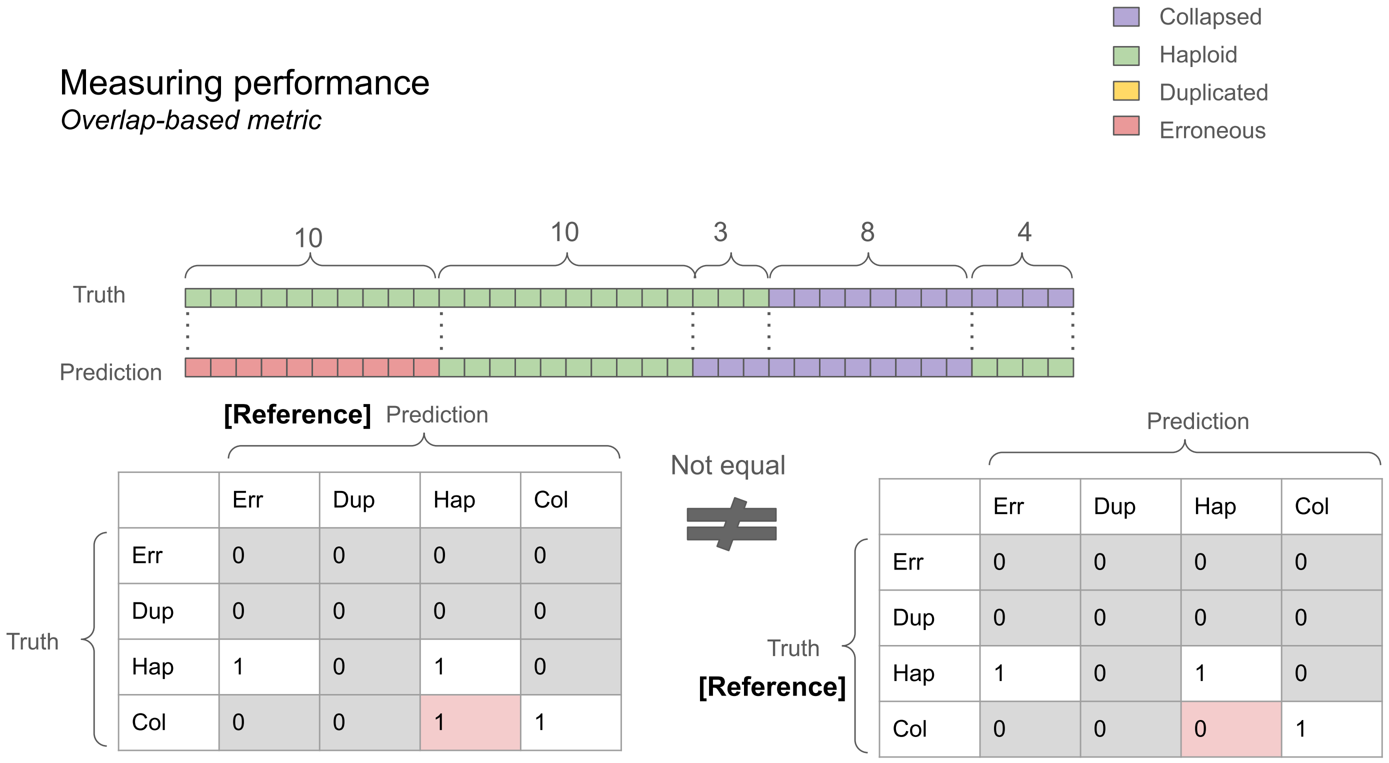


**Supplementary Fig 44 Asymmetry in overlap-based confusion matrix:** For the overlap-based metric, we create two confusion matrices: one using the prediction labels as the reference for computing overlap ratios, and one using the truth labels. These two matrices may not be equal. For example here the block with a predicted collapsed label has a length of 11 out of which 3 bases are classified as haploid (27%) therefore Col-Hap element in the right-handed matrix is zero. However if we use truth labels as the reference the last haploid block (with a length of 4) is 100% misclassified as collapsed , which is reflected in the left-hand matrix.

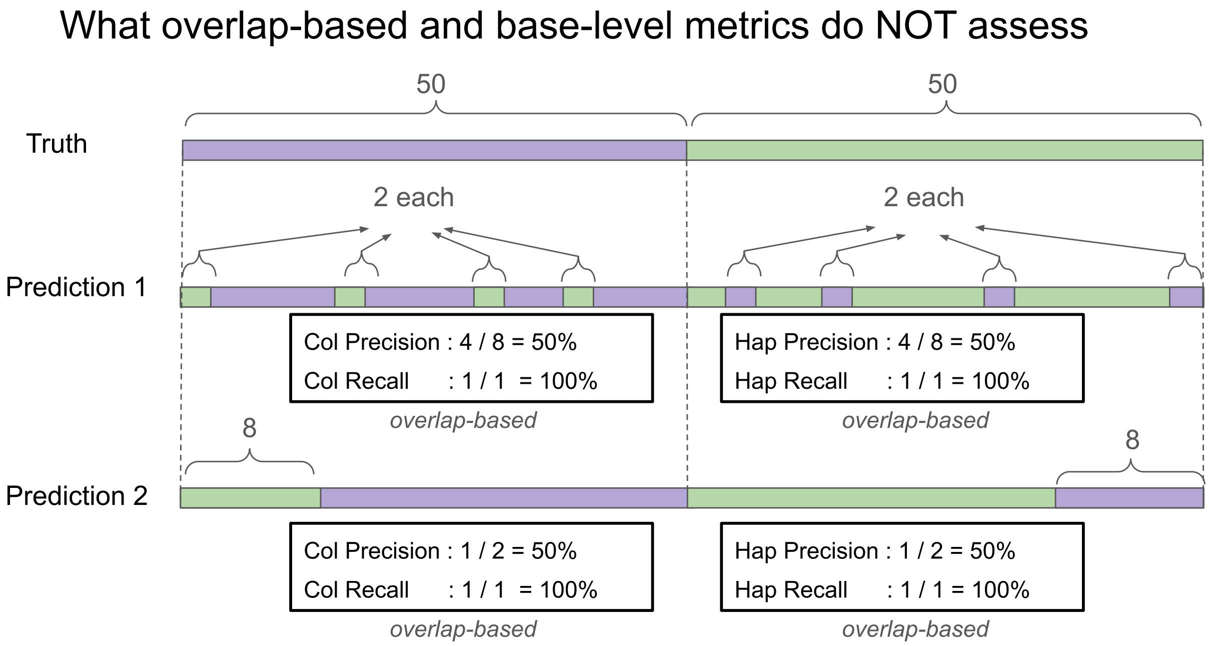


**Supplementary Fig 45 Overlap-based metric's issue:** Both overlap-based and base-level metrics have exactly the same values for these two predictions. However the lengths of true predictions in prediction 2 are closer to the truth events and it has less number of spurious blocks.

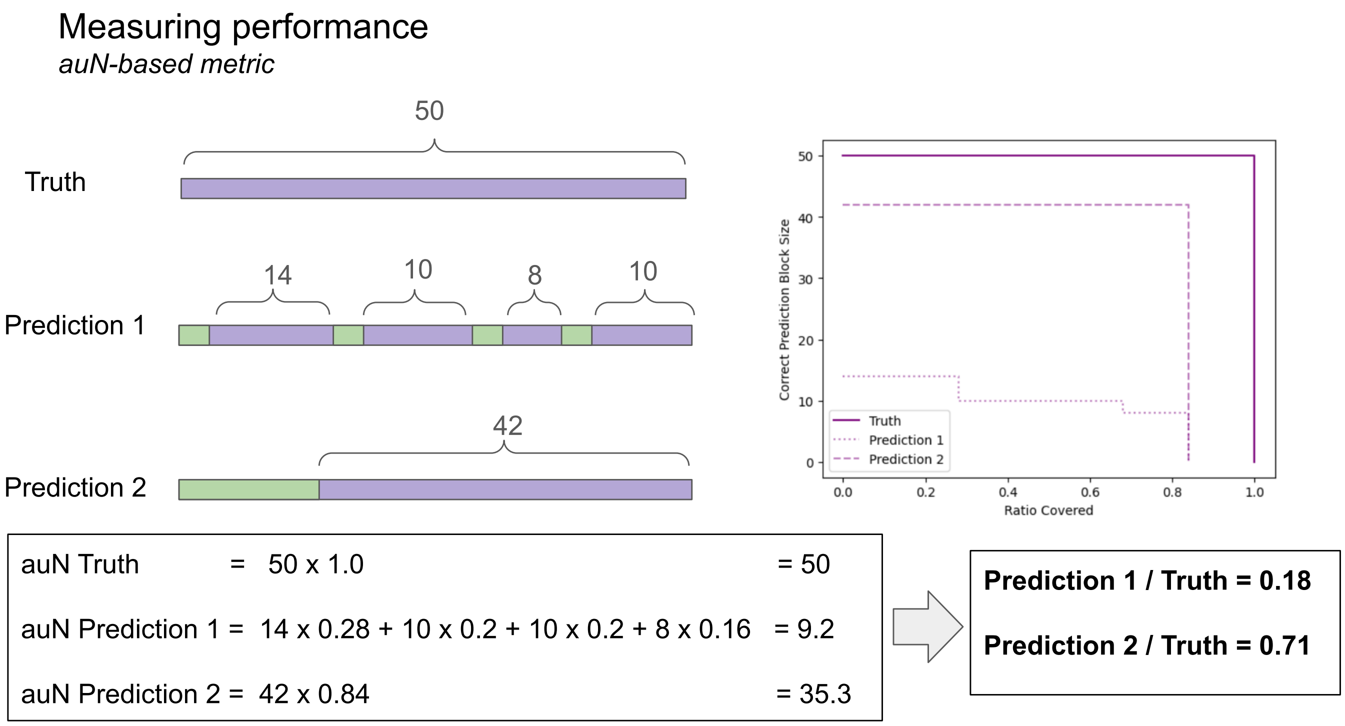


**Supplementary Fig 46 Measuring performance with auN-based metric:** For auN-based metric we compute the area under Nx curves using the lengths of the truth blocks and the lengths of the blocks predicted correctly for each label (as shown in the right-hand panel). Although both predictions have the same number of bases misclassified, the length of the 2nd one better match the truth label, which is reflected in the area under the Nx curve (auN). Each auN value is normalized using the auN of the truth blocks.
